# Connectivity characterization of the mouse basolateral amygdalar complex

**DOI:** 10.1101/807743

**Authors:** Houri Hintiryan, Ian Bowman, David L. Johnson, Laura Korobkova, Muye Zhu, Neda Khanjani, Lin Gou, Lei Gao, Seita Yamashita, Michael S. Bienkowski, Luis Garcia, Nicholas N. Foster, Nora L. Benavidez, Monica Y. Song, Darrick Lo, Kaelan Cotter, Marlene Becerra, Sarvia Aquino, Chunru Cao, Ryan Cabeen, Jim Stanis, Marina Fayzullina, Sarah Ustrell, Tyler Boesen, Zheng-Gang Zhang, Michael S. Fanselow, Peyman Golshani, Joel D. Hahn, Ian R. Wickersham, Giorgio A. Ascoli, Li I. Zhang, Hong-Wei Dong

## Abstract

The basolateral amygdalar complex (BLA) is implicated in behavioral processing ranging from fear acquisition to addiction. Newer methods like optogenetics have enabled the association of circuit-specific functionality to uniquely connected BLA cell types. Thus, a systematic and detailed connectivity profile of BLA projection neurons to inform granular, cell type-specific interrogations is warranted. In this work, we applied computational analysis techniques to the results of our circuit-tracing experiments to create a foundational, comprehensive, multiscale connectivity atlas of the mouse BLA. The analyses identified three domains within the classically defined anterior BLA (BLAa) that house target-specific projection neurons with distinguishable cell body and dendritic morphologies. Further, we identify brain-wide targets of projection neurons located in the three BLAa domains as well as in the posterior BLA (BLAp), ventral BLA (BLAv), lateral (LA), and posterior basomedial (BMAp) nuclei. Projection neurons that provide input to each nucleus are also identifed. Functional characterization of some projection-defined BLA neurons were demonstrated via optogenetic and recording experiments. Hypotheses relating function to connection-defined BLA cell types are proposed.

## INTRODUCTION

The basolateral amygdalar complex (BLA) contains the lateral amygdalar nucleus (LA), the basomedial- and basolateral amygdalar nuclei anterior- (BMAa, BLAa) and posterior (BMAp, BLAp) parts, and the ventral amygdalar nucleus (BLAv) ^1^. Although recognized mostly for its role in fear conditioning and extinction ^2–13^, the BLA is implicated in several behavior-related states and disorders including anxiety ^14^, autism ^15^, addiction ^16–18^, and valence ^19^. Traditional investigative methods of BLA have involved nuclei inactivation via ablative lesioning. However, advances in optogenetic and chemogenetic neurotechnology have enabled more specialized investigations of circuit-specific functional assignments. It is therefore timely to advance understanding of connectivity-specific neurons and test for their participation in behavioral networks. Relatedly, a collective effort is underway to distinguish and characterize the different brain-wide cell types based on connectivity, as well as additional discriminating elements such as molecular and electrophysiological signatures, in hopes of generating a cell type brain atlas.

Recent studies have demonstrated segregation of projection defined cell types in the mouse BLA ^20, 21^. Genetically distinct BLA cell populations with unique functional and connectional phenotypes also have been identified ^22^. For example, a population of BLA projection neurons that target the prelimbic cortical area (PL) are primarily activated during high fear conditions, while neurons that specifically innervate the infralimbic cortical area (ILA) respond predominantly during fear extinction ^23^. These functionally distinct cell groups defined by their projection targets were designated as “fear” and “extinction” neurons, respectively. As a secondary example, activation of pathways from the BLA to the lateral part of the central amygdala nucleus (BLA→CEAl) and to the anterodorsal bed nucleus of stria terminalis (BLA→adBST) result in anxiolytic-like behaviors ^14, 22^, while BLA projections to the ventral hippocampus (BLA→vHPC) have been implicated in anxiogenesis ^24^.

So where are all of these target-specific BLA projection neurons located and, with the supposition that these functionally specific, uniquely connected neurons are intermingled ^23, 25^, where would investigators begin to probe and identify cell type function? One approach would be to begin with a systematic and detailed overview of the connectivity architecture of the BLA that informs higher resolution cell type-specific interrogations. Although connectivity of the BLA has been described in the rat ^26^, it is less examined in the mouse. Further, a whole-brain input/output connectivity profile for the BLA has not been acquired for any species. We applied circuit-level pathway-tracing methods combined with computational techniques to provide a foundational, comprehensive, multiscale connectivity atlas of the mouse BLA complex. Our analyses identified three domains within the classically defined BLAa, each of which houses target-specific projection neurons with distinguishable cell body and dendritic morphologies. Brain-wide projection neurons that provide input to the BLAa domains were also identified. This work further describes the connectional input/output organization of neurons in the BLAp, BLAv, LA, and BMAp. Online resources that allow viewing of the injection cases, the mapped connectivity data (http://www.mouseconnectome.org/amygdalar/) (currently password protected), and an interactive wiring diagram will also be available. Functional hypotheses of connectivity-specific BLA cell types are proposed.

## RESULTS

### Circuit tracing, computational analyses, and visualizations

The Center for Integrative Connectomics (CIC) at USC performed the collection of connectivity data as part of the Mouse Connectome Project (MCP) (www.MouseConnectome.org). Surgically placed anterograde (Phal, BDA, AAV) and retrograde (CTb, FG, AAV retro Cre) pathway-tracers in each nuclei including the BLAa and adjacent BLAp, BLAv, and BMAp, LA (ventromedial region) obtained a comprehensive connectivity characterization of the BLA. Preliminary analysis of anterograde and retrograde pathway-tracing data from injections targeting multiple levels of the medial prefrontal cortex (MPF) and hippocampus (HPF) clearly delineated three BLAa domains with unique connectional profiles. These included the medial (BLA.am), lateral (BLA.al), and caudal (BLA.ac) domains. This finding was supported by the results of additional pathway-tracer injections targeted to the cerebral cortex, thalamus, and striatum that also indicated the existence ofBLA.am, BLA.al, and BLA.ac domains (Figure 1A-C; Supplementary Figures 1A-C & 2).

**Figure 1:**
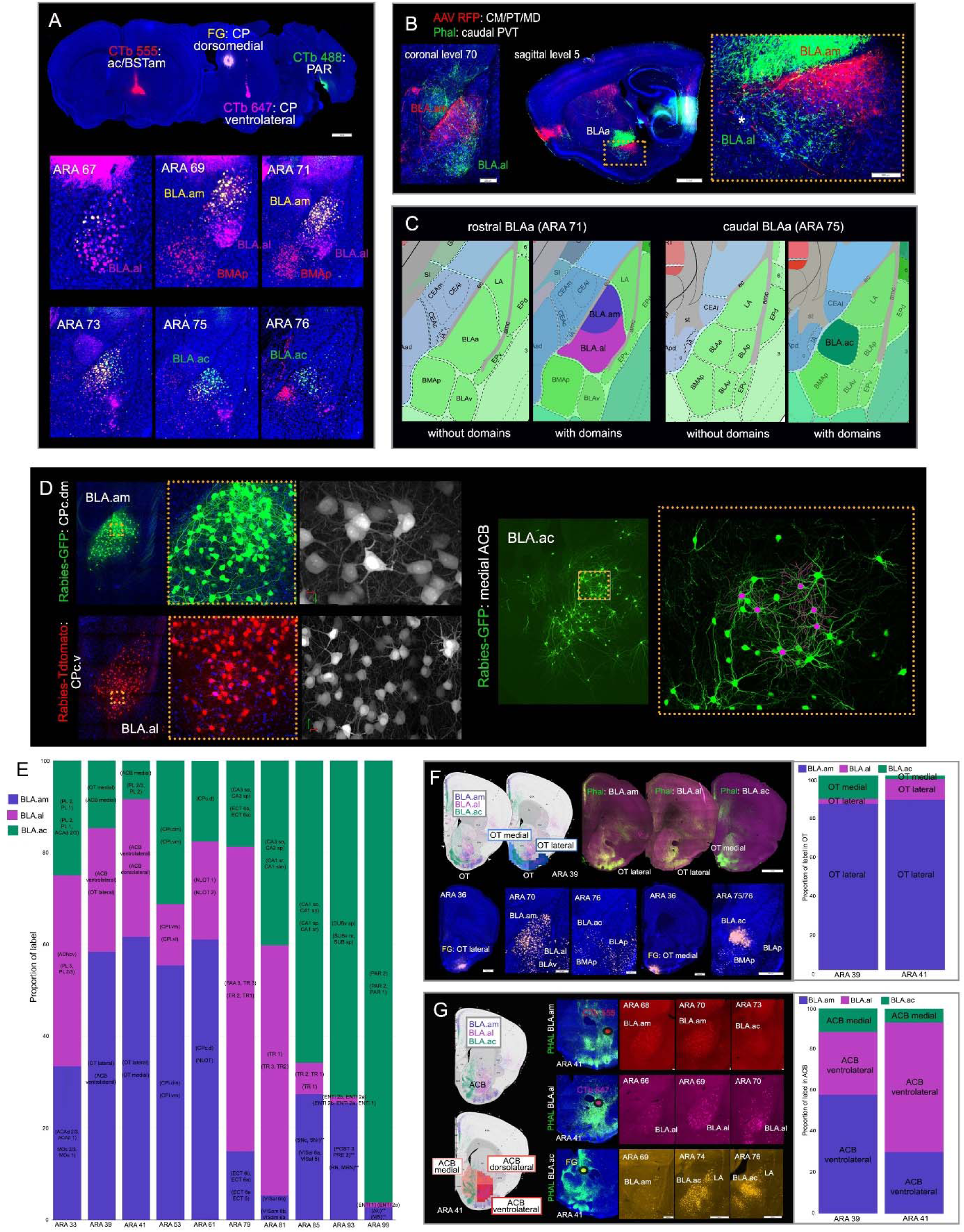
**A.** Four retrograde tracer injections placed in BSTam (CTb 555: red), dorsomedial caudal CP (FG: yellow), ventrolateral caudal CP (CTb 647: pink), and in PAR (CTb 488: green) clearly reveal uniquely connected projection neurons in the three domains of BLAa: BLA.am (FG), BLA.al (CTb 647), and BLA.ac (CTb 488). BMAp projection neurons that target BST are also labeled (CTb 555). Note the segregation between FG (yellow) and CTb 647 (pink) labeled cells in BLA.am and BLA.al, respectively at ARA levels 69 and 71. Also note the absence of CTb 488 (green) labeled cells in BLAa at ARA levels 67-71. *Abbreviations: ac: anterior commissure; BSTam: anteromedial bed nucleus of stria terminalis; CP: caudoputamen; PAR: parasubiculum.* **B.** Anterograde tracers AAV-RFP and Phal injected in different thalamic nuclei distinctly label BLA.am and BLA.al, which is evident in coronal (left) and sagittal (right) views. Inset shows magnified version of boxed BLAa region on sagittal section. * denotes boundary between BLAa and IA at ARA sagittal level 5. *Abbreviations: PT: parataenial thalamic nucleus; CM: central medial thalamic nucleus; MD: mediodorsal thalamic nucleus; PVT: paraventricular thalamic nucleus.* **C.** Atlas representations of rostral (ARA 71: left) and caudal (ARA 75: right) regions of BLAa with and without domains. **D.** Left: G-deleted rabies injected in dorsomedial caudal CP (GFP) and ventrolateral caudal CP (TdTomato) that distinctly label caudoputamen projecting BLAa neurons in BLA.am and BLA.al, respectively. Insets show magnified version of boxed regions. Grayscale images show clear labeling of somas and dendrites. Right: G-deleted rabies injected in medial ACB labels accumbens projecting BLA.ac neurons. Inset shows magnified version of boxed region. Soma and dendrites of neurons selected for reconstruction are shown in pink. *Abbreviations: CPc.dm: dorsomedial part of the caudal caudoputamen; CPc.v: ventral part of the caudal caudoputamen; ACB: nucleus accumbens.* **E.** Bar chart describes proportion of label per each ARA section from single anterograde tracer injections made in BLA.am, BLA.al, and BLA.ac (n=1 each). The ROIs for grids with the strongest projections from each injection displayed in parentheses (e.g. the strongest projections from BLA.am neurons at ARA 47 is to the CP and OT lateral; for BLA.al neurons, the strongest projections are to ACB ventrolateral and CP). Overall, the BLA.am has stronger projections to ARA 47 compared to BLA.al and BLA.ac neurons. Caudal levels (ARA 85-99) receive more projections from BLA.ac than from BLA.am or BLA.al. The strongest projections are mostly to hippocampal structures like CA1, SUB, and PAR. Importantly, each grid unit (square) can include more than one ROI [e.g. (PL2, PL1) or (PL2, PL1, ACAd2/3)]. In this case, ROIs are listed in descending proportion order. ** denotes anterograde projections that were not validated with retrograde tracers. *Abbreviations: PL: prelimbic cortical area; ACAd: dorsal anterior cingulate area; AONpv: anterior olfactory nucleus posteroventral part; MOs: secondary motor area; OT: olfactory tubercle; ACB: nucleus accumbens; CPi.dm: intermediate caudoputamen, dorsomedial part; CPi.vm: intermediate caudoputamen, ventromedial part; CPi.vl: intermediate caudoputamen, ventrolateral part; CPc.d; caudal caudoputamen, dorsal part; NLOT: nucleus of the lateral olfactory tract; ccg; genu of the corpus callosum; TR: postpiriform transition area; VISal: anterolateral visual area; VISam: anteromedial visual area; CA1 sr: CA1 stratum radiatum; CA1 slm: stratum lacunosum-moleculare; CA1 so: CA1 stratum oriens; CA1 sp: CA1 pyramidal layer; ECT: ectorhinal cortical area; CA3 so: CA3 stratum oriens; CA3 sp: CA3 pyramidal layer; SNc: substantia nigra, compact part; SNr: substantia nigra reticular part; SUBv sp: ventral subiculum pyramidal layer; SUBv m: ventral subiculum molecular layer; ENTl: entorhinal cortex lateral part; POST: postsubiculum; PRE: presubiculum; RR: retrorubral area; MRN: midbrain reticular nucleus; MB: midbrain; bic:brachium of the inferior colliculus.* **F.** BLA.am, BLA.al, and BLA.ac projections to olfactory tubercle (OT) at ARA 39, which was split into medial (light blue) and lateral (dark blue) regions. BLA.am and BLA.al neurons target OT lateral, while those in BLA.ac target OT medial. Bottom panels show validation of these projection patterns via retrograde FG injections. FG injected into OT lateral strongly labels BLA.am and BLA.al cells, but also BLAv cells, while a FG injection in OT medial strongly labels BLA.ac neurons and more lightly BLAp and BMAp neurons. The bar chart to the right quantifies and visualizes the density of BLAa→OT connections at ARA levels 39 and 41. **G.** Left panels: BLA.am, BLA.al, and BLA.ac projections to nucleus accumbens (ACB) at ARA 41, which was split into medial (light orange), dorsolateral (orange), and ventrolateral (dark orange) regions. BLA.am neurons target mostly ACB ventrolateral, BLA.al neurons target ACB dorsolateral, while those in BLA.ac innervate ACB medial. To the right, these BLAa→ACB projections are shown with Phal (green) injections made into BLA.am, BLA.al, and BLA.ac. Ellipses denote location of retrograde tracer injections, which were used to validate each BLAa→ACB connection. A CTb 555 injection in ACB ventrolateral regions labels BLA.am neurons (BLA.am→ACB ventrolateral), a CTb 647 injection in ACB dorsolateral regions labels BLA.al neurons (BLA.al→ACB dorsolateral), and a FG injection in medial ACB labels mostly BLA.ac neurons (BLA.ac→ACB medial). The bar chart to the right shows quantified and visualized density of BLAa→ACB connections at ARA levels 39 and 41. Note that although a slight dorsolateral and ventrolateral distinction in projections from the BLA.am and BLA.al are noticeable and validated with retrograde tracers, the bar graph shows densest input to ACB from both BLA.am and BLA.al is to ACB ventrolateral.

To determine BLAa domain characteristics and boundaries, we first identified representative cases demonstrating selective labeling in each domain. For each case, we histologically prepared all consecutive sections across the BLA and imaged the sections with an Olympus VS120 virtual slide microscope (Supplementary Figure 2). Next, we processed sections through our in-house image processing software Connection Lens. Briefly, each section was matched and warped to their corresponding atlas level of the Allen Reference Atlas (ARA) and the labeling was segmented (Figure 2A). The labeling from each case was then manually mapped and aggregated atop the ARA, and approximate boundaries were delineated for the BLA.am, BLA.al, and BLA.ac domains (Figure 2B: Supplementary Figure 3A). Boundaries between BLA.am and BLA.al also were assessed by examination of BLA cytoarchitecture (Nissl stain) (Supplementary Figure 3B). For validation purposes, the same data was used as a training set for a machine learning algorithm to compute the boundaries of the three BLAa domains. These boundary demarcations highly corroborated those of the manual analysis, resulting in an average agreement of 92% Supplementary Figure 3A). Utilizing these boundary approximations, anterograde and retrograde pathway-tracer injections were made to involve mostly BLA.am, BLA.al, or BLA.ac (Supplementary Figure 4A). G-deleted rabies expressing either red (TdTomato) ^27^ or green (GFP) ^28^ fluorescent protein was injected into select targets of each BLAa domain to reveal the morphological details of the domain-specific projection neurons (Figure 1D). Anterograde and retrograde tracer injections were also made in BLAp, BLAv, BMAp, and LA (ventromedial part).

**Figure 2:**
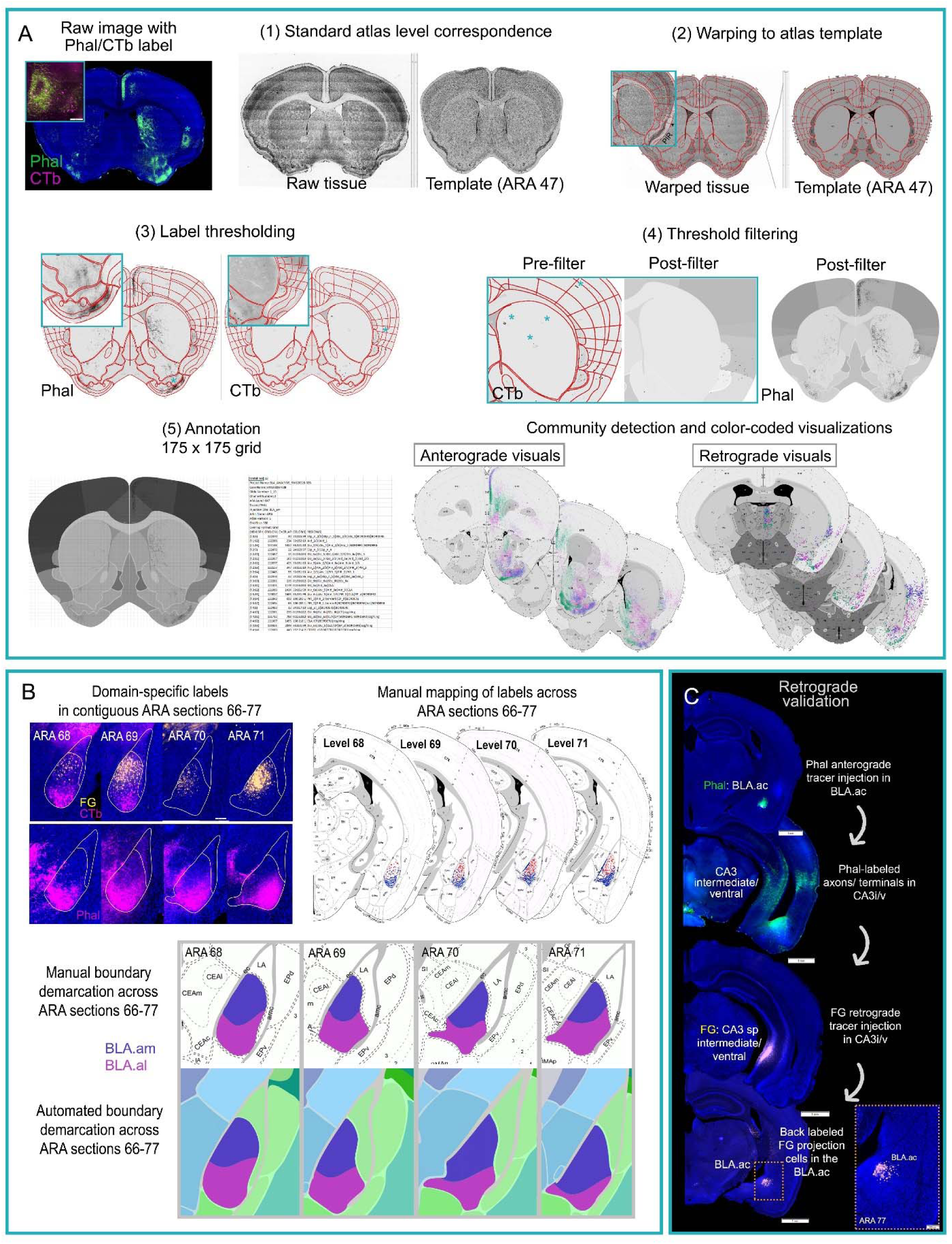
**A.** Our 2D post-image processing pipeline. Whole brain images with fluorescent tracers (e.g., Phal, green fibers and CTb, pink cells) are acquired via an Olympus VS120 slide scanner under 10x magnification. Next, images are imported into an in house software, Connection Lens, for (1) atlas correspondence to match each data section to its corresponding ARA atlas template, (2) to warp data sections to atlas templates, (3) to threshold labeling, (4) filter artifacts, and (5) annotate the labels. Boxed region shows magnified section to illustrate accuracy of registration in (2) [esp. in e.g., layers of the piriform cortex (PIR)]. Insets display magnified regions demonstrating accuracy of segmented Phal fibers and CTb cells in (3). Asterisks highlight artifacts that are filtered in (4) to reduce false positives. Annotation was performed at the grid level (175 × 175 pixels) to capture topographic labels within ROIs that otherwise would go undetected [e.g., olfactory tubercle (OT), nucleus accumbens (ACB)]. Overlap processing in (5) results in an Excel file with annotated values: pixel density for anterograde tracers and cell counts for retrograde tracers. Next, a modularity maximization (community detection) algorithm is applied to the annotated data to assign labels to an injection site based on label density. Community assignments are color-coded by injection site and visualized for 32 ARA sections, both anterograde and retrograde visuals. **B.** The workflow for delineating the boundaries for BLAa domains is presented (see also Supplementary Figures 2-3). First, cases with distinct labels within each domain were selected (n=7) and contiguous sections through the BLAa from ARA level 66 to ARA77 were collected. Sections were imaged and registered, and labels for each case were manually mapped onto a standard atlas. Each case was rendered into individual layers so that mapped data across cases could be aggregated. Borders were manually drawn guided by the mapped labels, but also by the Nissl stains for each case (Supplementary Figure 3). The same data was used as a training set for a machine learning algorithm, which produced similar delineations of BLAa domains. **C.** Retrograde validation of anterograde labeling. Phal injected in BLA.ac labels the CA3 intermediate/ventral CA3i/v) (BLA.ac→CA3i/v). A FG injection in CA3i/v sp layer back-labels projection cells in BLA.ac confirming the BLA.ac→CA3i/v connection. Inset is magnification of boxed region on the whole brain section clearly showing the confined CA3 projecting FG-labeled cells in BLA.ac.

To attain the individual brain-wide connectional patterns of each BLA region, one series of a 1-in-4 series of sections was selected, histologically prepared, imaged, and then processed through Connection Lens. The sections were assigned and warped to a preselected standard set of 32 ARA atlas levels ranging from levels 25-103. Threshold parameters were individually adjusted for each case and pathway-tracer and conspicuous artifacts in the threshold output were corrected (Figure 2A). Next, data at each of the 32 ARA levels was annotated at 175 × 175 pixel grid resolution. Subsequent Louvain community detection analyses assigned each injection site to a community based on labeling patterns with regard to individual grid cells (Figure 2A). The extent and complexity of the topographic projections from BLA to single ROIs (e.g., nucleus accumbens, caudoputamen, olfactory tubercle) necessitated the identification of more granular connections via grid analysis application of the community detection algorithm (Figure 1F-G). In further processing applied to both anterograde and retrograde labeling, the Louvain community assignments were color-coded by injection site and visualized for all 32 ARA sections (Figure 2A). The color-coded output of all community analyses will be available online (http://www.mouseconnectome.org/amygdalar/) (currently password protected) (Supplementary Figure 1D). These injection color codes are used consistently throughout this work. Community analysis was isolated across separate categories of injections. For example, data from the BLA.am, BLA.al, and BLA.ac were aggregated, analyzed, and visualized to identify regions of unique input and output. Given the close proximity of the BLAp to the BLA.al, a separate analysis comparing labeling from the two nuclei were conducted to validate their disparate connectivity profiles. The BLAv, BMAp, and LA were similarly grouped and analyzed. Intra-amygdalar connections were excluded from the analysis given the challenge to accurately distinguish labels from background in nuclei located close to injection sites. Brain-wide connections of all BLA projection neurons are presented in an interactive wiring diagram (link will be provided). In addition, data for all BLA nuclei were combined and subjected to community detection analysis whose output was visualized in a matrix. Within the matrix, injection sites grouped based on the strength of their input and output to ROIs are presented along a diagonal (Figure 3H-I; Figure 4F-G) (http://www.mouseconnectome.org/mcp/docs/BLA/brain_wide_matrix.pdf) (currently password protected).

**Figure 3:**
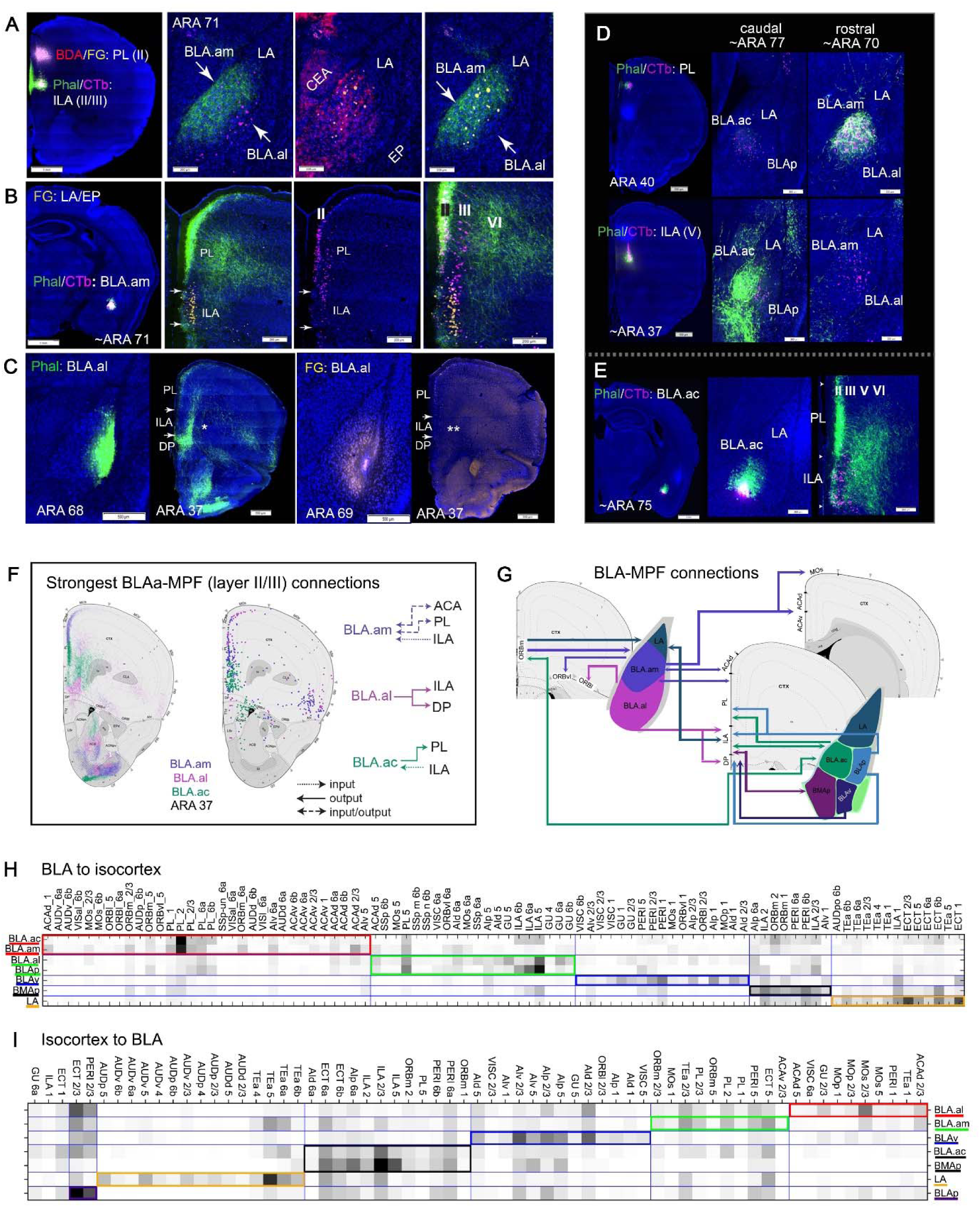
**A.** Double co-injection of BDA/FG in PL(II) and Phal/CTb in ILA(II/III) identified the medial (BLA.am) and lateral (BLA.al) domains of BLAa. Phal and BDA fibers and FG cells are present in BLA.am, while CTb cells are present in BLA.al. These suggest PL→BLA.am, ILA→BLA.am, and BLA.al→ILA connections. **B.** The first two connections were validated with a Phal/CTb injection in BLA.am, which shows strong fiber labeling in PL, especially layer II/III [BLA.am→PL(II/III)] and CTb labeling in PL(II) [PL(II)→BLA.am] and ILA(III) [ILA(III)→BLA.am]. The LA FG injection labels and clearly delineates the ILA. **C.** The BLA.al→ILA connection was validated with a Phal injection in BLA.al, which showed strong fiber labels in ILA and DP. A FG injection in the BLA.al confirms the absence of an ILA→BLA.al connection. ** denotes the ILA and lack of FG labeling. **D.** The BLA.ac also has a unique connection profile with medial prefrontal cortical areas (MPF). A Phal/CTb injection in PL shows CTb labeled BLA.ac projection cells, but very sparse Phal fiber labels in the BLA.ac, suggesting a BLA.ac→PL connection. Phal fibers from PL localize mostly in the more rostral BLA.am. A Phal/CTb injection in ILA(V) shows very strong fiber projections in BLA.ac suggesting a very strong ILA→BLA.ac connection. Only a few CTb cells are present in BLA.ac suggesting a weak BLA.ac→ILA connection. **E.** A Phal/CTb injection in BLA.ac validates these connections, showing strong projections to PL, especially layer II/III [BLA.ac→PL (II/III)] and CTb labeled cells in layers II-V of ILA (ILA→BLA.ac). *Abbreviations: PL: prelimbic cortical area; ILA: infralimbic cortical area; DP: dorsal peduncular area. CEA: central nucleus of the amygdala; EP: endopiriform cortical area.* **F.** Different BLAa domains share unique input/output connections with MPF, especially layers II/III, which is summarized in this schematic. Note (1) the reciprocal connections between the BLA.am and anterior cingulate cortex (ACA) and PL, (2) that the BLA.al is the only domain to project to ILA, and (3) the unique BLA.ac connection in which it heavily projects to PL, but receives strong input from ILA. **G.** Schematic summarizing the connections of all BLA nuclei with MPF areas. Connection are all color-coded. **H.** Community detection confined to BLA projections to isocortical areas was run and visualized in a matrix. The matrix was reordered such that injection sites grouped with their strongest projections are arranged along the diagonal. The grouped injection sites and their connections are boxed in different colors. The weighting of each connection is indicated by a color gradient from black (very strong) to white (very weak). Also matrix analysis was not predicated on grid based annotated data, but was ROI based instead. Based on connections to isocortical areas, BLA.am and BLA.ac were grouped together with projections to PL(I-III,VI), ORBm, and ACAd among the strongest. The BLA.al and BLAp were grouped with projections to ILA, GU, and PL(V) being the strongest. The BLAv shows strongest projections to AI, GU, and PERI, the BMAp to ILA, ORBm, PERI, and the LA to ECT, TEa, and ILA. **I.** Community detection confined to isocortical projections to BLA was run and visualized in a matrix, which shows the BLA.ac and BMAp grouped with their strongest input from ILA(II/III). The BLA.al, BLA.am, BLAv, LA, and BLAp were individually grouped. Note the strong inputs to BLAv from agranular insular areas (AI) and to LA from auditory cortices (AUD). *Abbreviations: PL: prelimbic cortical area; ILA: infralimbic cortical area; ORBm: orbital cortex, medial part; ACAd: anterior cingulate cortical area, dorsal part; GU: gustatory cortical area; PERI: perirhinal cortical area; ECT: ectorhinal cortical area; TEa; temporal association areas*.

**Figure 4:**
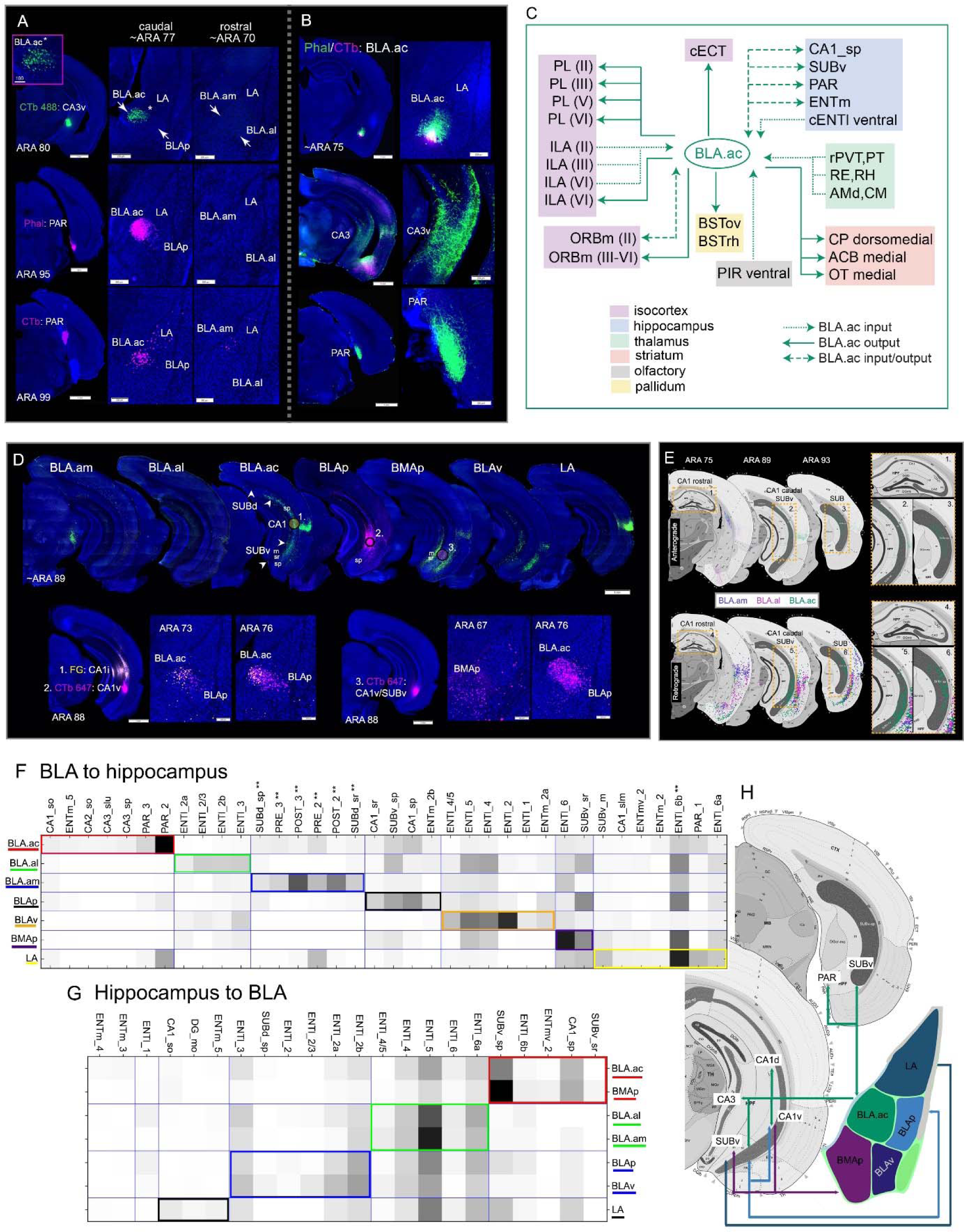
**A.** BLA.ac connections with hippocampal regions. A retrograde CTb 488 (green) tracer injected in ventral CA3 (CA3v) selectively labels BLA.ac projection neurons (BLA.ac→CA3v). Note the absence of labeled cells in rostral BLA.am. Boxed region is a magnification of CA3 projecting CTb labeled cells denoted with an asterisk. A Phal and CTb injection in parasubiculum (PAR) shows the BLA.ac→PAR and PAR→BLA.ac connection. **B.** These connections were validated with a Phal/CTb injection in BLA.ac that shows strong Phal fiber labels in CA3 and PAR. **C.** Summarized brain-wide connections of BLA.ac projection neurons. For full abbreviation list see Table 1. **D.** Top panels show projections from BLA.am, BLA.al, BLA.ac, BLAp, BMAp, BLAv, and LA neurons to hippocampal regions. The BLA.ac, BLAp, and BMAp show strongest projections, with BLA.ac projecting to sp layers of CA1 and SUBv (BLA.ac→CA1sp/SUBv sp) and BLAp to sp layers of CA1v and SUBv (BLAp→CA1v sp/SUBv sp), and BMAp to sr and m layers of CA1v and SUBv (BMAp→CA1v sr/SUBv m/sr). Bottom panels show validation of these connections with FG and CTb marked 1-3 on top panels. Injections 1 and 2 show FG and CTb injections in intermediate (CA1i) and ventral (CA1v) CA1, respectively. Both injections back-label projection neurons in BLA.ac, while only the injection in CA1v labels BLAp neurons. Injection 3 is a CTb injection in SUBv, which labels BLA.ac, BLAp, and BMAp neurons. **E.** Top panels show BLA.ac projections to anterior CA1 (ARA 75) and more caudal CA1 and SUB (ARA 89/93). Boxed regions are numbered and magnified to the right. Bottom panels show projections from CA1 and SUB back to BLA.ac (CA1v/SUBv→BLA.ac). These connections were validated with a Phal injection (see Figure 6N). **F.** Community detection confined to BLA projections to hippocampal areas was run and visualized in a matrix. The matrix was reordered such that grouped injection sites and their strongest projections are arranged along the diagonal. The grouped injection sites and their connections are boxed in different colors. The weighting of each connection is indicated by a color gradient from black (very strong) to white (very weak). Note the (1) strong projections from BLA.ac to PAR and CA3, (2) from BLAp to CA1 sp and SUBv sp, (3) from BLAv to ENTl (layers II and V), and (4) from BMAp to SUBv sr. ** indicates strong connections that were not validated. **G.** Community detection confined to hippocampal projections to BLA was run and visualized in a matrix. Note how the BLA.ac and BMAp are grouped together with strong inputs from CA1 and SUBv as are the BLA.am and BLA.al with strong input from ENTl. Also matrix analysis was not predicated on grid based annotated data, but was ROI based instead. **H.** Schematic summarizing the connections of all BLA nuclei with hippocampal areas. Connection are all color-coded. *Abbreviations: CA1 sr: CA1 stratum radiatum; CA1 sp: CA1 pyramidal layer; SUBv sr: ventral subiculum stratum radiatum; SUBv sp: ventral subiculum pyramidal layer; SUBv m: ventral subiculum molecular layer; ENTl: entorhinal cortical area, lateral part*.

**Table 1.**
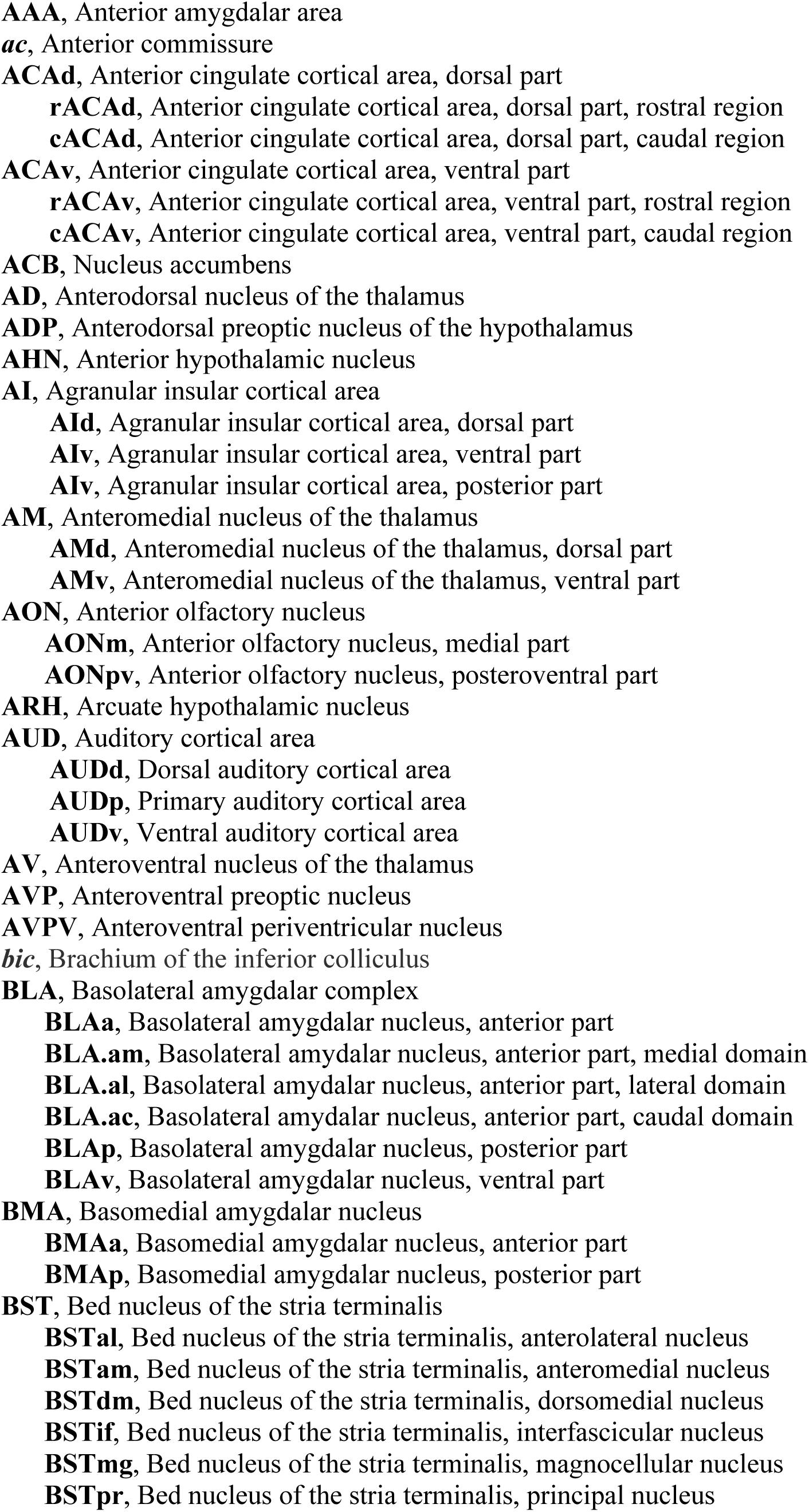
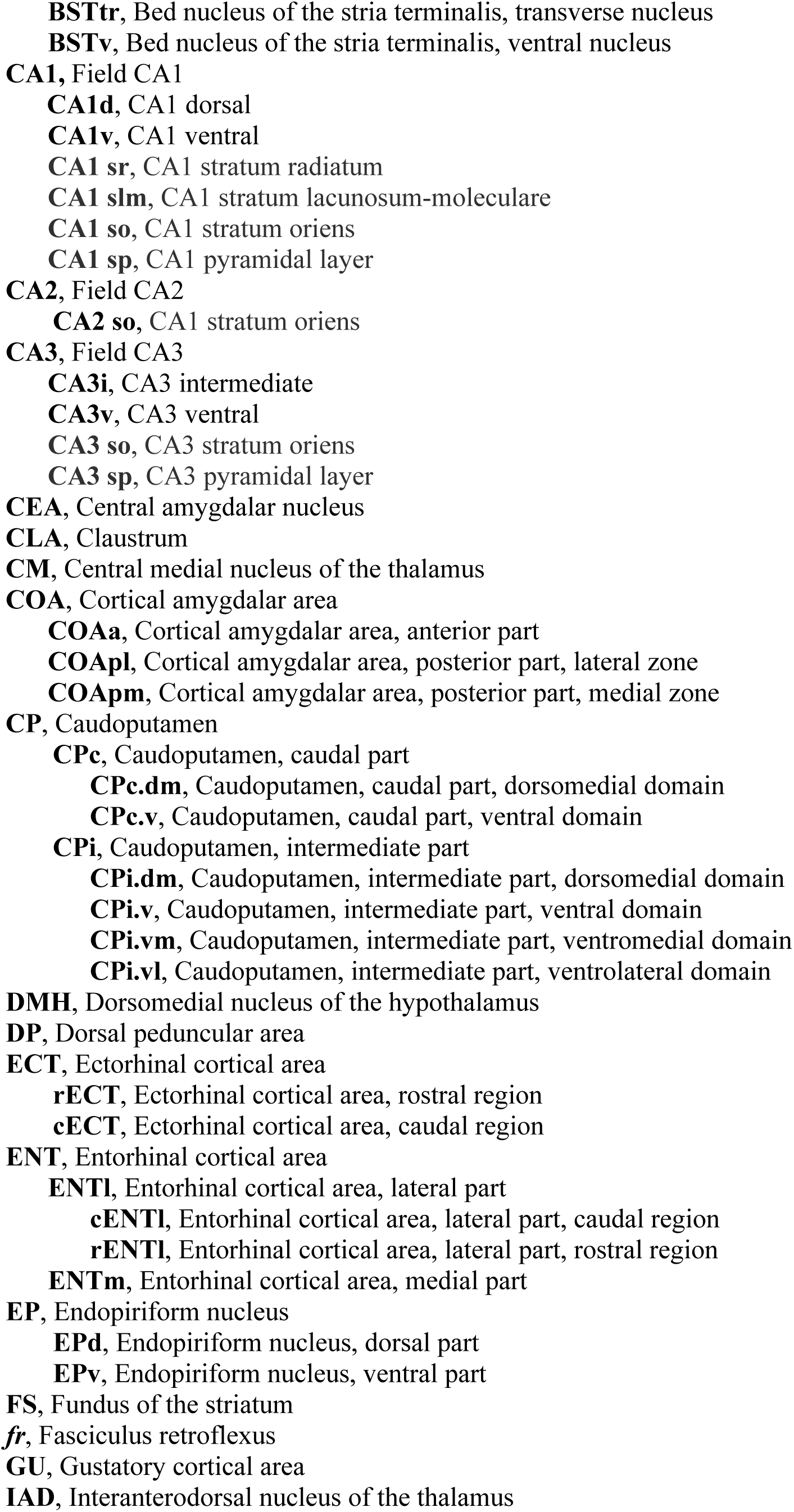
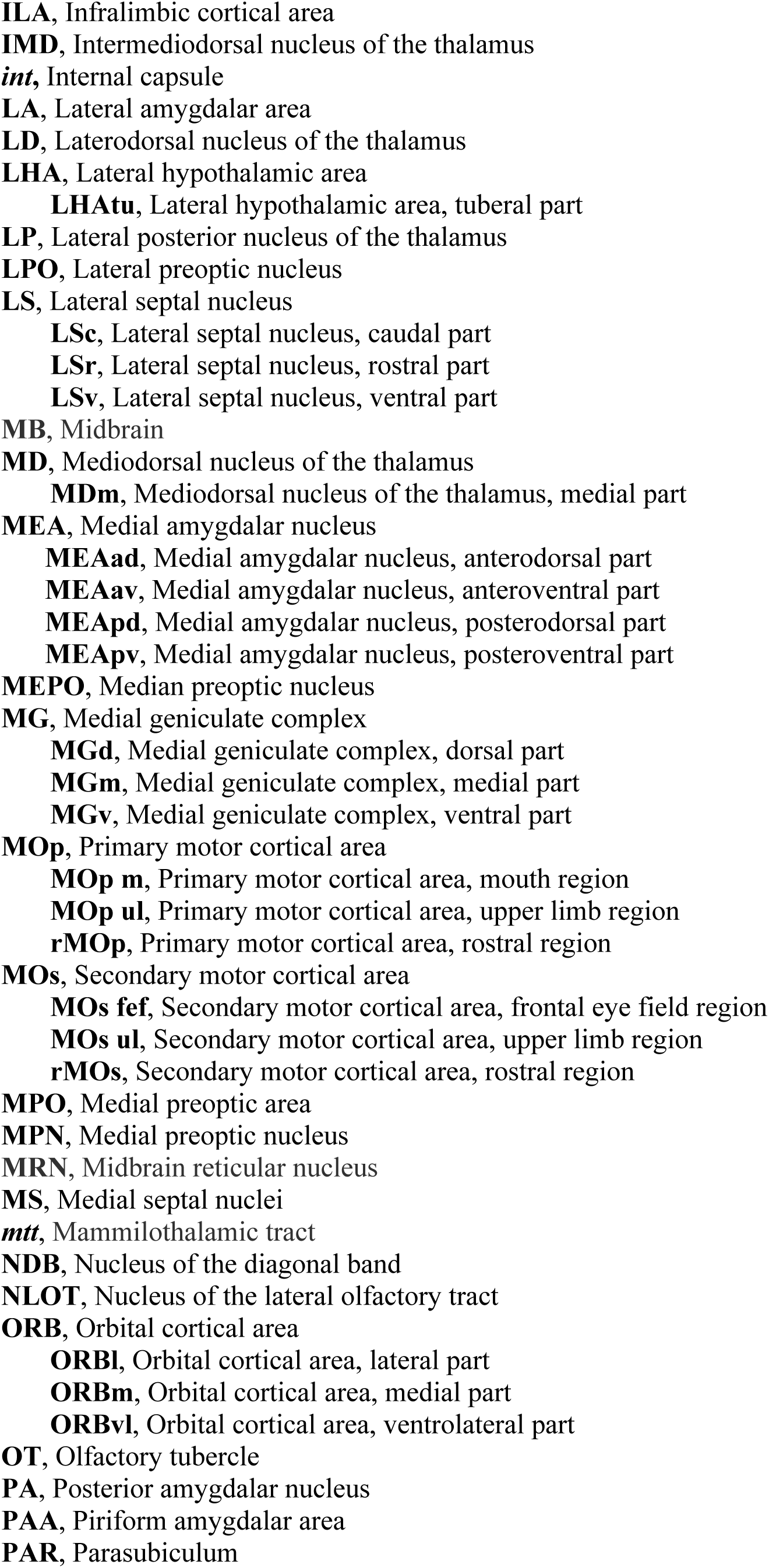
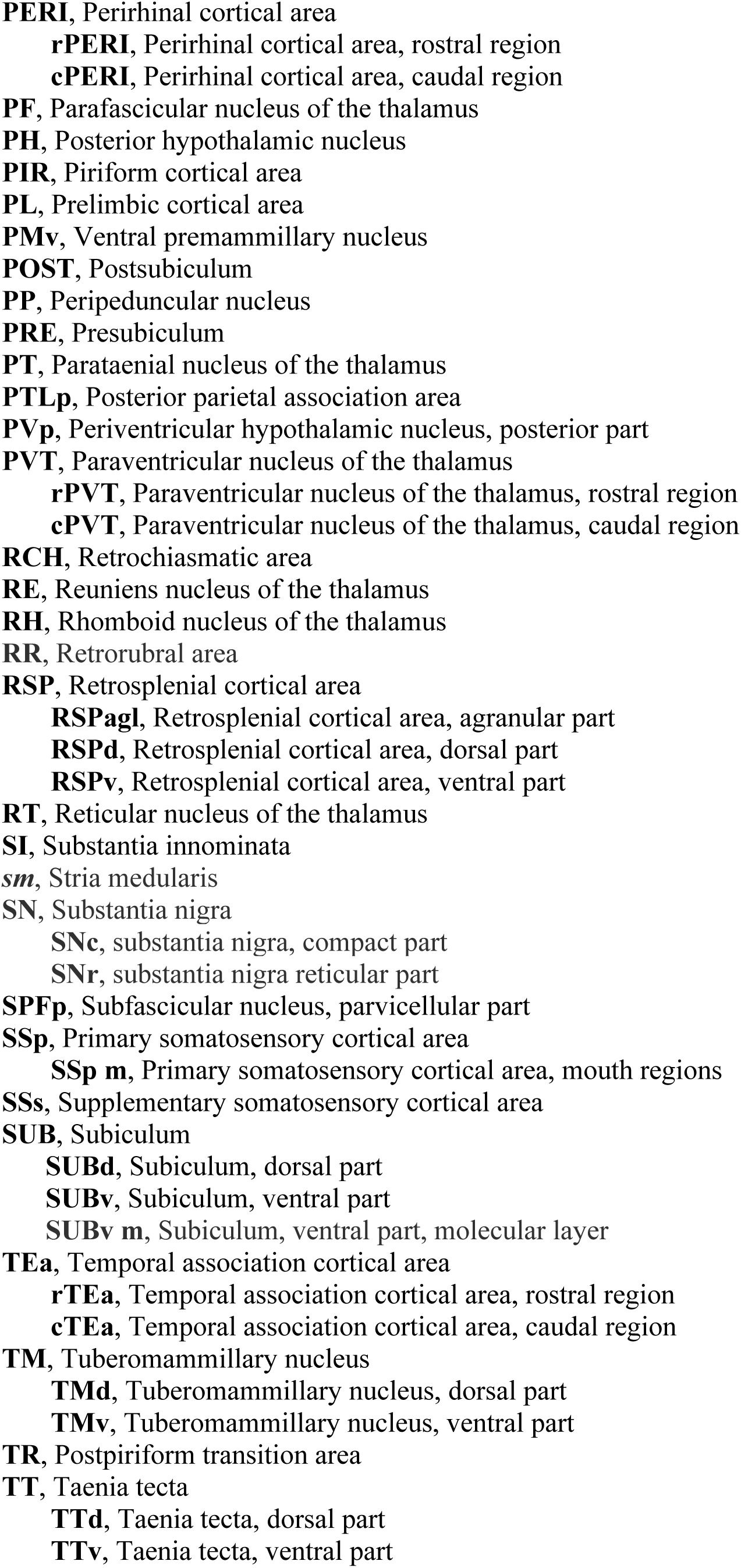
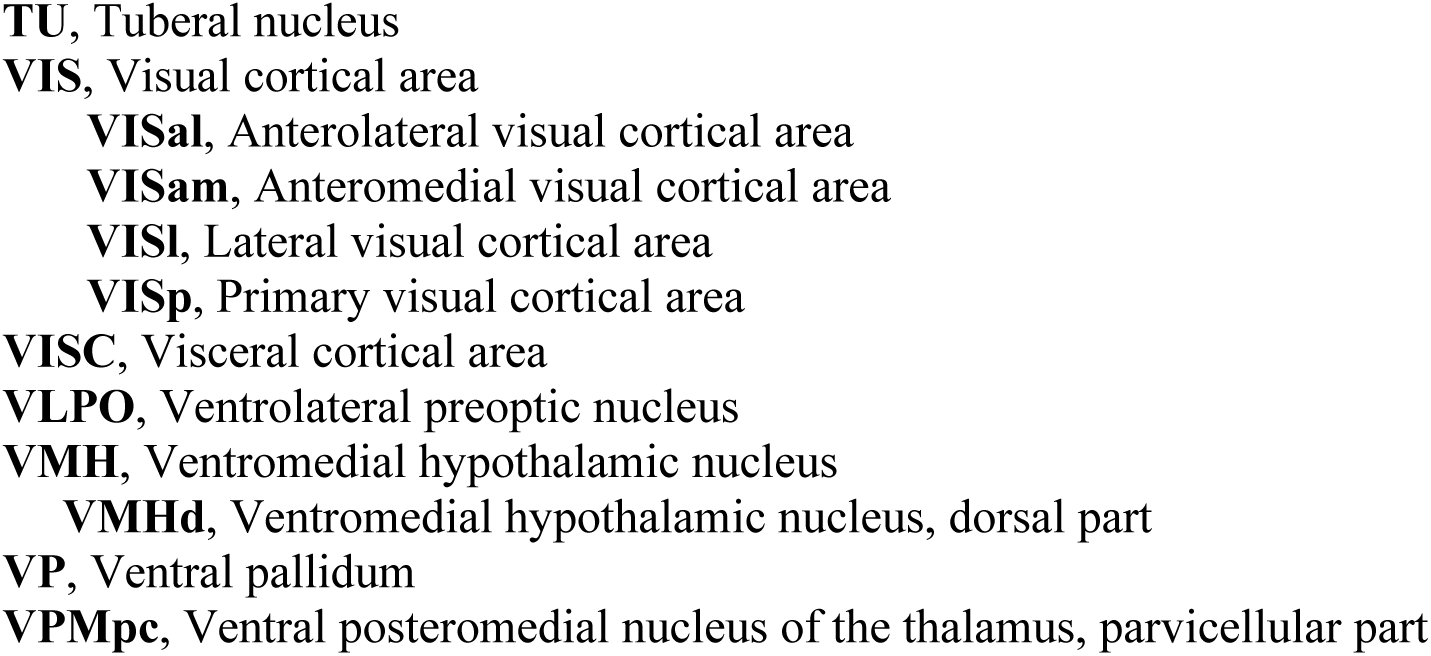
List of abbreviations

Generally, only the strongest connections are reported, all validated with at least one of the following methods. First, injections for all BLA regions were replicated and the consistency across the labelling patterns was manually assessed (Supplementary Figure 4B). Second, retrograde tracers were injected into regions of anterograde terminal labeling to validate the connections from the anterograde tracing, and also to reveal the location of the target-specific BLA projection neurons (Figure 2C). Anterograde tracers were injected into the location of back-labeled projection cells to validate the retrograde BLA injection data. Third, Cre-dependent anterograde AAV tracing experiments were conducted for some of the regions for validation purposes (Supplementary Figure 5). The BLA wiring diagram provided in the current report utilizes data from a total of 153 cases with 195 injections. Raw data for each BLA case is available online through the iConnectome web portal at www.MouseConnectome.org.

Lastly, to functionally characterize synaptic innervation of a few projection-defined BLA cell types, BLA projection neurons and their inputs were labeled with retrograde red beads and AAV-hSyn-ChR2-YFP (ChR2), respectively. Whole-cell voltage-clamp recordings were made from red bead-labeled BLA projection neurons during optogenetic stimulation of ChR2 fibers. Details of all methods are provided in the Online and Supplemental methods.

### Parcellation of the BLAa based on connectivity-defined cell types projecting to the medial prefrontal cortex (MPF) and hippocampal formation (HPF)

Previous studies have demonstrated that the BLAa shares extensive bidirectional connectivity with MPF areas like the infralimbic (ILA) and prelimbic (PL) cortical areas ^29–32^. These bidirectional connections between the BLAa and the ILA and PL were validated. Further, it was demonstrated that input to each cortical area originates from two regionally distinct neuronal populations in the BLAa. When a coinjection of an anterograde and retrograde tracer (BDA/FG) was made into the PL, anterogradely labeled fibers and retrogradely labeled projection cells overlap in the medial part of the BLAa (BLA.am), suggesting strong BLA.am→PL and PL→BLA.am connections (Figure 3A). A second coinjection (Phal/CTb) in the ILA of the same animal revealed that axonal terminals arising from the ILA also concentrate in the BLA.am (Figure 3A). However, the vast majority of CTb retrogradely labeled projection neurons distributed in the lateral part of the BLAa (BLA.al) (BLA.al→ILA) (Figure 3A,C), thereby forming a BLA.al→ILA→BLA.am connectivity circuit. This data suggested that the BLA.am and BLA.al potentially house connectionally-distinctive cell populations.

A Phal/CTb coinjection made into the BLA.am confirmed this, but also revealed the detailed regional and laminar specificities of the BLA.am-MPF connections. Dense axons arising from the BLA.am neurons strongly innervate layer II of PL, but also layers V and VI [BLA.am→PL(II,V,VI)]. ILA remains relatively devoid of these inputs, corroborating the limited BLA.am→ILA connection (Figure 3B). Further, retrogradely labeled PL projection neurons that target the BLA.am distribute primarily in layers II and III, and only in layer III of the ILA, confirming and refining the ILA(III)→BLA.am↔PL(II) pathway. Finally, only layer II of the PL contained intermingled anterogradely labeled axons and retrogradely labeled neurons suggesting a direct reciprocal connection with the BLA.am, while PL layers III (containing retrogradely labeled neurons projecting to BLA.am) and VI (receives light inputs from BLA.am) were predominantly unidirectional projections. A Phal/CTb coinjection in PL layer III confirmed the sparser PL(III)→BLA.am and BLA.am→PL(III) connections (Supplementary Figure 6A). Further, Phal and FG injections in the BLA.al validated the unidirectional BLA.al→ILA connection and revealed that inputs are to layers II-V [BLA.al→ILA(II-VI)] (Figure 3C).

The caudal subdivision of the BLAa (BLA.ac) initially was identified based on its unique projections to the hippocampus ^33^ and shows MPF connections distinct from those of the BLA.am and BLA.al. A retrograde tracer injection in the ventral division of CA3 (CA3v) selectively labeled projection neurons in the BLA.ac, while neurons in rostral sections (BLA.am) remained devoid of tracer label (Figure 4A). This connection was validated with a Phal injection in the BLA.ac, which showed labeled terminals in the CA3i and CA3v (Figure 4B). This subdivision also shares unique connections with the medial prefrontal cortex that differs from its medial and lateral BLAa counterparts (Figure 3F-G). The strongest projections from BLA.ac neurons to the MPF are to PL, primarily layer II, but also to layers III-VI, which only lightly projects back to the BLA.ac (Figure 3 D,F). Most PL projections to the BLAa are to the medial domain (PL→BLA.am). The strongest input to the BLA.ac is from ILA neurons [ILA(II/III)→BLA.ac] (Figure 3D), which in turn is the target of some BLA.ac projection neurons [BLA.ac→ILA(II-VI)] (Figure 3E-G). See Figure 3F-G for summarized connections of BLA and MPF.

#### Global networks of the BLA.am, BLA.al, and BLA.ac

A summarized overview of the brain-wide targets of BLA.am, BLA.al, and BLA.ac projection neurons demonstrates the discrete connectional patterns of each domain. For example, BLA.am and BLA.al neurons target a larger proportion of brain areas in rostral sections (ARA levels 33-73), while BLA.ac neurons target a larger proportion of brain structures in caudal sections (ARA levels 85-99) (Figure 1E).

### Cortical connections

#### Dorsal peduncular area (DP) and anterior cingulate cortical area (ACA)

The BLAa connects with MPF cortical regions outside of the ILA and PL. Projection cells in the BLA.al provide input to the deeper layers of the DP [(BLA.al→DP(II,III)], without receiving much reciprocated input (Supplemental Figure 6D-F). Further, the BLA.am contains projection neurons that strongly target the anterior cingulate cortex (ACA), and in more caudal sections, the adjacent secondary motor area (MOs) region suggested previously to correspond to the frontal eye field of the MOs (MOs-fef) ^34, 35^ (Figure 5A,D). These primarily unidirectional projections are directed toward both the dorsal (BLA.am→ACAd) and ventral (BLA.am→ACAv) divisions of the ACA across its rostral-caudal axis (Figure 5AX) (Supplementary Figure 6B). Only the rostral regions of the ACAd (ARA 29-33) contain projection neurons that project back to the BLA.am (Supplementary Figure 6B).

**Figure 5:**
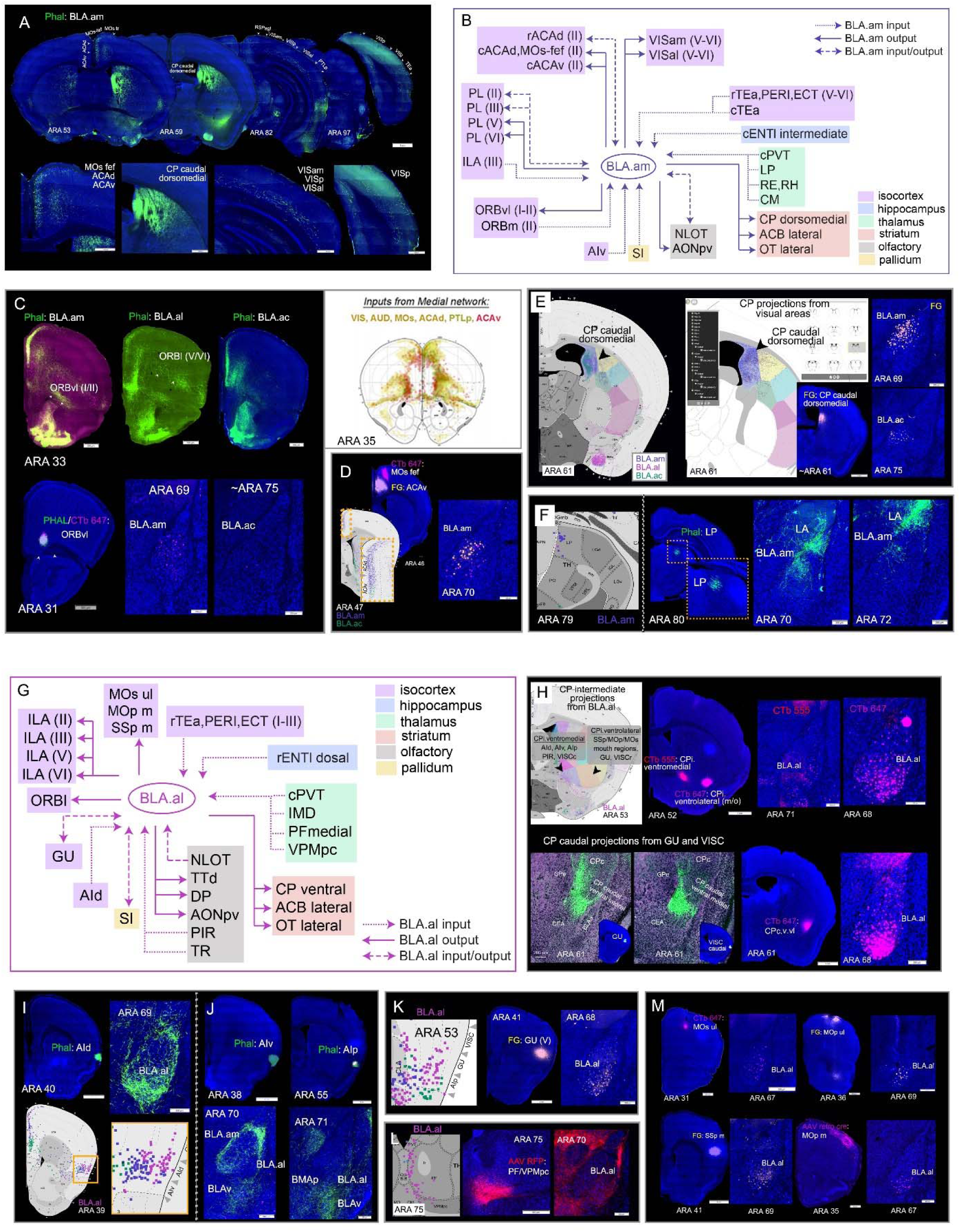
**A.** Projections from BLA.am neurons to regions associated with visual processing. Top panels show whole brain sections with Phal labeled fibers in ACAd, ACAv, MOs-fef, CP caudal dorsomedial, and deeper layers of VISam and VISal following a Phal injection in the BLA.am. Bottom panels show magnified versions of Phal labeled fibers in visual associated areas. *Abbreviations: ACAd: anterior cingulate cortical area, dorsal part; ACAv: anterior cingulate cortical area, ventral part; MOs-fef; secondary motor area, frontal eye field part.* **B.** Summarized brain-wide connections of BLA.am projection neurons. For full list of abbreviations see Table 1. **C.** BLAa connections with ORBvl. BLA.am neurons project to ORBvl (I/II), while BLA.al neurons project to ORBl (V/VI). Although projections can be seen in ORBvl in BLA.ac case, those connections were not validated with a CTb ORBvl injection, which solely back-labeled BLA.am neurons confirming only the BLA.am→ORBvl connection. Further, neurons in the ORBvl do not project back to BLA.am as shown by the Phal ORBvl injection. Schematic adapted from Zingg et al., 2014, demonstrating the pattern of inputs to frontal cortex from regions that process visual information like VIS, ACAd, ACAv, and PTLp. Note the similar pattern of Phal labeling to frontal cortical areas at ARA 33 from the BLA.am Phal injection. *Abbreviations: ORBvl: orbital area, ventrolateral part; ORBl: orbital area, lateral part; VIS: visual cortical areas; ACAd/ACAv: anterior cingulate cortex, dorsal and ventral parts; PTLp: posterior parietal association areas.* **D.** Anterograde visual shows BLA.am projections to ACAv and MOs-fef validated with retrograde injections of CTb and FG (BLA.am→ACAv/MOs-fef). **E.** Left: anterograde visual of BLA.am and BLA.ac projection fibers to CP caudal dorsomedial at ARA 61 superimposed with domains of the CP caudal (Hintiryan et al., 2016). Right: a screenshot of our cortico-striatal map at ARA 61 showing projections from visual areas like VISp (primary), VISam (anteromedial), VISal (anterolateral), ACA (anterior cingulate cortical area), RSP (retrosplenial cortical area), and PTLp (posterior temporal association areas) to the CP caudal dorsomedial (http://www.mouseconnectome.org/CorticalMap/page/map/5). A FG injection in this CP caudal dorsomedial region clearly back-labels BLA.am neurons confirming the BLA.am→CPc.dorsomedial connection. **F.** Left: retrograde visual with back-labeled neurons in visual LP following a retrograde tracer injection into BLA.am. Right: Phal injection in visual LP resulting in labeled fibers in BLA.am, and also the LA, confirming the LP→BLA.am and LP→LA projections. **G.** Summarized brain-wide connections of BLA.al neurons. For full list of abbreviations see Table 1. **H.** Top left shows anterograde visual of BLA.al projection fibers at ARA 53 superimposed with domains of the CP intermediate (CPi) (Hintiryan et al., 2016). BLA.al projection neurons target domains in ventral CPi, which also receive input from AI (agranular insular cortex), PIR (piriform cortex), VISC (visceral cortex), GU (gustatory cortex), and somatosensory and somatomotor regions associated with mouth (m/o). Retrograde tracers CTb 555 (red) and CTb 647 (pink) in ventral CPi back-label projection neurons in BLA.al. Bottom panels: two distinct regions identified in caudal ventral CP that receive strong input from GU and VISC (Hintiryan et al., 2016) also receive input from BLA.al projection neurons. A CTb injection in CP caudal ventral back-labels BLA.al neurons confirming the BLA.al→CPc.ventral projection. **I.** Top: AId neurons target BLA.al and LA as shown with an AId Phal injection (top). Bottom: BLA.al retrograde injection back-labels AId neurons (AId→BLA.al). **J.** AIv shows some weak projections to BLA.am and stronger projections to LA and BLAv. AIp neurons target mostly BLAv and BMAp. *Abbreviations: AId: agranular insular cortical area, dorsal part; AIv: agranular insular cortical area, ventral part; AIp: agranular insular cortical area, posterior part.* **K.** Retrograde visual shows BLA.al neurons in gustatory cortical area (GU) that project to BLA.al (GU→BLA.al). A FG injection in GU(V) reveals BLA.al projection neurons that target deeper layers of GU [BLA.al→GU(V)]. **L.** Retrograde visual shows back-labeled neurons in PF/VPMpc from BLA.al retrograde injection suggesting a PF/VPMpc→BLA.al connection. An anterograde AAV-RFP injection in the thalamic region validates the projection, showing some fiber labeling in BLA.al. *Abbreviations: PF: parafascicular nucleus; VPMpc: ventral posteromedial nucleus of thalamus, parvicellular part.* **M.** BLA.al projection neurons target somatomotor and somatosensory regions presumably associated with orofacial information processing including the MOs/MOp ul (upper limb) and SSp m/MOp m (mouth). Retrograde tracer injections in these regions clearly label BLA.al neurons. *Abbreviations: MOs: secondary motor area; MOp: primary motor area; SSp: primary somatosensory area*.

#### Orbitofrontal cortical areas (ORB)

Projection neurons in the BLA.am specifically target the superficial layers of the ventrolateral division of the ORB (ORBvl), while those in the BLA.al target the deeper layers of the lateral division (ORBl) [BLA.am→ORBvl(I/II); BLA.al→ORBl(V/VI)] (Figure 5C; Supplementary Figure 6I). Strong input from the ORBvl and ORBl back to these BLAa domains was not detected. Instead, input from the medial division of the ORB (ORBm) to the BLA.am and BLA.ac were notable [ORBm(II/III)→BLA.am/BLA.ac] with some reciprocated input [(BLA.am/BLA.ac→ORBm (II-VI)] (Supplementary Figure 6J).

#### Agranular insular (AI) and gustatory (GU) cortical areas

The BLAam and BLA.al share unique connections with the agranular insular (AI) cortical areas. Projections from BLAa to AI are observable. BLA.am neurons project to deeper layers of ventral division of the AI (AIv) [BLA.am→AIv(V/VI)]. Notable and topographic projections from AI projection neurons to the BLAa are also evident. The BLA.al is the primary recipient of input from the dorsal division of the AI (AId) (AId→BLA.al) (Figure 5I; Supplementary Figure 6K), while the BLA.am is the primary recipient of inputs from the ventral division of the AI (AIv) (AIv→BLA.am) (Figure 5J). Projections from AI to the BLA.ac were relatively sparse. In addition, BLA.al projection neurons that send axonal connections to the dorsally adjacent gustatory cortical area (GU) were evident (BLA.al→GU) and neurons in the GU(II-III) project back to the BLA.al [GU(II/III)→BLA.al] (Figure 5K).

#### Sensory cortical areas

The BLAa is directly connected with only a few sensory cortical areas. The BLA.am projection neurons target a region of the MOs previously identified as the MOs-fef (Figure 5A,E). In addition, BLA.am neurons send sparse terminations to deep layers of primary visual cortical areas (VISp), anteromedial (VISam) and anterolateral (VISal) [(BLA.am→VISam/VISal(V/VI)] (Figure 5A; Supplementary Figure 6C). The BLA.al on the other hand, projects to more rostral parts of the MOs, primary motor (MOp), and somatomotor (SSp) regions presumed to be associated with upper limb and orofacial information processing, respectively (Figure 5M).

#### Perirhinal (PERI), ectorhinal (ECT), and temporal association (TEa) cortical areas

Projections from the BLAa neurons to these three interconnected cortical areas are rather sparse. Some observable projections can be seen from the BLA.ac to the more caudal regions of the ECT, although the strongest projections to ECT arise from LA projection neurons (Supplementary Figure 7A) (X). Input from cells in PERI, ECT, and TEa to the BLAa are far greater and predominantly topographically arranged. Neurons in more superficial layers of the rostral TEa, PERI, and ECT project primarily to BLA.al [TEa/PERI/ECT(I-III)→BLA.al] (Supplementary Figure 7B-C), while those in the deeper layers target the BLA.am [PERI/ECT(V-VI)→BLA.am] (Supplementary Figure 7D). Strongest input to BLAa from TEa is provided by its more caudal region to the BLA.am (caudal TEa→BLA.am) (Supplementary Figure 7E). The rostral PERI/ECT/TEa shares more connections with somatic sensorimotor cortex, while its caudal regions are connected with visual cortices. See Figure 3H-I for BLA-isocortical connectivity matrices.

#### Claustrum (CLA)

Generally, the connections between the CLA and BLA are weak and the BLA.am, BLA.al, and BLA.ac are the only nuclei that send weak projections to the subcortical structure (Supplementary Figure 7I). The BLA does not receive much claustral input (BLA.am/BLA.al/BLA.ac→CLA) (Supplementary Figure 7I).

### Hippocampal formation

#### Lateral (ENTl) and medial (ENTm) entorhinal cortical areas

Direct communication between the BLAa and ENT is through strong topographic projections arising from ENTl neurons that terminate in the BLAa (Supplementary Figure 8A-C). Cells in the rostral (and generally dorsal) regions of the ENTl target the BLA.al (rostral/dorsal ENTl→BLA.al) (Supplementary Figure 8B). More caudal intermediate ENTl cells target the BLA.am (caudal/intermediate ENTl→BLA.am) and the most ventrally located ENTl cells target the BLA.ac (caudal/ventral ENTl→BLA.ac) (Supplementary Figure 8F). Connections between the BLAa and the ENTm are present, but sparse, and mostly mediated by the BLA.ac (BLA.ac→caudal ENTm; caudal ENTm→BLA.ac) (Supplementary Figure 8F).

#### CA1, CA3, and subiculum (SUB)

Of the three BLAa subdivisions, the BLA.ac is most strongly connected with hippocampal structures. Neurons in the BLA.ac target the CA3 (BLA.ac→CA3i/v), a defining connection of the BLA.ac (Figure 4A). In addition, BLA.ac projection neurons target the pyramidal layers of CA1 (BLA.ac→CA1_sp) (Figure 4D) and ventral subiculum (BLA.ac→SUBv) (Figure 4D) as well as the parasubiculum (BLA.ac→PAR) (Figure 4A-B), all regions that send inputs back to the BLA.ac (CA1_sp/SUBv/PAR→BLA.ac) (Figure 4E; Figure 6H). The connectivity with the CA3 and PAR are primarily unique to BLA.ac considering all amygdalar nuclei examined. See Figure 4H for summarized connections of BLA and HPF and Figure 4F-G for BLA-hippocampal connectivity matrices.

**Figure 6:**
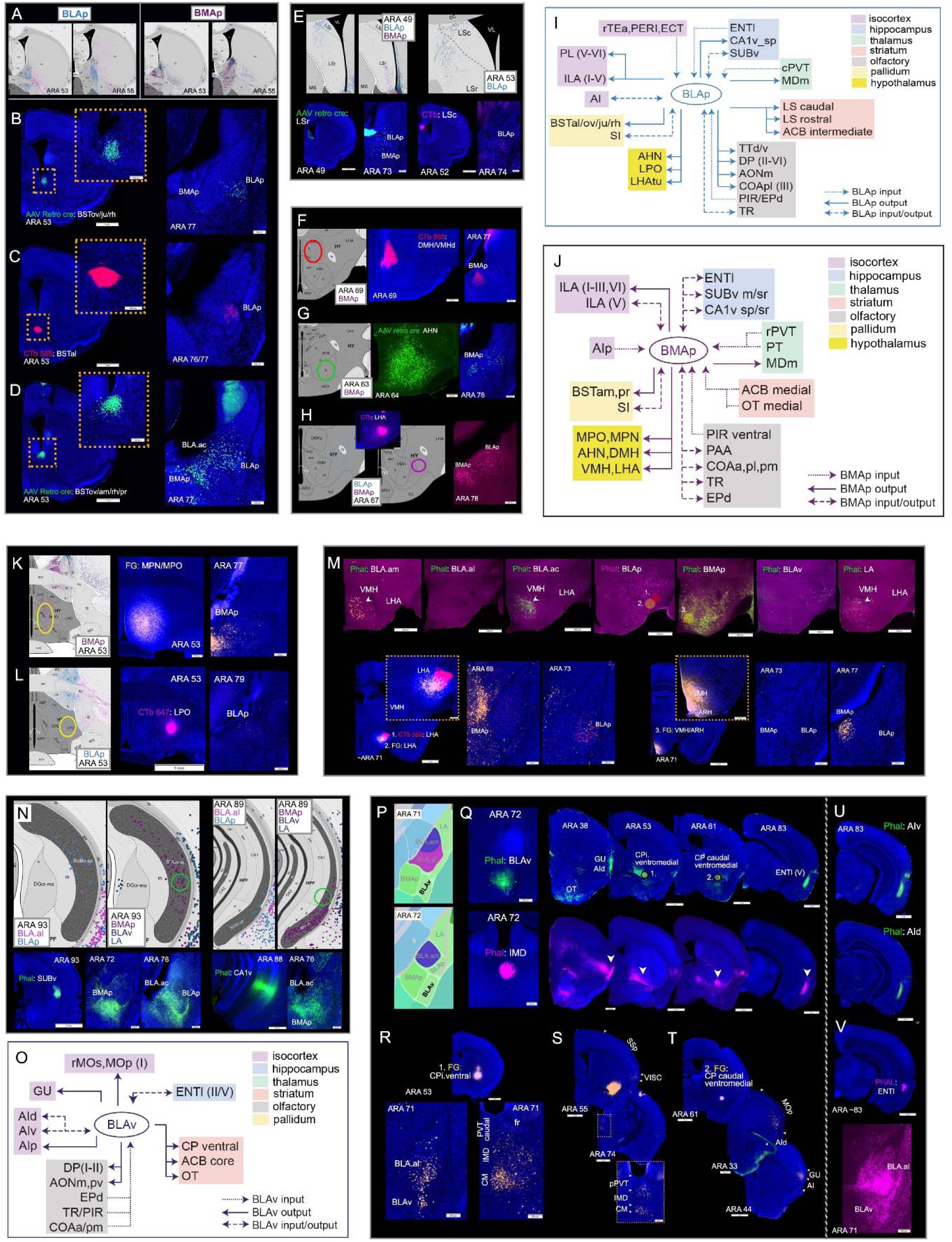
**A.** Anterograde visuals showing BLAp and BMAp neuron projections to BST at ARA levels 53 and 55. Note that BLAp neurons target more lateral parts of BST, while those in BMAp target medial BST. **B-C** validate BLAp projections to BST. Retrograde tracers in BSTov/ju/rh (B) and BSTal (C) back-label BLAp neurons validating the BLAp→BSTal/ov/ju/rh projections. Note the absence of labels in BMAp. Insets are magnification of injection site regions. **D.** Shows validation of BMAp→BSTam/pr projection. A retrograde tracer placed more medially than those in B and C such that it includes BSTam now labels BMAp neurons and some neurons in BLA.ac and LA. Inset shows magnification of injection site region. *Abbreviations: BST: bed nucleus of stria terminalis (BST); BSTov: BST anterior division, oval nucleus; BSTju: BST anterior division, juxstacapsular nucleus; BSTrh: BST anterior division, rhomboid nucleus; BSTal: BST anterior division, anterolateral nucleus; BSTam: BST anterior division, anteromedial nucleus; BSTpr: posterior division, principal nucleus.* **E.** Top panels: BLAp neuron projections to rostral (LSr) and caudal (LSc) lateral septal nuclei and BMAp projections to LSr. Bottom panels validate these connections (BLAp→LSr/c and BMAp→LSr). **F-G.** Retrograde injection in dorsomedial hypothalamic nucleus (DMH: F) and anterior hypothalamic nucleus (AHN: G) validates BMAp neuron projections to the hypothalamic regions (BMAp→DMH/AHN). **H.** Validation of BMAp/BLAp to lateral hypothalamic area (LHA) via LHA retrograde injection. Ellipse on anterograde visual denotes location of Phal injection for validation. **I.** Summarized brain-wide connections of BLAp neurons. **J.** Summarized brain-wide connections of BMAp neurons. For full list of abbreviations see Table 1. **K.** FG injection in medial preoptic nucleus (MPN)/medial preoptic area (MPO) validates BMAp neuron projections to those hypothalamic nuclei (BMAp→MPN/MPO). Ellipse on anterograde visual to the left denotes location of FG injection. **L.** CTb injection in lateral preoptic area (LPO) validates BLAp neuron projections to the hypothalamic nucleus (BLAp→LPO). Ellipse on anterograde visual to the left denotes location of FG injection. **M.** Top panels: projections from BLA.am, BLA.al, BLA.ac, BLAp, BMAp, BLAv, and LA to the tuberal region of the LHA and to ventromedial hypothalamic area (VMH). Strongest projections to VMH are from BMAp neurons, and strongest input to LHA are from neurons in both BLAp and BMAp. Bottom panels validate these connections. Retrograde tracer injections 1 (CTb) and 2 (FG) in LHA back-label both BLAp and BMAp projection neurons (BMAp/BLA→LHA), while FG injection 3 in VMH back-labels selectively BMAp projection neurons (BMAp→VMH). Insets show magnification of injection site regions. **N.** Top panels: retrograde visuals showing SUBv back-labeled cells from retrograde tracer injections in BLAp and BMAp (left) and CA1v labeled cells from BMAp retrograde tracer injection (right). Ellipses denote locations of Phal injections in bottom panels. Bottom panels: Phal injection in SUBv that labels BLA.ac, BLAp, and BMAp validating SUBv→BLA.ac/BLAp/BMAp connections. Phal injection in CA1v labels fibers in BLA.ac and BMAp, but not BLAp validating the CA1v→BLA.ac/BMAp connections. Retrograde visuals for BLA.ac in Figure 4E. **O.** Summarized brain-wide connections of BLAv neurons. For full list of abbreviations see Table 1. **P.** Shows the location of BLAv at ARA atlas levels 71 and 72. **Q.** Top panels show anterograde projections from BLAv, while bottom ones show anterograde projections from intermediodorsal thalamic nucleus (IMD) injection. Note the similarity in labeling from the two injection cases in the GU (gustatory cortical area), AId (agranular insular area, dorsal part), CP intermediate (CPi) ventromedial, CP caudal (CPc) ventromedial, and ENTl (entorhinal cortical area, lateral part) particularly layer V. **R.** FG injections in CP intermediate ventromedial back-labels projection neurons in BLA.al and BLAv (BLA.al/BLAv→CPi.ventral), but also in thalamic nuclei CM (central medial) and IMD. **S-T.** FG injection in CP caudal ventromedial, where BLAv and IMD project, back-labels neurons in regions proposed to be involved in gustatory/visceral processing like IMD, CM, VISC (visceral cortical area), MOp mouth regions, GU (gustatory cortical area), and AI (agranular insular cortical area). **U.** Like the BLAv and IMD, the AI also projects to ENTl(V) and a Phal injection in ENTl(V) shows that it projects back to BLAv, but also to BLA.al [ENTl(V)→BLAv/BLA.al].

### Olfactory connections

All three BLAa domains are connected with regions that process olfactory information. The BLA.am and BLA.al cells also project to the anterior olfactory nucleus posteroventral part (AONpv) (BLA.am/BLA.al→AONpv), without reciprocated input (Supplementary Figure 9A). The BLA.am and BLA.al cells project most strongly to all layers of the nucleus of the lateral olfactory tract (NLOT) (BLA.am/BLA.al→NLOT) and layer III NLOT cells project back to the BLA.am and BLA.al [(NLOT(III)→BLA.am/BLA.al] (Supplementary Figure 9C-D). Olfactory input is provided by projection cells located in the ventral part of the PIR that target most strongly the BLA.al and BLA.ac (ventral PIR→BLA.al/BLA.ac) (Supplementary Figure 9E). The BLA.al is also the recipient of inputs from neurons located in more dorsal parts of the PIR (dorsal PIR→BLA.al) (Supplementary Figure 9F). Another cortical olfactory areas that connects with the BLA.al includes the piriform-amygdalar transition area (TR). Specifically, TR cells provide some input to the BLA.al (TR→BLA.al) (Supplementary Figure 10H).

### Thalamic connections

Projections from BLAa neurons to the thalamus are light; however, all three domains primarily are innervated by neurons in the midline thalamic nuclei. Generally, projection neurons in rostral parts of the midline nuclei (ARA 57, 61) mostly target the BLA.ac, while the mid-caudal midline thalamic nuclei (ARA 69, 73, 75) target the BLA.am and BLAal (Supplementary Figure 11A). This is most evident with the parataenial (PT) and paraventricular (PVT) projections to the BLA.ac (Supplementary Figure 11A-B). The PVT extends a long distance across the rostro-caudal axis with rostral and caudal subdivisions that are generally accepted, but not delineated in rodent atlases. These divisions are based on anatomical ^36–39^ and behavioral distinctions ^40–42^. For example, the caudal PVT (cPVT) is shown to be a critical unit of the fear circuit ^43, 44^ and for behaviors related to anxiety and stress ^41, 45, 46^, while the rostral PVT (rPVT) is shown to be involved in ethanol consumption ^47^. Projection neurons in the rPVT preferentially target the BLA.ac (rPVT→BLA.ac) (Supplementary Figure 11B-C), while mostly segregated neuronal populations in the more cPVT primarily project to the BLA.am and BLA.al (cPVT→BLA.am/BLA.al) (Supplementary Figure 11B,F). Additional discriminating thalamic inputs are observed with PT neurons that primarily target the BLA.ac (PT→ BLA.ac) (Supplementary Figure 11B,D) and with the intermediodorsal (IMD), parafascicular [PF, medial part mPF)], and VPMpc neurons that primarily innervate the BLA.al (IMD/mPF/VPMpc→BLA.al) (Figure 5F; Supplementary Figure 11B,F). The BLA.am receives input from midline thalamic nuclei that also project to BLA.ac like the dorsal anteromedial thalamic nucleus (AMd), the reunions (RE), rhomboid (RH) and central medial (CM) (RE/RH/CM→BLA.am/BLA.ac) (Supplementary Figure 11B,E). Common thalamic input to both BLA.am and BLA.al is provided by the cPVT (cPVT→BLA.am/BLA/al) (Supplementary Figure 11B,F). Projections from cells in the visual lateral posterior thalamic nucleus (LP) are unique to the BLA.am (LP→BLA.am) (Figure 5F).

### Output to motor systems

#### Dorsal striatum: the caudoputamen (CP)

Projection neurons in all three BLAa domains target different regions of the CP. At the intermediate CP level CP (CPi, ARA 53), the BLA.al neurons target the ventromedial (CPi.vm) and ventrolateral (CPi.vl) regions, which are also targeted by the AI, PIR, VISC and by GU and somatosensory and somatomotor regions related to mouth areas, respectively ^34^ (BLA.al→CPi.vm/CPi.vl) (Figure 5H). At more caudal levels, BLA.al neurons target the ventral parts of the CP (CPc.v), whose medial aspect (CPc.v.vm) is heavily innervated by the GU and the lateral aspect (CPc.v.vl) by the VISC cortex ^34^ (BLA.al→CPc.v) (Figure 5H). In contrast, BLA.am and BLA.ac neurons target the dorsomedial regions of the CPi (CPi.dm; Figure 5A; Supplementary Figure 10B-C), a domain that receives convergent input from all visual cortical areas, the ACA, RSP, and ENTm ^34^ (BLA.am/BLA.ac→CPi.dm) (Supplementary Figure 10D-E). At caudal levels, projection neurons in BLA.am and BLA.ac innervate the dorsomedial domains (CPc.dm) (Figure 5A,E), which receive convergent, integrated information from visual, auditory, and higher order association areas like the ACA, PTLp ^34^ (BLA.am/BLA.ac→CPc.dm) (Figure 5E). Double retrograde injections in the dorsal and ventral parts of the caudal CP validates this and clearly illustrates the segregation of CP projecting BLA.am and BLA.al neuron populations (Figure 1A; Supplementary Figure 1C).

#### Ventral striatum: nucleus accumbens (ACB) and olfactory tubercle (OT)

Of all BLA neurons, those in the BLA.am and BLA.ac most strongly project to OT (Figure 1F). Within the ACB and OT, clear topographic projections originating from BLA.am and BLA.ac neurons are observed. Neurons in the BLA.ac preferentially target the medial aspect of the ACB (Figure 1F) and the OT (BLA.ac→medial ACB/medial OT) (Figure 1F), while BLA.am neurons target the lateral aspects of these striatal structures (BLA.am→lateral ACB/lateral OT) (Figure 1F-G). Some BLA.al neurons sparsely target the lateral OT and more strongly target the lateral ACB (BLA.al→lateral OT/ACB) (Figure 1F-G). Within the ACB, slight segregation of BLA.am and BLA.al projections are notable (Figure 1G).

#### Pallidum: substantia innominata (SI) and bed nuclei of stria terminalis (BST)

In the ventral pallidum, neurons in the BLA.al target the SI (BLA.al→SI) (Supplementary Figure 10I,K). The SI appears to be the only motor nucleus to provide input, although weak, to the BLAa, specifically to the BLA.am and to a lesser extent the BLA.al (SI→BLA.am/BLA.al) (Supplementary Figure 10J-K). In the caudal pallidum, the BLA.ac appears to be the only BLAa nucleus to target the bed nucleus of stria terminalis (BST) specifically its oval (BSTov) and rhomboid (BST.rh) nuclei (BLA.ac→BSTov/rh) (Figure 6B).

Summarized brain-wide connections of projection neurons in BLA.am (Figure 5B; Figure 7E; Supplemental Figure 13F), BLA.al (Figure 5G; Figure 7F; Supplemental Figure 13G), and BLA.ac (Figure 4C; Figure 7G; Supplemental Figure 13H) are provided in wiring diagrams and flatmap visualizations ^48^.

**Figure 7:**
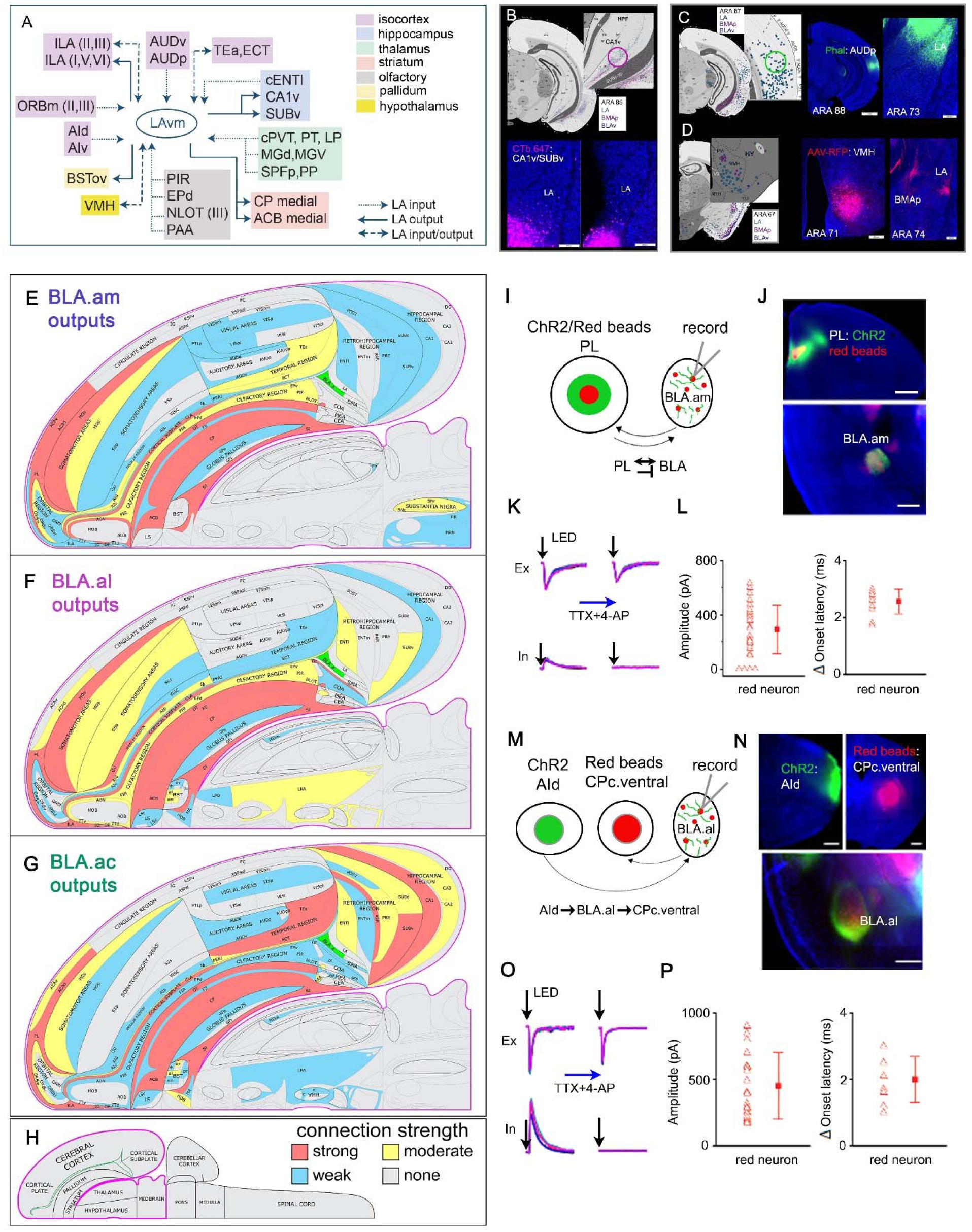
**A.** Summarized brain-wide connections of LA ventromedial neurons. For full list of abbreviations see Table 1. **B.** Anterograde visuals in top panels show weak projections from LA neurons to hippocampal regions SUBv and CA1v. Ellipse denotes location of CTb injection for validation. A CTb 647 validates this sparse connection. **C.** Retrograde visual shows labeled cells in auditory cortical areas (AUD) following a retrograde tracer injection in LA. Ellipse denotes location of Phal injection for validation. A Phal injection in the primary auditory cortical area (AUDp) shows strong anterograde label in LA validating the AUD→LA connection. **D.** LA receives strong input also from the ventromedial hypothalamic nucleus (VMH), as shown by the retrograde visual with labeled cells in the hypothalamic region following the injection of a retrograde tracer in LA. An AAV-RFP injection in VMH validates the VMH→LA connection showing strong label in the LA, but also in BMAp (VMH→BMAp). **E-G**. The outputs of the BLA.am (**E**), BLA.al (**F**), and BLA.ac (**G**) domains are represented at the macroscale (gray matter region resolution) on a partial flatmap representation of the mouse brain. The strength values of detected connections were binned into tertiles, and these are represented qualitatively as strong (red), moderate (yellow), and weak (blue). For definition of abbreviations, see X. **H.** Longitudinal half of the entire CNS flatmap showing orientation and major brain divisions (e.g., cerebral cortex, pallidum, thalamus). The key to the color codes for connection strength is also shown. **I-P.** Functional characterization of synaptic innervation of projection-defined BLA neurons. **I.** Schematic drawing of experimental design. AAV-hSyn-ChR2-YFP and retrograde microbeads (in red fluorescence) are injected into PL. Recordings are made from BLA neurons retrogradely labeled with red microbeads. Proposed circuit connectivity: PL projecting BLA.am neurons are monosynaptically innervated by excitatory PL projections as well as bisynpatic feedforward inhibition in BLA.am. **J.** Images of injection site (Upper) and BLA area (Lower). Note the green florescence from the projections of PL expressing YFP and soma of BLA neurons labeled in red. **K.** An example of whole-cell voltage-clamp recording of a BLA neuron with red fluorescence. Left: excitatory (at −70 mV) and inhibitory (at 0 mV) responses (superimposed traces) evoked by blue light stimulation (5ms). Right, responses from the same recorded cell in the presence of TTX and 4AP. Note only the monosynaptic excitatory responses are elicited. **L)** Left: A summary of amplitudes of monosynaptic excitatory connections recorded from BLA neurons retrogradely labeled from PL injection. Right: Differences in onset latencies between excitatory and inhibitory responses (as exampled in K) recorded from the same neuron. This short difference suggests the inhibition is bisynpatic feedforward input. Proposed circuit connectivity: PL projecting BLA.am neurons are monosynaptically innervated by excitatory PL projections as well as bisynpatic feedforward inhibition in BLA.am. **M-P.** CP projecting BLA neurons are innervated by AId. **M.** Schematic drawing of experimental design. AAV-hSyn-ChR2-YFP is injected in AId, while red retrobeads is injected in CPc.ventral. Recordings are made from CP projecting BLA.al neurons labeled with red retrobeads. Proposed connectivity: CPc.ventral projecting BLA.al neurons are monosynaptically innervated by AId projections. **N.** Images of injection sites (upper) and BLA region (lower). Note the intermingling of green fluorescent AI projections and CP projecting red neurons in BLA.al. **O.** An example of whole-cell voltage-clamp recording of a BLA neuron with red fluorescence similarly plotted as in **K**. Left: excitatory (at −70 mV) and inhibitory (at 0 mV) responses (superimposed traces) evoked by blue light stimulation (5 ms). Right, responses from the same recorded cell in the presence of TTX and 4AP. **P.** Summary of amplitudes and differences in onset latencies as in **L**.

### BLAa neuron morphology

To assess whether connectionally-distinct BLA.am, BLA.al, and BLA.ac projection neurons were morphologically different, representative cells in each domain were labeled via a G-deleted rabies injection in the dorsomedial CPc (Movie 1), ventral CPc (Movie 2), and in the medial ACB (Movie 3, Movie 4, Movie 5), respectively (Figure 1D). The SHIELD clearing protocol for thick sections was followed ^49, 50^ and sections were imaged using an Andor Dragonfly high speed spinning disk confocal microscope. Manual reconstruction and quantitative analysis of neurons was performed using Aivia (Figure 8D) and quantitative analysis was performed using Fiji and L-Measure.

**Figure 8:**
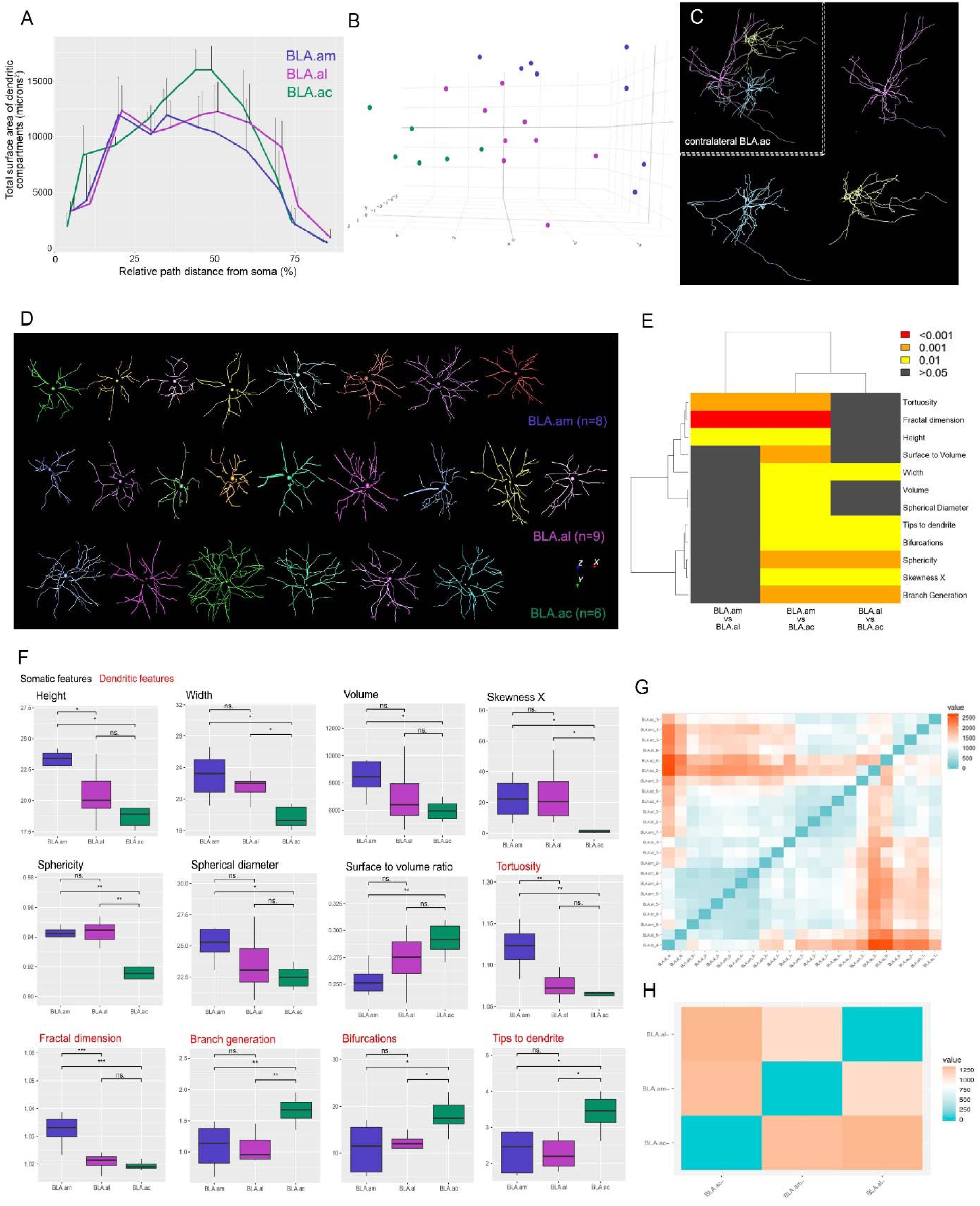
**A.** Result of Sholl-like analysis to show overall view of BLAa projection neuron dendritic morphology. Graph shows that dendrites of neurons within the BLA.ac have a larger surface area of dendritic compartments at ∼45% distance from the cell body compared to dendritic compartments of BLA.am and BLA.al neurons. This larger dendritic surface area suggests the potential for a greater number of synaptic contacts for BLA.ac neurons. **B.** Principal component analysis (PCA) shows segregation of BLAa domain-specific neurons based on measured morphological features (see Supplemental methods). **C.** Contralateral medial accumbens (ACB) projecting BLA.ac neurons. Neurons were labeled via a rabies virus injection in the medial ACB and neurons in contralateral BLA.ac were manually reconstructed. **D.** All reconstructed dorsal striatum projecting BLA.am (n=8) and BLA.al (n=9) neurons and ventral striatum projecting BLA.ac neurons (n=6). These reconstructions were used to assess differences in morphological features across the domain-specific projection neurons. **E-F.** Pairwise tests were run and the parameters that survived the false discovery rate (FDR) correction for multiple testing were reported. The significant group differences are presented with whisker plots in panel F and the degree of their significance is visualized in a matrix in E. For whisker plots, the center line represents the median, box limits show the upper and lower quartiles and the whiskers represent the minimum and maximum values. Somatic features are presented in black font, while dendritic features are in red. The dendrogram on top of the matrix shows the hierarchical clustering of groups based on feature similarity. It suggests that BLA.am and BLA.al neurons differ more from those in BLA.ac than they do from each other. **G.** Persistence-based neuronal feature vectorization framework was also applied to summarize pairwise differences between BLA.am, BLA.al, and BLA.ac projection neurons. The strength of the differences is presented as a gradient with blue showing no difference and orange the greatest differences. The individual neuron differences are aggregated in **H**, which shows once again that neurons within BLA.am, BLA.al, and BLA.ac all differ from one another, but that the greatest difference lies between BLA.am/BLA.al neurons versus BLA.ac neurons.

Standard measurements obtained for cell body morphology included volume, sphericity, spherical diameter, X, Y, Z and Euclidean skewness, cell body height, width, depth, and surface to volume ratio. Measurements for dendritic morphology included number of primary dendrites, branches, bifurcations, terminal tips, nodes, fractal dimension, tortuosity, local bifurcation angle, remote bifurcation angle, partition asymmetry, tips to dendrite ratio, branch generation, and Rall’s ratio. Pairwise Wilcoxon rank sum tests were run and the parameters that survived the false discovery rate (FDR) correction for multiple testing are reported. The significant group differences are presented with whisker plots (Figure 8F) and the FDR q-value significance level is visualized in a matrix (Figure 8E). Furthermore, we used principal component analysis (PCA) to reduce the dimensionality of all measured morphological parameters and create a 3D scatterplot illustrating the segregation of BLAa domain specific neurons based on the measured features (Figure 8B) ^51, 52^.

Statistically significant differences in several morphological features were detected across all pairwise comparisons; however, neurons in BLA.am and BLA.al generally showed greater similarity to one another than they each did to BLA.ac. Figure 8E shows a matrix and dendrogram plot created by hierarchical clustering with group similarity, demonstrating a greater number of significantly different features for BLA.ac neurons. In addition, the box plots in Figure 8F show the size of the relative differences between BLA.ac and the other divisions. We further corroborated these findings through Sholl-type and topological analysis. First, the traditional Sholl analysis showed that dendrites of BLA.am and BLA.al neurons mostly differed from those in BLA.ac within 100-200 nm radius from the cell body, with BLA.ac dendrites showing a greater number of intersections within that range (Supplementary Figure 1I). Second, a three-dimensional Sholl-like analysis showed that in BLA.ac neurons the distribution of dendritic surface area was more peaked towards the median of the relative path distance from the soma as opposed to the more uniformly distributed dendritic surface of BLA.am and BLA.al neurons (Figure 8A). Finally, persistence-based neuronal feature vectorization framework was also applied to summarize pairwise differences between BLA.am, BLA.al, and BLA.ac projection neurons, which showed once again that neurons within BLA.am, BLA.al, and BLA.ac all differ from one another, but that the greatest difference lies between BLA.ac neurons and those in BLA.am/BLA.al (Figure 8G-H). Details of methods and statistical output are provided in the Online and Supplemental methods.

#### Global networks of BLAp

Tracing data from the BLAp and BLA.al were aggregated and analyzed to emphasize differences in their connectivity architecture. Analysis showed that projection neurons located in the BLA.al target a different set of structures than those in the BLAp. For example, a large proportion of the label from BLAp neurons reside in the ACB (ARA 47), BST (ARA 53), and CA1 (ARA 91), while those from BLA.al neurons are to ACB, but also CP (ARA 37), CP and FS (ARA 53), and hardly any to ARA 91 (Supplementary Figure 12A-C).

### Cortical connections

Cortical targets of BLAp neurons are located primarily in the MPF. In particular, they heavily innervate the PL and ILA with laminar specificity [BLAp→ILA(I-V)/PL(V-VI)] (Supplementary Figures 1E, 6H). Input from these structures back to the BLAp are sparse (Supplementary Figure 1F). BLAp neurons also strongly target the DP with laminar specificity [BLAp→DP(II-VI)] without receiving significant input from this region (Supplementary Figure 6E-F). Deeper layers of the AI receive some scant input from BLAp neurons [(BLAp→AI(V/VI)] and AI neurons sparsely target the BLAp (AI→BLAp) (Supplementary Figure 6L). At more caudal cortical regions, no appreciable projections from BLAp to PERI/ECT/TEa were detected. Input to BLAp from neurons located in the more rostral parts of these regions were slightly more notable (rostral PERI/ECT/TEa→BLAp) (Supplementary Figure 7H).

### Connections with motor systems

Neurons in the BLAp target several different structures involved in motor output. Like the BLAa, BLAp cells provide input to the ACB, but to its intermediate part (BLAp→intermediate ACB) (Supplementary Figure 10A). Unlike BLAa neurons, cells in the BLAp do not as strongly target the CP (Supplementary Figure 5 B) or the OT (Figure 1F).

Instead, BLAp neurons target the lateral septum (LS), bed nucleus of the stria terminalis (BST), and the hypothalamus. Specifically, they target the caudal (LSc) and rostral (LSr) divisions of the LS (Figure 6E). They also preferentially target the lateral part of the BST including the anterolateral (BSTal), oval (BSTov), juxstacapsular (BSTju), and rhomboid (BSTrh) nuclei (BLAp→BSTal/ov/ju/rh) (Figure 6B-C). The anteromedial (BSTam) receives far less input in comparison (Figure 6A). Hypothalamic structures targeted by BLAp neurons include the lateral preoptic areas (LPO) (Figure 6L), anterior hypothalamic nucleus (AHN) (Figure 6G), and the lateral hypothalamic area (LHA) in particular the tuberal part (BLAp→LPO/LHAtu) (Figure 6M).

The strongest projections from BLAp neurons to hippocampal structures are to the CA1v_sp, but mostly to SUBv (BLAp→CA1v/SUBv) (Figure 4D). Projections from neurons in these areas back to BLAp also are observable (CA1v/SUBv→BLAp) (Figure 6N). Although projections to ENTl from these neurons are sparse (Supplementary Figure 8G), input from ENTl to the BLAp is evident (ENTl→BLAp) (Supplementary Figure 8G,B-C).

Like the BLA.am and BLA.al, neurons in the BLAp also provide input to ventral pallidal structures like the SI. In turn, SI neurons provide input back to the BLAp (Supplementary Figure 10I-K).

### Thalamic connections

Unlike the BLAa, the BLAp does not receive much thalamic input. Some neurons in the caudal PVT provide sparse input to the BLAp (cPVT→BLAp) (Supplementary Figure 11F). Also unlike the BLAa, projection neurons in the BLAp target the medial subdivision of the mediodorsal thalamic nucleus (MDm) (BLAp→MDm) (Supplementary Figure 4E, 11G). Most anatomy literature has demonstrated strong BLA→MD projections ^30, 53^, those projections arise mostly from BLAp and BMAp and not BLAa ^54^.

### Olfactory connections

BLAp neurons target the dorsal taenia tecta (TTd), to a much lesser extent its ventral counterpart the (TTv) (BLAp→TTd/TTv) (Supplementary Figure 9G), and the medial aspect of the AON (BLAp→AONm) (Supplementary Figure 9B). Input from olfactory areas to the BLAp is provided by the PIR (PIR→BLAp) (Supplementary Figure 9E-F) and EPd (EPd→BLAp) (Supplementary Figure 9H). Olfactory cortical areas that connect with the BLAp include the TR. Specifically, BLAp neurons project strongly to the TR (BLAp→TR) and cells in the TR project back to the BLAp (TR→BLAp) (Supplementary Figure 10G-H). Summarized brain-wide connections of projection neurons in BLAp is provided in a wiring diagram (Figure 6I).

#### BLAv, LA, and BMAp

Brain-wide connections of the BLAv, LA, and BMAp were examined to complete the connectional profile of the BLA complex.

### Global networks of BLAv

The BLAv is located ventral to the BLAa and BLAp, adjacent to the BMAp (Figure 6P) and is considered a part of the BLA complex ^1^. Compared to all other nuclei examined, the BLAv displays a selective and unique connectivity architecture. In particular is the striking resemblance between the targets of its projection neurons and the intermediodorsal nucleus of the thalamus (IMD). In cortical areas, strongest targets of the BLAv neurons include the rostral AId, AIv, and GU (BLAv→rostral AId/AIv/GU) (Figure 6Q). In turn, AId, AIv, and AIp neurons all target the BLAv (AId/AIv/AIp→BLAv) (Figure 5J; Supplementary Figure 6L). In more rostral cortical areas, BLAv neurons target layer 1 of the primary and supplemental motor cortices (Supplemental Figure 4F). The only connection between BLAv and MPF is through DP. Similar to the BLA.al, BLAv neurons provide input to the DP [(BLAv→DP(I/II)], while projections from DP back to BLAv were not observed (Supplemental Figure 6G,F).

Another common target of BLAv and IMD projection neurons is a specific, confined region within the rostral ENTl layer V, but also II [(BLAv→ENTl(II/V)] (Figure 6Q), which also receives dense input from AI, GU, and VISC (Figure 6U: Supplementary Figure 8D-E). Whereas the BLA.am, BLA.al, and BLA.ac neurons sparsely target the ENTl, input from BLAv neurons to ENTl(V) is dense. In turn, neurons in this ENTl region project back to BLAv [ENTl(V)→BLAv] (Figure 6V).

BLAv and IMD neurons also target similar motor regions like the CP and ACB. Within the intermediate CP (CPi, ARA 53), BLAv neurons target the ventromedial (CPi.vm) region, which is also targeted by the BLA.al (BLAv→CPi.vm) (Figure 6Q-S). In the caudal CP (CPc, ARA 61), BLAv neurons target ventral parts of the CP, specifically the medial aspect (CPc.v) (Figure 6Q), which is also innervated by the GU ^34^ (Figure 6T). Projection neurons in the BLAv and IMD also target a similar region in the core of the ACB (BLAv→ACBcore) (Supplemental Figure 10A). Projections from BLAv neurons to the OT were also observed (BLAv→OT) (Figure 1F; Figure 6Q).

The most prominent olfactory connection for the BLAv is the PIR. Projection neurons across the entire PIR provide input to the BLAv (PIR→BLAv) (Supplementary Figure 9E-F). Neurons in the dorsal part of the endopiriform nucleus (EPd) (EPd→BLAv) (Supplementary Figure 9H), TR (TR→BLAv) (Supplementary Figure 10H), the anterior (COAa) and posterior medial (COApm) cortical amygdala area (COAa/COApm→ BLAv) (COAa/pm→BLAv) (Supplementary Figure 13B,D) all provide appreciable input to the BLAv. In turn, BLAv neurons target the AONm and AONpv (BLAv→AONm/pv) (Supplementary Figure 9A-B). Summarized brain-wide connections of projection neurons in BLAv is provided in a wiring diagram (Figure 6O).

### Global networks of LA

#### Cortical connections

Similar to the BLA.al, projection neurons in the LA provide reasonable input to the ILA [LA→ILA(I-VI)] and neurons in ILA project back to LA [(ILA(II/III)→LA] (Supplementary Figure 1E-F). ORBm neurons also provide some input to the LA [ORBm(II/III)→LA)] and receive some input from the regions (LA→ORBm) (Supplementary Figure 6J). Within the AI, inputs from the LA neurons are sparse; however, projection neurons in the AId and AIv provide a fair amount of input to the LA (AId/AIv→LA) (Figure 5I-J). Cortical recipients of the strongest projections from LA neurons are the ECT, followed by the TEa (LA→ECT/TEa) (Supplemental Figure 7F) and neurons in the ECT and TEa also strongly target the LA (ECT/TEa→LA) (Supplemental Figure 7G). The LA also receives input from sensory cortical areas. The strongest input is provided by neurons in the primary (AUDp) and ventral (AUDv) auditory areas (AUDp/AUDv→LA) (Figure 7C).

#### Hippocampal formation

Although projections from LA neurons do not target the ENTl, neurons in the more caudal regions of the ENTl provide some input to the LA (ENTl→LA) (Supplementary Figure 8I). Some light projections from the LA to the ventral parts CA1 and SUB were also detected (LA→CA1v/SUBv) (Figure 7B).

#### Thalamic connections

Thalamic structures do not receive input from LA neurons, but projection neurons in several nuclei provide input to LA. These include the caudal PVT, PT, and LP (caudal PVT/PT/LP→LA) (Supplementary Figure 11D,H-I,L). The strongest inputs originate from neurons in the dorsal (MGd) and ventral (MGv) parts of the medial geniculate nucleus (MG) and in the SPFp (subfascicular nucleus, parvicellular part) and PP (peripeduncular nucleus) (MGd/MGv/SPFp/PP→LA (Supplementary Figure 11J-K).

#### Olfactory connections

The LA does not project strongly to olfactory structures; however, neurons in olfactory related regions like the PIR, EPd, and NLOT(III) provide some input to LA [PIR/EPd/NLOT(III)→LA] (Supplementary Figure 9C,E,H). Cortical olfactory areas like the piriform-amygdala area (PAA) also provide some input to the LA (PAA→LA), while others like the COApl receive some input (LA→COApl) (Supplementary Figure 13C).

#### Connections to motor systems

Similar to all other BLA complex nuclei, LA projection neurons strongly target the ACB, in particular a specific region in the medial part of the ACB (LA→ACB medial) (Supplementary figure 10A). LA neurons also weakly target the BST, primarily the oval nucleus (LA→BSTov) (Figure 6D) and the CP (LA→CP), specifically to the CPi.medial, a division that also receives input from the ECT and TEa caudal (Supplementary Figure 10F,D). Finally, although LA neurons provide weak input to the hypothalamic VMH (LA→VMH) (Figure 6M), neurons in the VMH send strong projections back to the LA (VMH→LA) (Figure 7D). Summarized brain-wide connections of projection neurons in LA is provided in a wiring diagram (Figure 7A).

### Global networks of BMAp

#### Cortical connections

The two most notable cortical regions to receive input from BMAp projection neurons is the medial prefrontal and lateral entorhinal cortical areas. Specifically, BMAp neurons target the ILA [BMAp→ILA(II-VI)] and in turn, ILA V neurons provide input to the BMAp [ILA(V)→BMAp] (Supplementary Figure 1E-F). In addition, the BMAp is the recipient of input from AIp projection cells (AIp→BMAp) (Figure 5J) and BMAp neurons provide some input to the ORBm (BMAp→ORBm) (Supplementary Figure 6J).

#### Thalamic connections

Like the BLAp, BMAp neurons send weak projections to the MDm (BMAp→MDm) (Supplementary Figure 11G) and like the BLA.ac receives input from the rostral PVT (rPVT→BMAp) and PT (PT→BMAp) (Supplementary Figure 11D,I).

#### Connections to motor systems

##### Ventral striatum: ACB and OT

Like all other BLA projection neurons, those in BMAp target the ACB, in particular the medial region where input from BLA.ac, BLAp, and LA also terminate (BMAp→ACB medial) (Supplementary Figure 10A). BMAp neurons also sparsely project to medial OT, the same region that receives considerable input from the BLA.ac (BMAp→OT medial) (Figure 1F; Supplementary Figure 6D).

##### Ventral pallidum: substantia innominata (SI)

Projection neurons in the BMAp target the SI (BMAp→SI) and in turn the BMAp receives input from neurons located in different regions of the SI (SI→BMAp) (Supplementary Figure 10I-K).

##### BST and hypothalamic areas

Similar to the BLAp, neurons in the BMAp strongly project to BST, in particular to the anteromedial (BSTam) and principal (BSTpr) nuclei (BMAp→BSTam/BSTpr) (Figure 6A,D). As such, while BLAp neurons preferentially target nuclei in the lateral BST, BMAp neurons target nuclei in more medial BST (Figure 6B-D). BMAp neurons also send appreciable projections to many hypothalamic structures like the medial preoptic area (MPO), medial preoptic nucleus (MPN) (Figure 6K) (BMAp→MPO/MPN), the anterior (AHN) (Figure 6G), dorsomedial (DMH) (Figure 6F), ventromedial (VMH) (Figure 6M), and lateral (LHA) (Figure 6H) hypothalamic areas (BMAp→AHN/DMH/VMH/LHA).

##### Hippocampal formation

BMAp projection neurons appreciably target the deep layers of the ENTl [BMAp→ENTl(V/VI)], making them the only BLA projection neurons outside of the BLAv to provide strong input to ENTl (Supplementary Figure 8H). Like all other BLA nuclei, the BMAp receives input from the ENTl (Supplementary Figure 8H). Similar to BLA.ac and BLAp, BMAp neurons project to the stratum radiatum of ventral CA1 (CA1v_sr) and the SUBv, except BMAp neurons target specifically the molecular (SUBv_m) and stratum radiatum (SUBv_sr) layers of the SUBv, which distinguishes BMAp injections from those in the BLAa and BLAp (BMAp→CA1v_sr/SUBv_m/sr) (Figure 4D). Neurons in CA1v_sp and especially SUBv project back to the BMAp (CA1v_sp/SUBv→BMAp) (Figure 6N).

#### Olfactory connections

Although BMAp neurons do not target the PIR, similar to the BLA.al, BLAv, and LA, the BMAp receives strong input from the ventral PIR (PIRv→BMAp) (Supplementary Figure 9E). Of all nuclei, the BMAp is most strongly connected with other olfactory cortical areas like the piriform-amygdalar area (PAA), the cortical amygdalar areas (COA), and the piriform-amygdalar transition area (TR). Specifically, the BMAp receives input from the PAA (PAA→BMAp) (Supplementary Figure 13C) and from neurons in the anterior (COAa) (Supplementary Figure 13B), posterior lateral (COApl) (Supplementary Figure 13C), and posterior medial (COApm) (Supplementary Figure 13D) cortical amygdala areas (COAa/COApl/COApm→BMAp). Neurons in the BMAp project back to each of these areas (BMAp→COAa/COApl/COApm/PAA) (Supplementary Figure 13A-E). Finally, neurons in BMAp project to TR (BMAp→TR) and neurons in the TR project back to the BMAp (TR→BMAp) (Supplementary Figure 10G-H). Strongest BMAp connections with olfactory nuclei appear to be with the dorsal part of the endopiriform nucleus (EPd). BMAp projection neurons target EPd (BMAp→EPd) and in turn EPd projection neurons target the BMAp (EPd→BMAp) (Supplementary Figure 9H). Summarized brain-wide connections of projection neurons in BMAp is provided in a wiring diagram (Figure 6J). Connections of all BLA regions are summarized and presented in a global wiring diagram (Figure 9).

**Figure 9:**
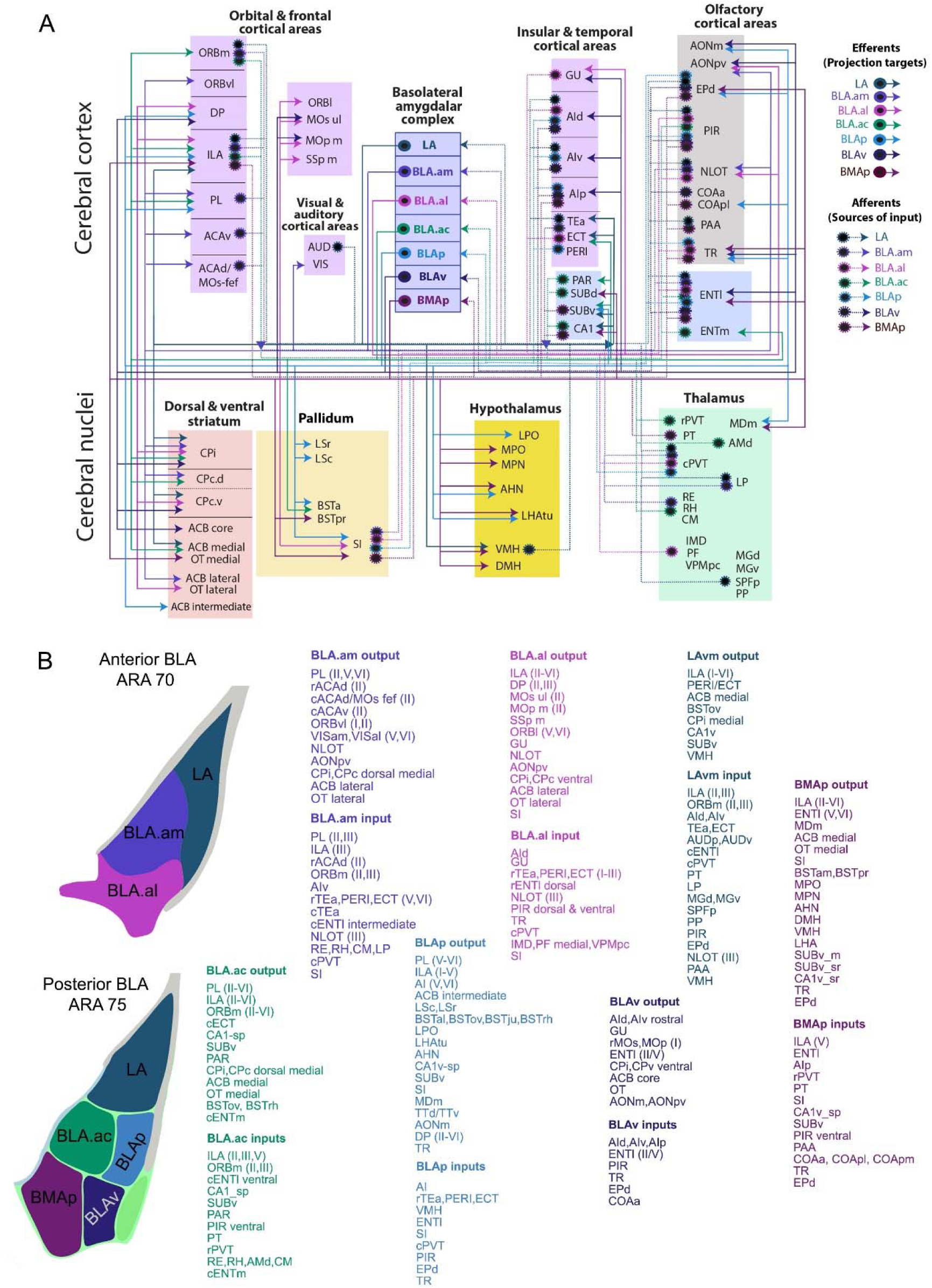
**A.** The brain-wide wiring diagram of the BLA complex. **B**. A summary of all input and output connections for BLA.am, BLA.al, BLA.ac, BLAp, BLAv, LA, and BMAp. For list of abbreviations, see Table 1.

### Functional characterization of synaptic innervation of projection-defined BLA neurons

Thus far, we showed uniquely connected neuron populations in all BLA nuclei. Such information lays the groundwork for future functional interrogations of these connectionally-defined BLA subtypes. We applied optogenetics and electrophysiological recordings to demonstrate several proof of principle examples. First, we further characterized the nature of synaptic connectivity between BLAam and PL and second we demonstrate a functional bisynaptic pathway through AId→BLA.al→CPc.ventral.

Coinjections of AAV-hSyn-ChR2-YFP (ChR2) and retrograde red microbeads were made into the PL, which delivered ChR2 to PL axonal terminals in the BLA.am (green fibers) and retrogradely labeled PL-projecting BLA.am neurons (red neurons) (Figure 7I). Consistent with the anatomical tracing results, YFP-labeled PL axons and terminals intermingled with red retrobeads labeled PL-projecting neurons in the BLAam were evident (Figure 7J). In brain slice preparations, whole-cell voltage-clamp recordings were made from fluorescence-labeled PL-projecting neurons, while PL axons in BLA were optically stimulated with blue light. As shown in Figure 7K, a 5-ms pulse of blue light elicited an excitatory current and a nearly concurrent inhibitory current in the recorded neuron clamped at −70 and 0 mV respectively when the slice was bathed in the normal external solution. In the presence of TTX (1 µM) and 4-AP (1 mM), which eliminate polysynaptic responses ^55, 56^, the inhibitory currents disappeared and the excitatory current was reduced in amplitude (Figure 7L, right panel), indicating that PL projections are excitatory and make monosynaptic innervation of PL projecting neurons in BLA. In addition, the differences in onset latencies between excitatory and inhibitory responses (Figure 7L right panel) recorded from the same neuron suggests bisynpatic feedforward inhibitory inputs, likely from the local inhibitory neurons innervated by the PL projections. Altogether, these results demonstrated that BLAam→PL projecting neurons were directly innervated by PL→BLAam projections, and also implied the existence of local feedforward inhibitory circuits for PL driving bisynaptic inhibition onto BLAam→PL projecting neurons (Figure 7L). The latter may function as a balanced control of activity in BLA. Next, we explored the synaptic connectivity between AId and BLA. Studies demonstrate that BLA affects motor behavior through its outputs to CEA ^57^. We show that the caudoputamen (CP) is one of the largest projection targets of the BLAa, which lead us to hypothesize that BLAa potentially integrates information from different cortical areas and in turn affects motor outputs through its projections to CP. Therefore, we tested to see whether we could demonstrate a functional AId→BLA.al→CPc.ventral connection, which could potentially be involved in gustatory/visceral information processing (see discussion). Accordingly, AAV-hSyn-ChR2-YFP was injected into AId, which generated dense projections to the BLAal. Meanwhile, red retrobeads were injected into CPc.ventral to retrogradely label BLA.al neurons (Figure 7M-N). Whole-cell voltage-clamp recordings were made from fluorescent-labeled CP-projecting BLA.al while AId axons in BLA were optically stimulated via blue light (Figure 7O). This demonstrated that BLA.al→CP projecting neurons are monosynaptically innervated by axonal inputs from AId. These results support a direct functional neural pathway of AId→BLA.al→CPc.ventral (Figure 7P). The existence of these neural pathways strongly suggest that BLAa may play an essential role in integrating top-down and bottom-up information and mediating the motor behaviors through its direct and indirect connections to CP. Guided by the anatomical data presented in this work, such functional connectivity studies can be performed for projection neurons across the BLA complex along with studies that directly test the behavioral output of the functional connections. Details of all methods are provided in the Online methods.

## DISCUSSION

We combined classic and viral tracing with various computation techniques like machine learning based image processing to delineate three domains within BLAa that house projection neurons with unique connectional profiles and morphological features. We provided the brain-wide connections of projection neurons located in all BLA nuclei including BLA.am (Figure 5B; Figure 7E; Supplemental Figure 13F), BLA.al (Figure 5G; Figure 7F; Supplemental Figure 13G), BLA.ac (Figure 4C; Figure 7G; Supplemental Figure 13H), BLAp (Figure 6I), BMAp (Figure 6J), BLAv (Figure 6O), and LA (Figure 7A). Consequently, we constructed a comprehensive interactive wiring diagram of the mouse BLA complex (Figure 9A; link will be provided). Data are presented online as open resources through our MCP website in various formats (www.MouseConnectome.org). Multi-fluorescent images of each amygdala case that are registered onto the standard ARA can be viewed through our iConnecome Viewer (link will be provided). The color-coded visualized output of the community assignments for all injections will also be presented online (http://www.mouseconnectome.org/amygdalar/) (currently password protected). These allow users to view all anterograde and retrograde connections within a common anatomical frame. Morphological reconstructions of BLA.am (Movie 1), BLA.al (Movie 2), and BLA.ac (Movies 3-5) projection neurons are also available online (<u>NeuroMorpho.org</u>) ^58^. Finally, we demonstrate that this foundational BLA roadmap can be used to functionally characterize synaptic innervation of connectionally unique BLA cell types. To our knowledge, this is the most comprehensive connectivity dataset from a single report in any mammalian BLA study. As reviewed in the Introduction, the BLA is implicated in a wide range of behaviors from fear to addiction, probably due to the rich variety of BLA neuron subtypes with complex connectional profiles. This data can be applied to interrogate the behavioral specificity of projection-defined BLA cell types. Some potential functional hypotheses based on the unique connectivity of BLA projection neurons are discussed below.

### BLA.am neurons and visual and spatial information

The BLA.am projection neurons target regions that process visual information and guide eye movement and orientation, including the deep layers of secondary visual areas (VISam, VISal), anterior cingulate cortex (ACAd and ACAv), MOs-fef, ORBvl ^35^, and dorsomedial division of the CP ^34^. In particular, ORBvl is in a cortical network of structures that process visual and spatial information ^35^. In turn, visual and spatial information reach the BLA.am from the caudal TEa, which receives abundant inputs from visual and auditory areas ^35^, from visual LP thalamic nucleus ^59^ and from the thalamic RE, which contains head direction cells ^60^. BLA connections to primary visual cortex are reported in primates ^61, 62^ although, to our knowledge, not in mice. The relevance of these connections are apparent in the context of primate ^63, 64^ and human ^65^ studies showing BLA involvement in the processing of facial expressions; however, they may also be relevant in rodents. Although BLA (BLAa, BLAp) neurons responsive to only visual cues are sparse in rats, neurons responsive to a variety of sensory modalities, including visual, auditory, somatosensory and gustatory stimuli, are dispersed throughout the BLA (BLAa, BLAp) ^66^. For example, although unimodal audition responsive neurons are localized in the lateral nucleus, multimodal auditory responsive neurons are mostly found in the basolateral and central amygdala. The same study showed that significantly more multimodal neurons that respond to stimuli previously paired in a conditioning task are located in the BLA (BLAa, BLAp) and the central nucleus ^66^. Our data suggests that multimodal neurons involving visual information may be located in the BLA.am. Since re-evaluation of stimuli for outcome prediction is one of the functional responsibilities of the BLA ^19, 67^, it is reasonable for BLA neurons to be equipped with the ability to survey and communicate about stimuli of various modalities, including those that are visual in nature.

### BLA.al neurons and gustatory information

The BLA.al has unique connections that suggest its role in a network of structures involved in gustatory processes. Largely distinct areas within the insular cortex (AId, AIv, AIp, and GU) are shown to code palatable and unpalatable tastants with neurons responding to sweet tastants located more rostrally than those that respond to bitter ^68–70^. Further, unique amygdalar projections from each of these “sweet” and “bitter” cortical areas to the basolateral and central nuclei respectively, are involved in processing the hedonic valence of the tastants ^71^. Our data show that although AI provides some input to BLA in general, the strongest projections appear to arise from rostral AId (the region that corresponds to the site of sweet responding neurons) to the BLA.al, corroborating the label patterns observed in more recent studies of the gustatory cortex ^71^. Projection cells targeting the AId and GU are also primarily located in the BLA.al. Further, BLA.al projection cells specifically target CP domains that receive convergent inputs from the AI, GU, and somatic and somatomotor mouth regions ^34, 72^, suggesting a role for BLA.al in feedback mechanisms potentially involving these regions. In fact, we showed that AId fibers directly innervate BLA.al projection neurons that target the CPc.ventral, validating the functional connection among these three regions. Further, based on its combined input/output connectivity to brain-wide structures, the BLA.al is most similar to the BLAv (http://www.mouseconnectome.org/mcp/docs/BLA/brain_wide_matrix.pdf) (currenty password protected), a region most likely involved in gustatory/visceral information processing as discussed below. Interestingly, BLA.al receives input from rostral regions of ECT and PERI, two cortical areas that share extensive bidirectional connectivity with somatic sensorimotor areas ^35^. Thalamic input to BLA.al comes from the caudal PVT, the IMD, medial part of PF, and VPMpc. The medial part of the thalamic PF nucleus was recently shown to project to AI ^73^, the caudal PVT receives input from the AI ^36^, and the VPMpc receives gustatory information from the medial parabrachial nucleus (PBN) within the gustatory taste pathway ^74^. Further the BLA.al shares reciprocal connections with IMD and BLAv, two potential sites of visceral/gustatory information processing (discussed below).

Recording data shows that cells responsive to multiple stimuli including oral sensory (e.g., taste), somatosensory, and auditory are located in the BLA and that cells that become responsive to these stimuli following their association with a reinforcement are preferentially located in the BLA ^66^. Further, a variety of taste neurons have been identified in BLA ^75^ and following odor-taste associations the number of BLA neurons that respond to both taste and odor dramatically increases ^76^. Given its connectivity architecture, BLA.al could be a candidate region in which these conditioned responsive cells involving taste stimuli are located. BLAp and BLAv are also possible locations.

### BLA.ac neurons and reinstatement of drug seeking behavior

The caudal part of the BLAa (BLA.ac) displays a connectivity fingerprint distinct from its rostral BLA.am and BLA.al counterparts. The most striking of these are its connections with hippocampal regions like SUBv, CA1v, but most particularly the CA3 and PAR. Connections with these latter two regions appear to be exclusive to the BLA.ac. Compared to the BLA.am and BLA.al, which provide input to the lateral parts of the ACB and OT ^33, 77–79^, projection cells in the BLA.ac provide input predominantly to their medial parts and in particular the medial shell of the ACB ^79^. Unique BLA.ac thalamic input is provided by the PT and rostral part of PVT ^39^. What could the specificity of BLA.ac neuronal connections potentially suggest about the information they process?

Reexposure to the environment in which drugs are self-administered can facilitate relapse following periods of abstinence. This contextually promoted vulnerability for relapse is modeled in laboratory animals using a paradigm called context-induced reinstatement. The BLA ^16–18^, medial shell of the ACB ^80–82^, SUBv ^83^, CA1v ^84^, CA3 ^84, 85^, PVT ^86^, and PL/ILA ^16, 87^ are all contributors to the circuitry that meditates the contextual reinstatement of drug seeking behaviors. Specifically, interactions between the BLA and medial ACB shell ^88^, PL/ILA ^89, 90^, and the PVT ^86^ are shown to be involved (for review see ^91^). Interestingly, similar to the medial shell of the ACB, the medial OT, and not lateral OT, is an important site for the reinforcing effects of psychostimulant drugs and self-administration ^92–94^ and it has been proposed that the medial OT is the ventral extension of the medial ACB ^92^. Not surprisingly, the potential contributions of a medial ACB+medial OT complex in context-induced reinstatement has been suggested ^81, 95, 96^. Connections of the BLA.ac with the PAR are in accord with the possibility of its role in contextual reinstatement given that the PAR is a primary site for spatial encoding, containing rich amounts of grid cells, head direction cells, and border cells ^97^. Within this context, input from the rostral PVT to the BLA.ac is also reasonable given that it is this division of the PVT that receives input from the subiculum, rather than the caudal PVT, which is influenced more by AI inputs ^36^. Of course, this is only one of many potential hypotheses regarding the BLA.ac suggested by its connectivity profile. Optical stimulation of projections from the BLA to ventral hippocampus are shown to be anxiogenic ^24^ and to reduce social interaction ^98^. The BLA.ac would be a likely candidate for exploring these particular circuits given its connections with the ventral hippocampus. Other candidates for this hypothesis would include the BLAp and BMAp.

### BLAv neurons and visceral/gustatory information

Another nucleus that is considered a part of the BLA, but has received little attention, is the ventral part of the BLA or BLAv, identified in mouse ^1^, but not rat ^99^ brain atlases. This nucleus appears to be in a network of regions associated with gustatory and visceral information including the AI, GU, IMD, ENTl, CP, and BLA.al. The most interesting finding regarding this nucleus is the striking similarity between its selective projection targets and those of the intermediodorsal nucleus of the thalamus (IMD). The IMD is grouped into the dorsal midline thalamic nuclei speculated to be involved in viscero-limbic functions ^38^. These shared targets include the AI, ventral medial parts of the intermediate CP, the ventral medial part of the caudal CP, and a very distinct region in layer V of the ENTl ^100, 101^. The ventral parts of the CPi is a region where inputs from AI, GU, VISC, and somatomotor regions associated with mouth regions converge ^34^. The ventral part of the caudal CP is a region that receives densest input from the VISC ^34^, but also from GU, primary motor cortex associated with mouth regions, and from thalamic CM, a region known for its dense projections to GU and VISC cortices ^38^. This restricted region of the ENTl(V) also receives strong input from the AId and AIp. Finally, strong input to BLAv is provided by AI and PIR.

The potential for a BLA.al influence within this network is also plausible given that (1) the BLA.al projects to the same CP regions as the BLAv and IMD, (2) it receives input from the IMD, (3) receives input from the restricted ENTl(V) region that is connected with the IMD, BLAv, AI, GU, and VISC, and (4) the BLA.al is directly connected with the BLAv.

### The BLA and emotional conditioning

The BLA is most recognized for its involvement in fear conditioning, the archetypal paradigm used to study acquisition and retention of fearful memories. This conditioned emotional response and its resistance to extinction mimic the persistent pathological fears that manifest in human post-traumatic stress disorder (PTSD) ^102–104^. The BLA, ILA, PL, and hippocampus (HPF) have all been implicated in fear conditioning. BLA is implicated in the acquisition, consolidation, and retrieval of extinction memory ^2–13^. Further, distinct basal amygdalar cell populations connected with either the HPF or medial prefrontal areas (MPF) are shown to be activated under either conditions of fear or extinction, respectively ^3^. Therefore, the intricate connections among the BLA, MPF, and HPF can guide hypotheses regarding the individual roles of the subregional neuron connections in the fear acquisition/extinction circuits.

The HPF is shown to contextually modulate extinction ^105, 106^ and it is suggested that basal amygdalar nuclei serve as the primary site of contextual input to the amygdala ^107, 108^. So where in the basal amygdala could this hippocampal contextual information be received? The HPF is connected with neurons in BLAp and BMAp through CA1v and SUBv, but also with neurons in BLA.ac. Aside from the CA1 (dorsal and ventral sp layers), CA3v, and SUBv, the BLA.ac neurons are strongly connected with PAR, a primary site for spatial encoding that contains rich amounts of grid cells, head direction cells, and border cells ^97^, all ideal information for contextual information processing.

The ILA is predominantly involved in fear extinction consolidation and recall, while the PL is implicated in persistent fear expression, resistance to extinction, and contextual renewal ^12, 109–115^. Immediate early gene studies corroborate this inverse relationship of the ILA and PL ^116^. Indeed, PL projecting BLA are activated during fear expression, while ILA projecting BLA neurons show increased activity when extinction memory is expressed ^23^. Our data show that ILA targeting BLA neurons are primarily located in BLA.al, BLA.ac, BLAp, BMAp, and LA. The laminar specificity of these projections are also demonstrated. PL projecting neurons primarily are located in BLA.am and BLA.ac, which target layer II, and in the BLAp, which project to deeper PL layers.

Input from PL to BLA are strongest to the BLA.am. Most rat anatomy literature has demonstrated strong ILA→LA ^31, 117^ projections and although here we show strong projections from ILA to LA, the strongest input from ILA to BLA are to BLA.ac and BMAp.

The ACA is also proposed to be involved in extinction of contextual fear learning ^118^. Of all BLA neurons, those in BLA.am appear to have the strongest connections with the ACA. BLA.am neurons project strongly to both ACAd and ACAv, while neurons in the more anterior regions of ACAd and ACAv provide input to BLA.am.

### BLA, orbitofrontal connections, and representations of value

Extensive research has implicated the BLA in processes that require an animal to update the representations of a CS value to make appropriate decisions, like in the case of fear extinction ^119, 120^. Both rats ^121–123^ and monkeys ^124, 125^ with BLA lesions are unable to adjust their behavior accordingly upon updating of the reinforcer value. This is the case whether the reinforcers are devalued or inflated ^126^ suggesting that the BLA tracks a state value ^120^. That is, animals with loss of BLA function are unable to accurately predict the rewarding or punishing outcome of a reinforcer that enables the selection of the more advantageous behavior. It is speculated that the connections of the BLA with the lateral part of the orbitofrontal cortex (ORBl) are relevant for the appropriate assessment of expected outcomes and consequently the selection of appropriate action ^127–129^, a reasonable expectation given that the ORB is speculated to be involved in goal-directed behaviors and its dysfunction renders patients unable to make proper judgments based on anticipated consequences ^130–132^. Tract tracing studies in rats report strong projections from BLA to ORB, typically the ORBl and ORBvl ^133, 134^. Our data show that BLA projection neurons targeting the ORBvl(I/II) reside in the BLA.am and those that target the ORBl(V/VI) are in the BLA.al. Projections from ORBvl and ORBl back to BLA were not detected ^54^ although BLA bound ORB neurons were found in the medial part of the ORB (ORBm), which target mainly BLA.am and BLA.ac.

### BLAp, BMAp and anxiety

The BLA→BST pathway is shown to regulate anxiety. Specifically, BLA targets to the oval nucleus of the BST are shown to be anxiogenic, while optogenetic stimulation of BLA inputs to the anterodorsal BST, which include the BSTam and BSTal, promote anxiolysis ^22^. Our data provide insight into the location of these projectionally and functionally disparate cell types. Some BLA projection neurons giving rise to anterodorsal BST input reside mostly in BLAp and BMAp, and while BMAp neurons largely avoid the BST oval, BLAp neurons densely innervate it. Further, it has been shown that activating projections from the ventromedial prefrontal cortex (vmPFC) to the BMAp reduces anxiety and enhances extinction of acquired cued and contextual fear ^135^. BMAp projecting neurons in the vmPFC were shown to be located specifically in layer V. Our data validate this and show that these vmPFC projections are from layer V of the ILA. Our data also show that hippocampal contextual information can reach the BMAp via input from the SUBv ^136–138^.

## ONLINE METHODS

All data were produced as part of the Mouse Connectome Project (MCP, www.MouseConnectome.org) at the Center for Integrative Connectomics (CIC) at USC.

### Subjects

Data from 8-week old male C57BL/6J mice (n=153; Jackson Laboratories) were used to trace all pathways reported in the current report. Animals were housed in pairs in a room that was temperature (21-22 °C), humidity (51%), and light controlled (12-hr light:12-hr dark cycle). The mice were allowed at least 1 week to adapt to their living conditions prior to stereotaxic surgeries for the delivery of tracers. Subjects had ad libitum access to tap water and mouse chow throughout the experiments. All experiments were conducted according to the regulatory standards set by the National Institutes of Health Guide for the Care and Use of Laboratory Animals and by the institutional guidelines set by the Institutional Animal Care and Use Committee at USC.

### Surgical methods

Experimental procedures for tracer injections have been described previously ^34, 35^. Anterograde and retrograde tracers were delivered to anatomically delineated regions across the brain to assess their connectivity patterns. Surgeries for tracer infusions were performed under isoflurane anesthesia (Hospira, Inc.). Mice were initially anesthetized in an induction chamber primed with isoflurane and were subsequently mounted to the stereotaxic apparatus where they were maintained under anesthetic state via a vaporizer (Datex-Ohmeda). The isoflurane was vaporized and mixed with oxygen (0.5 L/min) and nitrogen (1 L/min). The percent of isoflurane in the gas mixture was maintained between 2 and 2.5 throughout the surgery. Tracers were delivered iontophoretically using glass micropipettes whose outside tip diameters measured approximately 10-30 µm. A positive 5 µAmp, 7-second alternating injection current was delivered for 10 minutes (Stoelting Co.).

### Tracing strategies

Anterograde tracers included *Phaseolus vulgaris* leucoagglutinin (Phal; 2.5%; Vector Laboratories) and adeno-associated viruses encoding enhanced green fluorescent protein (AAV GFP; AAV2/1.hSynapsin.EGFP.WPRE.bGH; Penn Vector Core) or tdTomato (AAV RFP; AAV1.CAG.tdtomato.WPRE.SV40; Penn Vector Core). Retrograde tracers included cholera toxin subunit b conjugates 647 and 555 (CTb; AlexaFluor conjugates, 0.25%; Invitrogen), Fluorogold (FG; 1%; Fluorochrome, LLC), and AAVretro-EF1a-Cre (AAV retro Cre; Viral Vector Core; Salk Institute for Biological Studies). Anterograde and retrograde tracers were injected either in combination (e.g., coinjection of Phal with CTb 647) or individually in a triple anterograde (Phal, AAV-GFP, AAV-RFP) or a quadruple retrograde (CTb 647, CTb 555, FG, AAV retro Cre) injection design.

To trace the connectivity of projection defined neurons, a Cre-dependent anterograde tracing strategy was employed, which involved the delivery of Cre to projection neurons of interest via an injection of AAVretro-EF1a-Cre into their target regions. Next, AAV1.CAG.FLEX.eGFP.WPRE.bGH and AAV1.CAG.FLEX.TdTomato.WPRE.bGH (Addgene; plasmid ID #s 51502, 51503) were delivered to the different Cre-expressing BLA neuronal populations. Finally, to retrieve the morphological information from different amygdalar projection neurons, Gdel-RV-4tdTomato ^27^ and Gdel-RV-4eGFP ^28^ injections were made into two different downstream targets of the neurons.

### Tissue processing and imaging in 2D

Either one (Phal, CTb, FG, AAV retro Cre, Gdel-RV) or three (Cre- and non-Cre-dependent AAVs) weeks was allowed for tracer transport after which animals were perfused and their brains were extracted. For all but the morphology studies involving Gdel-RV injections, a 2D tissue processing workflow was followed. After an overdose injection of sodium pentobarbital, each animal was transcardially perfused with approximately 50 ml of 0.9% NaCl followed by 50 ml of 4% paraformaldehyde solution (PFA; pH 9.5). The brains were post-fixed in 4% PFA for 24-48 hours at 4°C after which they were embedded in 3% Type I-B agarose (Sigma-Aldrich) prior to sectioning. Four series of coronal sections were sliced at 50-µm thickness with a Compresstome (VF-700, Precisionary Instruments, Greenville, NC) and prepared for immunofluorescence staining.

One series of sections was immunostained for the antigen of interest (Phal or AAVretro-EF1a-Cre) using the free-floating method (see Supplemental methods for details). All sections were counterstained with a fluorescent Nissl stain (NeuroTrace® 435/455; Invitrogen, #N21479) and then were mounted and coverslipped using 65% glycerol. Sections were scanned at 10x magnification as high-resolution virtual slide image (VSI) files using an Olympus VS120 high-throughput microscope using identical exposure parameters across all cases.

### Post image acquisition data processing

Sections from each case were assigned and registered to a standard set of 32 corresponding Allen Reference Atlas levels ranging from 25-103 (see Supplemental methods for details). Threshold parameters were individually adjusted for each case and tracer. Conspicuous artifacts in the threshold output were corrected in Photoshop. Oversaturation at the injection site prevents the detection of labeled fibers and boutons at the injection sites (BLAa) and their surrounding areas (LA, CEA). Therefore, with this injection strategy, it is difficult to assess intra-amygdalar connections especially within the BLA complex and the CEA. Due to proximity to the injections sites, these areas appear as dense label in the threshold output subsequently affecting annotation and analysis. To obviate this, these areas of potentially false dense label at or surrounding the injection sites (specifically the BLA complex and CEA) were filtered out at the thresholding stage for all BLAa cases.

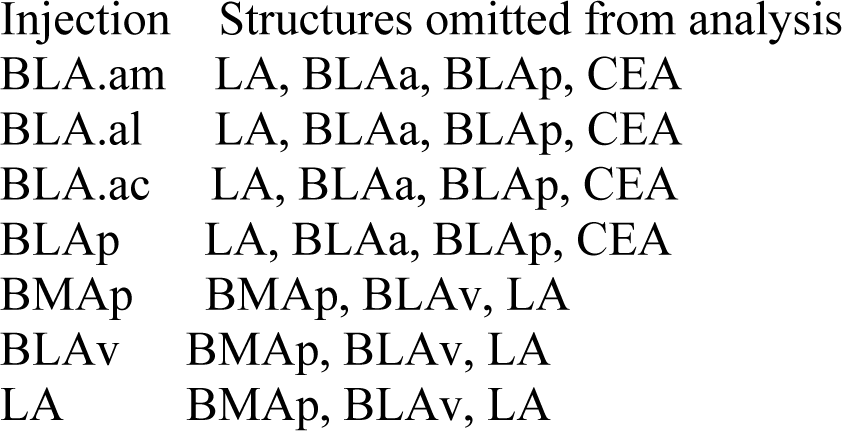

### Data analysis and visualization

#### Injection site analysis

Some injection sites were rescanned under lower exposure parameters to acquire a more accurate view of the size and location of the injections. Sections containing the injections for each case were re-registered to ARA levels that best corresponded to the target BLA complex structures. For injections that spanned more than a single section, each section was registered and analyzed. An injection site annotation algorithm (Supplemental methods) was run on the sections to identify injection location. The annotation was run atop a custom ARA atlas that contained the manually delineated boundaries of the BLAa domains.

#### Injection site validation

Importantly, injections reported for the BLAa domains are not entirely confined. For instance, the BLA.al AAV-RFP injection encroached into the BLA.am and BLAp (Supplementary Figure 4A). Consequently, although the BLA.al AAV-RFP injection largely represents BLA.al projections, it also shows output more specific to BLA.am and BLAp. Similarly, the BLA.al FG injection spread into the LA and BLA.am showing labeling patterns of mostly BLA.al, but also of BLA.am and LA. Tracer spread is expected given the small size of the domains and their close proximity to one another. In fact, infusion spread across ROIs is a valid and pervasive concern for all neuroanatomical studies. Several measures were taken to mitigate this. First, injection sites that were mostly confined to a single domain were selected for analysis so that the label would primarily represent connections of the target domain both in terms of presence and intensity of label. This strategy, combined with the grouping of the three domains for community detection analysis, which assigned grids to injection sites with the greatest pixel intensity value, helped to visualize labeling originating most likely from the target domain. Second, each of the injections was repeated in at least two cases for verification purposes. Third, only tracing data that was validated is reported. Anterograde labels were validated with retrograde tracers, while retrograde labels were validated with anterograde tracers. For example, in the BLA.al FG injection above, back-labeled cells were observed in rostral ACAd. However, an AAV injection in the ACAd primarily labeled the BLA.am, suggesting that the ACAd labels were from tracer spread into the BLA.am (Supplementary Figure 4D). In the case of the BLA.al AAV-RFP injection, the MDm and PL were labeled. However, retrograde injections placed in these two regions showed that the MDm label most likely originates from BLAp, while the PL label most likely resulted from tracer spread into the BLA.am (Supplementary Figure 4E). Further, the PL label from the BLA.al was most evident in the analysis in which this domain was paired with the BLAp. However, once the BLA.al was grouped with the BLA.am and BLA.ac for community detection analysis, projections to the PL got assigned to injection sites in the BLA.am or BLA.ac since injections in these two domains resulted in far denser labels in the PL than the injection in the BLA.al. Finally, Cre-dependent anterograde tracing also was utilized to parse out connections unique to the individual domains (Supplementary Figure 5).

#### Machine-learning assisted BLAa boundary demarcation

To determine BLAa domain characteristics and boundaries, we first identified representative cases demonstrating selective labeling in each domain. For each case, we histologically prepared all consecutive sections across the BLA and imaged the sections with an Olympus VS120 virtual slide microscope. Sections were then processed through our in-house image processing software Connection Lens. Briefly, each section was matched and warped to their corresponding atlas level of the Allen Reference Atlas (ARA) and the labeling was segmented and the data were used as a training set for a machine learning algorithm (see Supplemental methods for details).

#### Community detection

We performed community detection (modularity maximization) on both grid and ROI annotated data. As a first step, the overlap annotation per each group was aggregated into a single matrix. We set a minimum threshold value of 0.0045 for labeled pixels (anterograde) and 8 for labeled cells (retrograde), and removed entries that did not meet the threshold. These minimum values were set to exclude potential false positives, but also to exclude extremely light connections that would be difficult to explain (e.g., 2 labeled cells or a few labeled fibers in an ROI). Once the aggregated matrix was constructed, we further normalized so that the total labelling across each injection site (typically close in the first place) was adjusted to equal with the injection site featuring maximum total labeling. On this normalized matrix we applied the Louvain ^139^ algorithm at a single scale (gamma 1.0). As the result of this greedy algorithm is non-deterministic, we performed 100 separate executions, and subsequently calculated a consensus community structure ^140^ to characterize the 100 executions as a single result.

#### Connectivity matrices, community color coding and data visualization

For matrix visualization, we applied the grid_communities algorithm to modularize ROIs based on community assignment, as well as prioritize connections along the diagonal. Such a visualization provides a high level overview of the connectivity (Figure 3H-I; Figure 4F-G).

Community coloring provides more detail (i.e. drill down from the matrix overview) by visually encoding segmentation by community, and ultimately, injection site (Figure 1A, anterograde and retrograde visuals). To carry out this process, we developed software to programmatically march through each segmented pixel per each image. Using either the grid location or ROI name, the algorithm looked up the corresponding community assigned during the consensus community step. Using a table containing a color assigned (by the authors) to each injection site, the algorithm retrieved the injection site associated with the community, and colored the pixel with the corresponding injection site color value. A subsequent step took advantage of the fact that each pixel was assigned to only a single community by aggregating all colorized images corresponding to a given atlas level into a single representative image. See Supplemental methods for details on the generation of proportional stacked bar charts.

### 3D workflow for assessment of neuronal morphology

To assess whether projection neurons located in the 3 domains of the BLAa were morphologically distinct, representative neurons in each of the domains were labeled via a G-deleted rabies injection in the caudal dorsomedial caudoputamen (for BLA.am), caudal ventral caudoputamen (for BLA.al), and in the medial accumbens (for BLA.ac) (Figure 1D). One week was allowed for tracer transport following injections, after which the animals were perfused. A 3D tissue processing workflow was followed for implementation of the SHIELD clearing protocol ^49, 50^. See Supplementary methods for details on the SHIELD clearing method and for our 3D imaging protocol.

#### 3D reconstructions, visualization, and analysis of neuronal morphology

Manual reconstruction of the neurons was performed using Aivia (version.8.5, DRVision) (Figure 8D) (Movies 3-5), and geometric processing of neuron models was performed using the Quantitative Imaging Toolkit (QIT) ^141^ (also available at http://cabeen.io/qitwiki). Although CPc.dorsomedial projecting BLAa neurons primarily are located in BLA.am, some are also located in BLA.al and BLA.ac. As such, rabies injections in the CPc.dorsomedial label mostly BLA.am, but also some BLA.al and BLA.ac neurons. To ensure the traced neurons in fact solely originated from their respective domains, for all cases, labeled cells visually grouped in the center of the boundaries of each domain (X-Y axes) were traced. Neurons located within the BLA.am were identified for inclusion in the analysis, but those close to the BLA.al or BLA.ac border were excluded. Similarly, neurons within BLA.ac were identified for inclusion, but those close to the border of the BLAp were omitted. In addition, domain-specific neurons in deeper parts of the tissue (z axis) farthest away from the edges of the sectioned tissue were selected for reconstruction. To assist in the accurate selection of neurons, maximum intensity projections of 25 µm were created that aided in the identification of the BLAa domain boundaries. In total, eight neurons were traced for the BLA.am, nine for the BLA.al, and six for the BLA.ac.

Reconstructions of individual neurons posed a challenge due to dense labeling in the BLA.am and BLA.al regions (Figure 1D) (Movies 1-2). All labeled neurons in the BLA.ac contralateral to the injection site were also traced (n=3; Figure 8C) (Movies 4-5). The ipsilateral neurons were embedded within densely labeled clusters, while the contralateral neurons were relatively sparser; thus, these contralateral neuron provided more exact reconstructions for assessing the accuracy of those on the ipsilateral side (video link to be provided). Statistical comparisons between the ipsilateral and contralateral BLA.ac neurons were also run on all previously described measured morphological features (discussed below).

Due to the differences in slice thickness (250 µm for BLA.am and BLA.al and 400 µm for BLA.ac), we restricted our morphological analysis to neurites that were sufficiently close to the soma. We accomplished this by applying the NeuronTransform module in QIT to trim the contiguous portion of neurites that measured farther than 300 nm away from the center of the soma using the Euclidean distance (Supplementary Figure 1G). Due to the anisotropic dimensions of the voxels and spatial undersampling relative to the curvature of the dendrite, we applied a local regression filter to address the aliasing artifact and to regularize dendritic tortuosity. Specifically, the NeuronFilter module in QIT was used to apply a locally weighted scatter-plot smoother (LOESS), which is a low bias approach that makes minimal assumptions (Supplementary Figure 1H) ^142^. LOESS was implemented using a locally quadratic function with the five nearest neighbor neuron vertices before and after each point of estimation endpoints and bifurcation points were excluded from filtering.

Importantly, CP-projecting BLA.am/BLA.al neurons and ACB-projecting BLA.ac neurons were selected as representative domain specific BLA→striatum neurons and are not representative of the entire population of neurons within the 3 domains. It is possible that MPF projecting BLAa neurons display morphological features that differ from BLAa→MPF or BLAa→AI neurons. Such hypotheses can be investigated in the future.

#### Statistical analysis

To obtain an overall view of the dendritic morphology of BLA projection neurons located in the BLA.am, BLA.al, and BLA.ac, we applied the classic ^143^ and modified Sholl ^144^ analysis using Fiji ImageJ and L-Measure, respectively. Quantitative morphological parameters depicting the somas and dendrites were obtained from Aivia and statistical analyses were performed using the R computing environment (RStudio Version 1.1.463). Standard measurements representing cell body morphology included volume, sphericity, spherical diameter, X, Y, Z and Euclidean skewness, cell body height, width, depth, and surface to volume ratio. Measurements of dendritic morphology included number of primary dendrites, branches, bifurcations, terminal tips, nodes, fractal dimension, tortuosity, local bifurcation angle, remote bifurcation angle, partition asymmetry, tips to dendrite ratio, branch generation, and Rall’s ratio. Using all of the measured morphological parameters principal component analysis (PCA) was run to reduce the dimensionality and create a 3D scatterplot ^51, 52^. The PCA shows the segregation of BLAa domain specific neurons based on the measured features (Figure 8B).

Two-sided pairwise Wilcoxon rank sum tests were performed, and the parameters that survived FDR correction for multiple testing with a q-value <0.05 are reported. The significant group differences are presented with whisker plots (Figure 8F), and the degree of their significance is visualized in a matrix plot (Figure 8E). See Supplemental methods for details regarding output of these statistical analyses.

In a final analysis, we characterized our data using an alternative tool that uses topological data analysis that retains potentially more morphological information. Specifically, we used the persistence-based neuronal feature vectorization framework ^145^ to summarize pairwise distances between neurons ^146^. Our experiments used code provided by the authors online, where we first computed persistence diagrams using NeuronTools (https://github.com/Nevermore520/NeuronTools) and then computed inter-neuron distances using the Wasserstein metric (https://bitbucket.org/grey_narn/geom_matching/src). The results were visualized in matrix plot created using R (Figure 8G-H).

### Slice recordings

Three weeks following the viral injections of AAV-hSyn-ChR2-YFP (UPenn Vector Core) and red retrograde microbeads (Lumafluor Inc.), animals were decapitated following isoflurane anesthesia and the brain was rapidly removed and immersed in an ice-cold dissection buffer (composition: 60 mM NaCl, 3 mM KCl, 1.25 mM NaH2PO4, 25 mM NaHCO3, 115 mM sucrose, 10 mM glucose, 7 mM MgCl2, 0.5 mM CaCl2; saturated with 95% O2 and 5% CO2; pH = 7.4). Coronal slices at 350 μm thickness were sectioned by a vibrating microtome (Leica VT1000s), and recovered for 30 min in a submersion chamber filled with warmed (35°C) ACSF (composition:119 mM NaCl, 26.2 mM NaHCO3, 11 mM glucose, 2.5 mM KCl, 2 mM CaCl2, 2 mM MgCl2, and 1.2 NaH2PO4, 2 mM Sodium Pyruvate, 0.5 mM VC). BLA neurons (labeled with fluorescent microbeads) surrounded by EYFP+ fibers were visualized under a fluorescence microscope (Olympus BX51 WI). Patch pipettes (∼4–5 MΩ resistance) filled with a cesium-based internal solution (composition: 125 mM cesium gluconate, 5 mM TEA-Cl, 2 mM NaCl, 2 mM CsCl, 10 mM HEPES, 10 mM EGTA, 4 mM ATP, 0.3 mM GTP, and 10 mM phosphocreatine; pH = 7.25; 290 mOsm) were used for whole-cell recordings. Signals were recorded with an Axopatch 700B amplifier (Molecular Devices) under voltage clamp mode at a holding voltage of –70 mV for excitatory currents, filtered at 2 kHz and sampled at 10 kHz. Tetrodotoxin (TTX, 1 μM) and 4-aminopyridine (4-AP, 1 mM) were added to the external solution for recording monosynaptic responses to blue light stimulation (5 ms pulse, 3 mW power, 10–30 trials).

## ACKNOWLEDGEMENTS

This work was supported by NIH/NIMH U01MH114829 (HWD), NIH/NIMH R01MH094360 (HWD), and NIH/NIMH U19MH114821 (JH/PA).

## COMPETING FINANCIAL INTERESTS

The authors declare no competing financial interests.

All links and movies will be available upon publication of the manuscript.

## References

1. Dong, H.W. The Allen reference atlas: A digital color brain atlas of the C57Bl/6J male mouse (John Wiley & Sons Inc, 2008).

2. Falls, W.A., Miserendino, M.J. & Davis, M. Extinction of fear-potentiated startle: blockade by infusion of an NMDA antagonist into the amygdala. The Journal of neuroscience : the official journal of the Society for Neuroscience 12, 854–863 (1992).

3. Herry, C., et al. Switching on and off fear by distinct neuronal circuits. Nature 454, 600–606 (2008).

4. Herry, C. & Mons, N. Resistance to extinction is associated with impaired immediate early gene induction in medial prefrontal cortex and amygdala. The European journal of neuroscience 20, 781–790 (2004).

5. Laurent, V. & Westbrook, R.F. Distinct contributions of the basolateral amygdala and the medial prefrontal cortex to learning and relearning extinction of context conditioned fear. Learning & memory (Cold Spring Harbor, N.Y.) 15, 657–666 (2008).

6. Lin, C.H., Lee, C.C. & Gean, P.W. Involvement of a calcineurin cascade in amygdala depotentiation and quenching of fear memory. Molecular pharmacology 63, 44–52 (2003).

7. Lin, C.H., Yeh, S.H., Lu, H.Y. & Gean, P.W. The similarities and diversities of signal pathways leading to consolidation of conditioning and consolidation of extinction of fear memory. The Journal of neuroscience : the official journal of the Society for Neuroscience 23, 8310–8317 (2003).

8. Lu, K.T., Walker, D.L. & Davis, M. Mitogen-activated protein kinase cascade in the basolateral nucleus of amygdala is involved in extinction of fear-potentiated startle. The Journal of neuroscience : the official journal of the Society for Neuroscience 21, Rc162 (2001).

9. Maren, S. Seeking a spotless mind: extinction, deconsolidation, and erasure of fear memory. Neuron 70, 830–845 (2011).

10. Myers, K.M. & Davis, M. Behavioral and neural analysis of extinction. Neuron 36, 567–584 (2002).

11. Quirk, G.J. & Mueller, D. Neural mechanisms of extinction learning and retrieval. Neuropsychopharmacology : official publication of the American College of Neuropsychopharmacology 33, 56–72 (2008).

12. Sierra-Mercado, D., Padilla-Coreano, N. & Quirk, G.J. Dissociable roles of prelimbic and infralimbic cortices, ventral hippocampus, and basolateral amygdala in the expression and extinction of conditioned fear. Neuropsychopharmacology : official publication of the American College of Neuropsychopharmacology 36, 529–538 (2011).

13. Zimmerman, J.M. & Maren, S. NMDA receptor antagonism in the basolateral but not central amygdala blocks the extinction of Pavlovian fear conditioning in rats. The European journal of neuroscience 31, 1664–1670 (2010).

14. Tye, K.M., et al. Amygdala circuitry mediating reversible and bidirectional control of anxiety. Nature 471, 358–362 (2011).

15. Sharp, B.M. Basolateral amygdala and stress-induced hyperexcitability affect motivated behaviors and addiction. Translational psychiatry 7, e1194 (2017).

16. Fuchs, R.A., et al. The role of the dorsomedial prefrontal cortex, basolateral amygdala, and dorsal hippocampus in contextual reinstatement of cocaine seeking in rats. Neuropsychopharmacology : official publication of the American College of Neuropsychopharmacology 30, 296–309 (2005).

17. Marinelli, P.W., Funk, D., Juzytsch, W. & Le, A.D. Opioid receptors in the basolateral amygdala but not dorsal hippocampus mediate context-induced alcohol seeking. Behavioural brain research 211, 58–63 (2010).

18. Neisewander, J.L., et al. Fos protein expression and cocaine-seeking behavior in rats after exposure to a cocaine self-administration environment. The Journal of neuroscience : the official journal of the Society for Neuroscience 20, 798–805 (2000).

19. Corbit, L.H. & Balleine, B.W. Double dissociation of basolateral and central amygdala lesions on the general and outcome-specific forms of pavlovian-instrumental transfer. The Journal of neuroscience : the official journal of the Society for Neuroscience 25, 962–970 (2005).

20. Beyeler, A., et al. Organization of valence-encoding and projection-defined neurons in the basolateral amygdala. Cell reports 22, 905–918 (2018).

21. McGarry, L.M. & Carter, A.G. Prefrontal cortex drives distinct projection neurons in the basolateral amygdala. Cell reports 21, 1426–1433 (2017).

22. Kim, S.Y., et al. Diverging neural pathways assemble a behavioural state from separable features in anxiety. Nature 496, 219–223 (2013).

23. Senn, V., et al. Long-range connectivity defines behavioral specificity of amygdala neurons. Neuron 81, 428–437 (2014).

24. Felix-Ortiz, A.C., et al. BLA to vHPC inputs modulate anxiety-related behaviors. Neuron 79, 658–664 (2013).

25. Janak, P.H. & Tye, K.M. From circuits to behaviour in the amygdala. Nature 517, 284 (2015).

26. Pitkanen, A. Connectivity of the rat amygdaloid complex. in The amygdala (ed. J.P. Aggleton) 31–116 (Oxford University Press, 2000).

27. Chatterjee, S., et al. Nontoxic, double-deletion-mutant rabies viral vectors for retrograde targeting of projection neurons. Nature neuroscience 21, 638 (2018).

28. Wickersham, I.R., Sullivan, H.A. & Seung, H.S. Production of glycoprotein-deleted rabies viruses for monosynaptic tracing and high-level gene expression in neurons. Nature protocols 5, 595 (2010).

29. Little, J.P. & Carter, A.G. Synaptic mechanisms underlying strong reciprocal connectivity between the medial prefrontal cortex and basolateral amygdala. Journal of Neuroscience 33, 15333–15342 (2013).

30. McDonald, A.J. Organization of amygdaloid projections to the mediodorsal thalamus and prefrontal cortex: a fluorescence retrograde transport study in the rat. Journal of Comparative Neurology 262, 46–58 (1987).

31. McDonald, A.J., Mascagni, F. & Guo, L. Projections of the medial and lateral prefrontal cortices to the amygdala: a Phaseolus vulgaris leucoagglutinin study in the rat. Neuroscience 71, 55–75 (1996).

32. Sesack, S.R., Deutch, A.Y., Roth, R.H. & Bunney, B.S. Topographical organization of the efferent projections of the medial prefrontal cortex in the rat: an anterograde tract-tracing study with Phaseolus vulgaris leucoagglutinin. Journal of Comparative Neurology 290, 213–242 (1989).

33. Bienkowski, M.S., et al. Integration of gene expression and brain-wide connectivity reveals the multiscale organization of mouse hippocampal networks. Nature neuroscience 21, 1628 (2018).

34. Hintiryan, H., et al. The mouse cortico-striatal projectome. Nature neuroscience 19, 1100–1114 (2016).

35. Zingg, B., et al. Neural networks of the mouse neocortex. Cell 156, 1096–1111 (2014).

36. Li, S. & Kirouac, G.J. Sources of inputs to the anterior and posterior aspects of the paraventricular nucleus of the thalamus. Brain structure & function 217, 257–273 (2012).

37. Moga, M.M., Weis, R.P. & Moore, R.Y. Efferent projections of the paraventricular thalamic nucleus in the rat. The Journal of comparative neurology 359, 221–238 (1995).

38. Van der Werf, Y.D., Witter, M.P. & Groenewegen, H.J. The intralaminar and midline nuclei of the thalamus. Anatomical and functional evidence for participation in processes of arousal and awareness. Brain research. Brain research reviews 39, 107–140 (2002).

39. Vertes, R.P. & Hoover, W.B. Projections of the paraventricular and paratenial nuclei of the dorsal midline thalamus in the rat. The Journal of comparative neurology 508, 212–237 (2008).

40. Bhatnagar, S., Huber, R., Nowak, N. & Trotter, P. Lesions of the posterior paraventricular thalamus block habituation of hypothalamic-pituitary-adrenal responses to repeated restraint. Journal of neuroendocrinology 14, 403–410 (2002).

41. Heydendael, W., et al. Orexins/hypocretins act in the posterior paraventricular thalamic nucleus during repeated stress to regulate facilitation to novel stress. Endocrinology 152, 4738–4752 (2011).

42. Salazar-Juarez, A., Escobar, C. & Aguilar-Roblero, R. Anterior paraventricular thalamus modulates light-induced phase shifts in circadian rhythmicity in rats. American journal of physiology. Regulatory, integrative and comparative physiology 283, R897–904 (2002).

43. Li, Y., Dong, X., Li, S. & Kirouac, G.J. Lesions of the posterior paraventricular nucleus of the thalamus attenuate fear expression. Frontiers in behavioral neuroscience 8, 94 (2014).

44. Penzo, M.A., et al. The paraventricular thalamus controls a central amygdala fear circuit. Nature 519, 455–459 (2015).

45. Barson, J.R. & Leibowitz, S.F. GABA-induced inactivation of dorsal midline thalamic subregions has distinct effects on emotional behaviors. Neuroscience letters 609, 92–96 (2015).

46. Li, Y., et al. Orexins in the paraventricular nucleus of the thalamus mediate anxiety-like responses in rats. Psychopharmacology 212, 251–265 (2010).

47. Barson, J.R., Ho, H.T. & Leibowitz, S.F. Anterior thalamic paraventricular nucleus is involved in intermittent access ethanol drinking: role of orexin receptor 2. Addiction biology 20, 469–481 (2015).

48. Swanson, L.W. Brain maps 4.0—Structure of the rat brain: An open access atlas with global nervous system nomenclature ontology and flatmaps. Journal of Comparative Neurology 526, 935–943 (2018).

49. Kim, S.Y., et al. Stochastic electrotransport selectively enhances the transport of highly electromobile molecules. Proceedings of the National Academy of Sciences of the United States of America 112, E6274–6283 (2015).

50. Park, Y.G., et al. Protection of tissue physicochemical properties using polyfunctional crosslinkers. Nature biotechnology (2018).

51. Tsiola, A., Hamzei-Sichani, F., Peterlin, Z. & Yuste, R. Quantitative morphologic classification of layer 5 neurons from mouse primary visual cortex. Journal of Comparative Neurology 461, 415–428 (2003).

52. Vasques, X., Vanel, L., Villette, G. & Cif, L. Morphological neuron classification using machine learning. Frontiers in neuroanatomy 10, 102 (2016).

53. Aggleton, J. & Mishkin, M. Projections of the amygdala to the thalamus in the cynomolgus monkey. Journal of comparative neurology 222, 56–68 (1984).

54. Matyas, F., Lee, J., Shin, H.S. & Acsady, L. The fear circuit of the mouse forebrain: connections between the mediodorsal thalamus, frontal cortices and basolateral amygdala. The European journal of neuroscience 39, 1810–1823 (2014).

55. Cruikshank, S.J., Urabe, H., Nurmikko, A.V. & Connors, B.W. Pathway-specific feedforward circuits between thalamus and neocortex revealed by selective optical stimulation of axons. Neuron 65, 230–245 (2010).

56. Petreanu, L., Mao, T., Sternson, S.M. & Svoboda, K. The subcellular organization of neocortical excitatory connections. Nature 457, 1142–1145 (2009).

57. Swanson, L.W. & Petrovich, G.D. What is the amygdala? Trends in neurosciences 21, 323–331 (1998).

58. Ascoli, G.A. Sharing neuron data: carrots, sticks, and digital records. PLoS biology 13, e1002275 (2015).

59. Tohmi, M., Meguro, R., Tsukano, H., Hishida, R. & Shibuki, K. The extrageniculate visual pathway generates distinct response properties in the higher visual areas of mice. Current biology : CB 24, 587–597 (2014).

60. Jankowski, M.M., et al. Nucleus reuniens of the thalamus contains head direction cells. eLife 3 (2014).

61. Freese, J.L. & Amaral, D.G. The organization of projections from the amygdala to visual cortical areas TE and V1 in the macaque monkey. The Journal of comparative neurology 486, 295–317 (2005).

62. Iwai, E. & Yukie, M. Amygdalofugal and amygdalopetal connections with modality-specific visual cortical areas in macaques (Macaca fuscata, M. mulatta, and M. fascicularis). The Journal of comparative neurology 261, 362–387 (1987).

63. Gothard, K.M., Battaglia, F.P., Erickson, C.A., Spitler, K.M. & Amaral, D.G. Neural responses to facial expression and face identity in the monkey amygdala. Journal of neurophysiology 97, 1671–1683 (2007).

64. Hoffman, K.L., Gothard, K.M., Schmid, M.C. & Logothetis, N.K. Facial-expression and gaze-selective responses in the monkey amygdala. Current biology : CB 17, 766–772 (2007).

65. Hortensius, R., et al. The role of the basolateral amygdala in the perception of faces in natural contexts. Philosophical transactions of the Royal Society of London. Series B, Biological sciences 371 (2016).

66. Uwano, T., Nishijo, H., Ono, T. & Tamura, R. Neuronal responsiveness to various sensory stimuli, and associative learning in the rat amygdala. Neuroscience 68, 339–361 (1995).

67. Malkova, L., Gaffan, D. & Murray, E.A. Excitotoxic lesions of the amygdala fail to produce impairment in visual learning for auditory secondary reinforcement but interfere with reinforcer devaluation effects in rhesus monkeys. The Journal of neuroscience : the official journal of the Society for Neuroscience 17, 6011–6020 (1997).

68. Accolla, R., Bathellier, B., Petersen, C.C. & Carleton, A. Differential spatial representation of taste modalities in the rat gustatory cortex. The Journal of neuroscience : the official journal of the Society for Neuroscience 27, 1396–1404 (2007).

69. Chen, X., Gabitto, M., Peng, Y., Ryba, N.J. & Zuker, C.S. A gustotopic map of taste qualities in the mammalian brain. Science (New York, N.Y.) 333, 1262–1266 (2011).

70. Peng, Y., et al. Sweet and bitter taste in the brain of awake behaving animals. Nature 527, 512–515 (2015).

71. Wang, L., et al. The coding of valence and identity in the mammalian taste system. Nature 558, 127–131 (2018).

72. Sripanidkulchai, K., Sripanidkulchai, B. & Wyss, J.M. The cortical projection of the basolateral amygdaloid nucleus in the rat: a retrograde fluorescent dye study. The Journal of comparative neurology 229, 419–431 (1984).

73. Mandelbaum, G., et al. Distinct Cortical-Thalamic-Striatal Circuits through the Parafascicular Nucleus. Neuron 102, 636–652.e637 (2019).

74. Saper, C. & Loewy, A. Efferent connections of the parabrachial nucleus in the rat. Brain research 197, 291–317 (1980).

75. Fontanini, A., Grossman, S.E., Figueroa, J.A. & Katz, D.B. Distinct subtypes of basolateral amygdala taste neurons reflect palatability and reward. The Journal of neuroscience : the official journal of the Society for Neuroscience 29, 2486–2495 (2009).

76. Desgranges, B., Ramirez-Amaya, V., Ricano-Cornejo, I., Levy, F. & Ferreira, G. Flavor preference learning increases olfactory and gustatory convergence onto single neurons in the basolateral amygdala but not in the insular cortex in rats. PloS one 5, e10097 (2010).

77. Krettek, J.E. & Price, J.L. Amygdaloid projections to subcortical structures within the basal forebrain and brainstem in the rat and cat. The Journal of comparative neurology 178, 225–254 (1978).

78. Novejarque, A., Gutierrez-Castellanos, N., Lanuza, E. & Martinez-Garcia, F. Amygdaloid projections to the ventral striatum in mice: direct and indirect chemosensory inputs to the brain reward system. Frontiers in neuroanatomy 5, 54 (2011).

79. Wright, C.I., Beijer, A.V. & Groenewegen, H.J. Basal amygdaloid complex afferents to the rat nucleus accumbens are compartmentally organized. The Journal of neuroscience : the official journal of the Society for Neuroscience 16, 1877–1893 (1996).

80. Bossert, J.M., Gray, S.M., Lu, L. & Shaham, Y. Activation of group II metabotropic glutamate receptors in the nucleus accumbens shell attenuates context-induced relapse to heroin seeking. Neuropsychopharmacology : official publication of the American College of Neuropsychopharmacology 31, 2197–2209 (2006).

81. Fuchs, R.A., Ramirez, D.R. & Bell, G.H. Nucleus accumbens shell and core involvement in drug context-induced reinstatement of cocaine seeking in rats. Psychopharmacology 200, 545–556 (2008).

82. Gibson, G.D., Millan, E.Z. & McNally, G.P. The nucleus accumbens shell in reinstatement and extinction of drug seeking. The European journal of neuroscience (2018).

83. Bossert, J.M. & Stern, A.L. Role of ventral subiculum in context-induced reinstatement of heroin seeking in rats. Addiction biology 19, 338–342 (2014).

84. Lasseter, H.C., Xie, X., Ramirez, D.R. & Fuchs, R.A. Sub-region specific contribution of the ventral hippocampus to drug context-induced reinstatement of cocaine-seeking behavior in rats. Neuroscience 171, 830–839 (2010).

85. Luo, A.H., Tahsili-Fahadan, P., Wise, R.A., Lupica, C.R. & Aston-Jones, G. Linking context with reward: a functional circuit from hippocampal CA3 to ventral tegmental area. Science (New York, N.Y.) 333, 353–357 (2011).

86. Hamlin, A.S., Clemens, K.J., Choi, E.A. & McNally, G.P. Paraventricular thalamus mediates context-induced reinstatement (renewal) of extinguished reward seeking. The European journal of neuroscience 29, 802–812 (2009).

87. Bossert, J.M., et al. Ventral medial prefrontal cortex neuronal ensembles mediate context-induced relapse to heroin. Nature neuroscience 14, 420–422 (2011).

88. Millan, E.Z. & McNally, G.P. Accumbens shell AMPA receptors mediate expression of extinguished reward seeking through interactions with basolateral amygdala. Learning & memory (Cold Spring Harbor, N.Y.) 18, 414–421 (2011).

89. Fuchs, R.A., Eaddy, J.L., Su, Z.I. & Bell, G.H. Interactions of the basolateral amygdala with the dorsal hippocampus and dorsomedial prefrontal cortex regulate drug context-induced reinstatement of cocaine-seeking in rats. The European journal of neuroscience 26, 487–498 (2007).

90. Mashhoon, Y., Wells, A.M. & Kantak, K.M. Interaction of the rostral basolateral amygdala and prelimbic prefrontal cortex in regulating reinstatement of cocaine-seeking behavior. Pharmacology, biochemistry, and behavior 96, 347–353 (2010).

91. Marchant, N.J., Kaganovsky, K., Shaham, Y. & Bossert, J.M. Role of corticostriatal circuits in context-induced reinstatement of drug seeking. Brain research 1628, 219–232 (2015).

92. Ikemoto, S. Dopamine reward circuitry: two projection systems from the ventral midbrain to the nucleus accumbens–olfactory tubercle complex. Brain research reviews 56, 27–78 (2007).

93. Ikemoto, S., Qin, M. & Liu, Z.H. The functional divide for primary reinforcement of D-amphetamine lies between the medial and lateral ventral striatum: is the division of the accumbens core, shell, and olfactory tubercle valid? The Journal of neuroscience : the official journal of the Society for Neuroscience 25, 5061–5065 (2005).

94. Shin, R., Qin, M., Liu, Z.H. & Ikemoto, S. Intracranial self-administration of MDMA into the ventral striatum of the rat: differential roles of the nucleus accumbens shell, core, and olfactory tubercle. Psychopharmacology 198, 261–270 (2008).

95. Bossert, J.M., Poles, G.C., Wihbey, K.A., Koya, E. & Shaham, Y. Differential effects of blockade of dopamine D1-family receptors in nucleus accumbens core or shell on reinstatement of heroin seeking induced by contextual and discrete cues. Journal of Neuroscience 27, 12655–12663 (2007).

96. Xie, X., et al. Subregion-specific role of glutamate receptors in the nucleus accumbens on drug context-induced reinstatement of cocaine-seeking behavior in rats. Addiction biology 17, 287–299 (2012).

97. Boccara, C.N., et al. Grid cells in pre- and parasubiculum. Nature neuroscience 13, 987–994 (2010).

98. Felix-Ortiz, A.C. & Tye, K.M. Amygdala inputs to the ventral hippocampus bidirectionally modulate social behavior. The Journal of neuroscience : the official journal of the Society for Neuroscience 34, 586–595 (2014).

99. Swanson, L. Brain maps: structure of the rat brain (Gulf Professional Publishing, 2004).

100. Berendse, H.W. & Groenewegen, H.J. Organization of the thalamostriatal projections in the rat, with special emphasis on the ventral striatum. The Journal of comparative neurology 299, 187–228 (1990).

101. Berendse, H.W. & Groenewegen, H.J. Restricted cortical termination fields of the midline and intralaminar thalamic nuclei in the rat. Neuroscience 42, 73–102 (1991).

102. Rasmusson, A.M. & Charney, D.S. Animal models of relevance to PTSD. Annals of the New York Academy of Sciences 821, 332–351 (1997).

103. Wolpe, J. & Rowan, V.C. Panic disorder: a product of classical conditioning. Behaviour research and therapy 26, 441–450 (1988).

104. Yehuda, R. Current status of cortisol findings in post-traumatic stress disorder. The Psychiatric clinics of North America 25, 341–368, vii (2002).

105. Corcoran, K.A., Desmond, T.J., Frey, K.A. & Maren, S. Hippocampal inactivation disrupts the acquisition and contextual encoding of fear extinction. The Journal of neuroscience : the official journal of the Society for Neuroscience 25, 8978–8987 (2005).

106. Hobin, J.A., Ji, J. & Maren, S. Ventral hippocampal muscimol disrupts context-specific fear memory retrieval after extinction in rats. Hippocampus 16, 174–182 (2006).

107. Maren, S. & Fanselow, M.S. Synaptic plasticity in the basolateral amygdala induced by hippocampal formation stimulation in vivo. The Journal of neuroscience : the official journal of the Society for Neuroscience 15, 7548–7564 (1995).

108. Phillips, R.G. & LeDoux, J.E. Differential contribution of amygdala and hippocampus to cued and contextual fear conditioning. Behavioral neuroscience 106, 274–285 (1992).

109. Baeg, E.H., et al. Fast spiking and regular spiking neural correlates of fear conditioning in the medial prefrontal cortex of the rat. Cerebral cortex (New York, N.Y. : 1991) 11, 441–451 (2001).

110. Chang, C.H. & Maren, S. Strain difference in the effect of infralimbic cortex lesions on fear extinction in rats. Behavioral neuroscience 124, 391–397 (2010).

111. Corcoran, K.A. & Quirk, G.J. Activity in prelimbic cortex is necessary for the expression of learned, but not innate, fears. The Journal of neuroscience : the official journal of the Society for Neuroscience 27, 840–844 (2007).

112. Knapska, E., et al. Functional anatomy of neural circuits regulating fear and extinction. Proceedings of the National Academy of Sciences of the United States of America 109, 17093–17098 (2012).

113. Knapska, E. & Maren, S. Reciprocal patterns of c-Fos expression in the medial prefrontal cortex and amygdala after extinction and renewal of conditioned fear. Learning & memory (Cold Spring Harbor, N.Y.) 16, 486–493 (2009).

114. Milad, M.R. & Quirk, G.J. Neurons in medial prefrontal cortex signal memory for fear extinction. Nature 420, 70–74 (2002).

115. Quirk, G.J., Russo, G.K., Barron, J.L. & Lebron, K. The role of ventromedial prefrontal cortex in the recovery of extinguished fear. The Journal of neuroscience : the official journal of the Society for Neuroscience 20, 6225–6231 (2000).

116. Zelikowsky, M., et al. Prefrontal microcircuit underlies contextual learning after hippocampal loss. Proceedings of the National Academy of Sciences of the United States of America 110, 9938–9943 (2013).

117. McDonald, A.J., Shammah-Lagnado, S.J., Shi, C. & Davis, M. Cortical afferents to the extended amygdala. Annals of the New York Academy of Sciences 877, 309–338 (1999).

118. Frankland, P.W., Bontempi, B., Talton, L.E., Kaczmarek, L. & Silva, A.J. The involvement of the anterior cingulate cortex in remote contextual fear memory. Science (New York, N.Y.) 304, 881–883 (2004).

119. Holland, P.C. & Gallagher, M. Amygdala circuitry in attentional and representational processes. Trends in cognitive sciences 3, 65–73 (1999).

120. Morrison, S.E. & Salzman, C.D. Re-valuing the amygdala. Current opinion in neurobiology 20, 221–230 (2010).

121. Hatfield, T., Han, J.S., Conley, M., Gallagher, M. & Holland, P. Neurotoxic lesions of basolateral, but not central, amygdala interfere with Pavlovian second-order conditioning and reinforcer devaluation effects. The Journal of neuroscience : the official journal of the Society for Neuroscience 16, 5256–5265 (1996).

122. Johnson, A.W., Gallagher, M. & Holland, P.C. The basolateral amygdala is critical to the expression of pavlovian and instrumental outcome-specific reinforcer devaluation effects. The Journal of neuroscience : the official journal of the Society for Neuroscience 29, 696–704 (2009).

123. Ostlund, S.B. & Balleine, B.W. Differential involvement of the basolateral amygdala and mediodorsal thalamus in instrumental action selection. The Journal of neuroscience : the official journal of the Society for Neuroscience 28, 4398–4405 (2008).

124. Izquierdo, A. & Murray, E.A. Selective bilateral amygdala lesions in rhesus monkeys fail to disrupt object reversal learning. The Journal of neuroscience : the official journal of the Society for Neuroscience 27, 1054–1062 (2007).

125. Murray, E.A. & Izquierdo, A. Orbitofrontal cortex and amygdala contributions to affect and action in primates. Annals of the New York Academy of Sciences 1121, 273–296 (2007).

126. Wassum, K.M., Ostlund, S.B., Maidment, N.T. & Balleine, B.W. Distinct opioid circuits determine the palatability and the desirability of rewarding events. Proceedings of the National Academy of Sciences of the United States of America 106, 12512–12517 (2009).

127. Lichtenberg, N.T., et al. Basolateral Amygdala to Orbitofrontal Cortex Projections Enable Cue-Triggered Reward Expectations. The Journal of neuroscience : the official journal of the Society for Neuroscience 37, 8374–8384 (2017).

128. Saddoris, M.P., Gallagher, M. & Schoenbaum, G. Rapid associative encoding in basolateral amygdala depends on connections with orbitofrontal cortex. Neuron 46, 321–331 (2005).

129. Schoenbaum, G., Chiba, A.A. & Gallagher, M. Orbitofrontal cortex and basolateral amygdala encode expected outcomes during learning. Nature neuroscience 1, 155–159 (1998).

130. Bechara, A., Damasio, H., Tranel, D. & Anderson, S.W. Dissociation Of working memory from decision making within the human prefrontal cortex. The Journal of neuroscience : the official journal of the Society for Neuroscience 18, 428–437 (1998).

131. Bechara, A., Damasio, H., Tranel, D. & Damasio, A.R. Deciding advantageously before knowing the advantageous strategy. Science (New York, N.Y.) 275, 1293–1295 (1997).

132. Rolls, E.T. The orbitofrontal cortex. Philosophical transactions of the Royal Society of London. Series B, Biological sciences 351, 1433–1443; discussion 1443-1434 (1996).

133. Kita, H. & Kitai, S.T. Amygdaloid projections to the frontal cortex and the striatum in the rat. The Journal of comparative neurology 298, 40–49 (1990).

134. Krettek, J.E. & Price, J.L. Projections from the amygdaloid complex to the cerebral cortex and thalamus in the rat and cat. The Journal of comparative neurology 172, 687–722 (1977).

135. Adhikari, A., et al. Basomedial amygdala mediates top-down control of anxiety and fear. Nature 527, 179 (2015).

136. Canteras, N.S. & Swanson, L.W. Projections of the ventral subiculum to the amygdala, septum, and hypothalamus: a PHAL anterograde tract-tracing study in the rat. The Journal of comparative neurology 324, 180–194 (1992).

137. Petrovich, G.D., Canteras, N.S. & Swanson, L.W. Combinatorial amygdalar inputs to hippocampal domains and hypothalamic behavior systems. Brain research. Brain research reviews 38, 247–289 (2001).

138. Petrovich, G.D., Risold, P.Y. & Swanson, L.W. Organization of projections from the basomedial nucleus of the amygdala: a PHAL study in the rat. The Journal of comparative neurology 374, 387–420 (1996).

139. Blondel, V.D., Guillaume, J.L., Lambiotte, R. & Lefebvre, E. Fast unfolding of communities in large networks. Journal of Statistical Mechanics: Theory and Experiment 2008, P10008 (2008).

140. Lancichinetti, A. & Fortunato, S. Consensus clustering in complex networks. Scientific reports 2, 336 (2012).

141. Cabeen, R., Laidlaw, D. & Toga, A. Quantitative imaging toolkit: Software for interactive 3D visualization, data exploration, and computational analysis of neuroimaging datasets. in Proceedings of the International Society for Magnetic Resonance in Medicine (ISMRM) (2018).

142. Cleveland, W.S. & Devlin, S.J. Locally weighted regression: an approach to regression analysis by local fitting. Journal of the American statistical association 83, 596–610 (1988).

143. Kroner, A., et al. TNF and increased intracellular iron alter macrophage polarization to a detrimental M1 phenotype in the injured spinal cord. Neuron 83, 1098–1116 (2014).

144. Scorcioni, R., Polavaram, S. & Ascoli, G.A. L-Measure: a web-accessible tool for the analysis, comparison and search of digital reconstructions of neuronal morphologies. Nature protocols 3, 866 (2008).

145. Li, Y., Wang, D., Ascoli, G.A., Mitra, P. & Wang, Y. Metrics for comparing neuronal tree shapes based on persistent homology. PloS one 12, e0182184 (2017).

146. Kerber, M., Morozov, D. & Nigmetov, A. Geometry helps to compare persistence diagrams. Journal of Experimental Algorithmics (JEA) 22, 1.4 (2017).

