## Supplemental figures for "Connectivity characterization of the mouse basolateral amygdalar complex"

#### Supplementary Figure 1

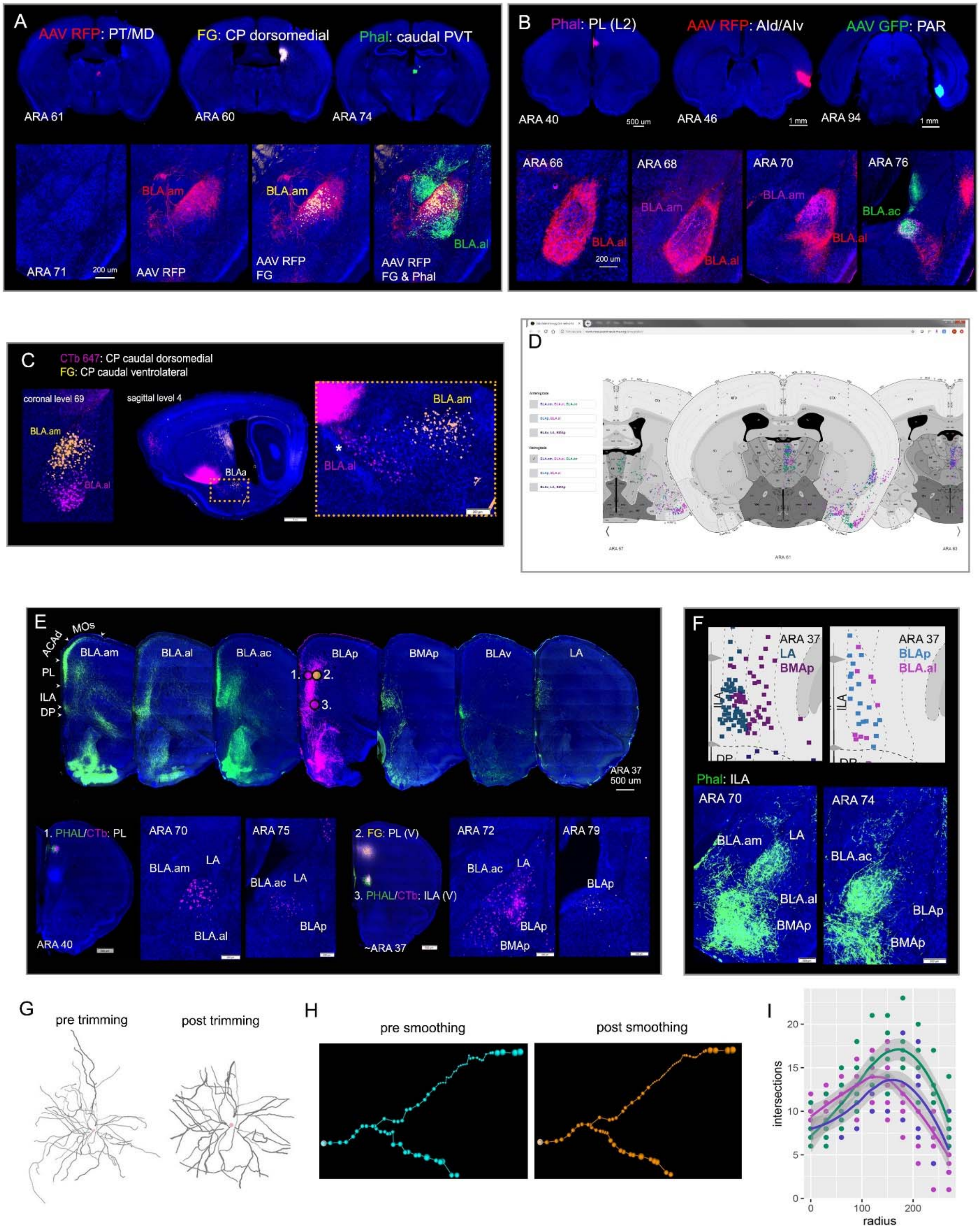

**Supplementary Figure 1**

**A.** Tracer injections made across thalamic and striatal nuclei show the distinct connections of BLA.am and BLA.al domains. Anterograde tracers AAV RFP and Phal placed thalamic PT/MD and caudal PVT, label either BLA.am or BLA.al, respectively. In the same animal, a FG injection in CP caudal dorsomedial labels BLA.am projection neurons, which overlap with AAV RFP labeled fibers from PT. *Abbreviations: PT: parataenial thalamic nucleus; MD: mediodorsal thalamic nucleus; PVT: paraventricular thalamic nucleus.* **B.** Additional example of multiple anterograde tracer injections placed in different cortical regions within the same animal labeling distinct BLAa domains across ARA levels 66-76. Anterograde Phal in PL (pink) labels fibers in BLA.am, while AAV RFP in AId/AIv (red) labels fibers in BLA.al, and AAV GFP in PAR (green) labels fibers in BLA.ac. *Abbreviations: PL: prelimbic cortical area; AId: agranular insular area, dorsal part; AIv: agranular insular area, ventral part; PAR: parasubiculum.* **C.** Retrograde tracers CTb 647 (pink) and FG (yellow) injected in either the dorsomedial or ventrolateral part of the caudal caudoputamen distinctly label BLA.am and BLA.al projection neurons, respectively, which is evident in both coronal (left) and sagittal (right) views. Inset shows magnified version of boxed region on the sagittal section. **D.** Snapshot of the application created to view all of the quantified and analyzed data, which will be freely available online (http://www.mouseconnectome.org/amygdalar/) (currently password protected). From the left tabs, users can select the anterograde or retrograde dataset to view: (1) BLA.am, BLA.al, BLA.ac; (2) BLAp, BLA.al; or (3) BLAv, LA, BMAp. Using the left-right arrows at the bottom, users can navigate through the 32 different ARA levels. **E.** Top row shows projections to medial prefrontal cortex from projection neurons in BLA.am, BLA.al, BLA.ac, BLAp, BMAp, BLAv, and LA. Connections from BLAp to deeper layers of ILA and PL were validated with retrograde tracer injections placed in 1. superficial layers of PL(II/III) (CTb: pink), 2. deeper layers of PL(V/VI) (FG: yellow), and 3. deeper layers of ILA(V/VI) (CTb: pink). Bottom row: CTb injection in PL(II/III) weakly labels BLAp neurons. This injection also validates BLA.am/BLA.ac→PL(II/III) connection. FG in PL(V/VI) and CTb in ILA(V/VI) back-label mostly BLAp neurons confirming BLAp→PL/ILA (deeper layers). *Abbreviations: PL: prelimbic cortical area; ILA: infralimbic cortical area.* **F.** Validation of strong ILA→LA and ILA(V)→BMAp connections and weak ILA→BLAp and ILA→BLA.al connections. Top row: retrograde visuals showing back-labeled neurons in ILA following retrograde tracers placed in LA, BMAp, and BLAp. Bottom row: validation of those connections with an ILA Phal injection (injection 3 in panel E). Phal label can be seen in LA, BMAp, BLA.am, and BLA.ac but far less in the BLA.al and BLAp. **G.** Due to differences in slice thickness for the reconstruction of BLA.am (250 μm), BLA.al (250 μm), and BLA.ac (400 μm) projection neurons, morphological analysis of neurites was restricted by trimming the portions of the dendritic branches that measured farther than 300 nm from the soma. The left shows a reconstructed striatum projecting BLA.ac neuron in its entirety, while the right shows the same neuron with applied trimming. **H.** Due to anisotropic voxels and spatial undersampling relative to the real curvature of the dendrite, a locally weighted scatter-plot smoother (LOESS) was applied to the data. The left shows curvature prior to smoothing, while the left shows post smoothing curvature. **I.** Result of Sholl analysis to show overall view of BLAa projection neuron dendritic morphology. Scatter plot shows that dendrites of neurons within BLA.am and BLA.al mostly differ from those in BLA.ac within 100-200 nm radius from the cell body, with BLA.ac dendrites showing a greater number of intersections within that range.

#### Supplementary Figure 2

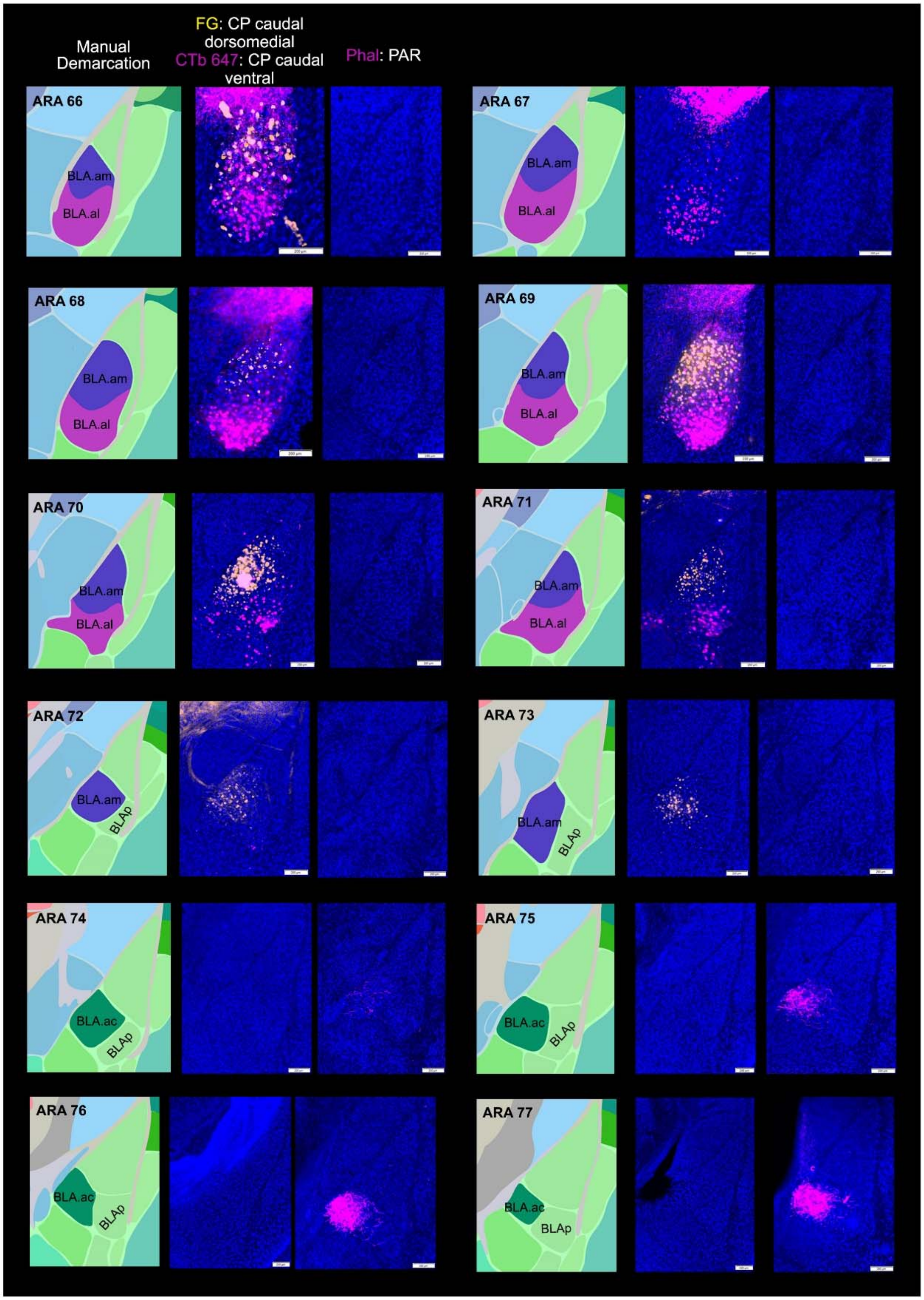

**Supplementary Figure 2**

For demarcation of BLA.am, BLA.al, and BLA.ac borders, cases with distinct labels within each domain were selected and contiguous sections through the BLAa from ARA level 66 to ARA77 were collected (n=7). In the interest of space, two representative cases are shown: one in which FG and CTb were injected in CP caudal dorsomedial and CP caudal ventral, respectively and another in which Phal was injected in PAR. Note the clear FG-labeled BLA.am projection neurons from ARA 67-73. At level 74 also note the absence of FG labeling in BLA.am and the start of Phal label from the PAR injection. CTb labeled BLA.al projection neurons are observed from ARA 66-71, after which they disappear at ARA 72 and BLAp appears. Initially, the borders were drawn manually guided by mapped labels, but also by Nissl stains for each case (Supplementary Figure 3B). The same data was used as a training set for a machine learning algorithm, which produced similar border delineations of BLAa domains (Supplementary Figure 3A).

#### Supplementary Figure 3

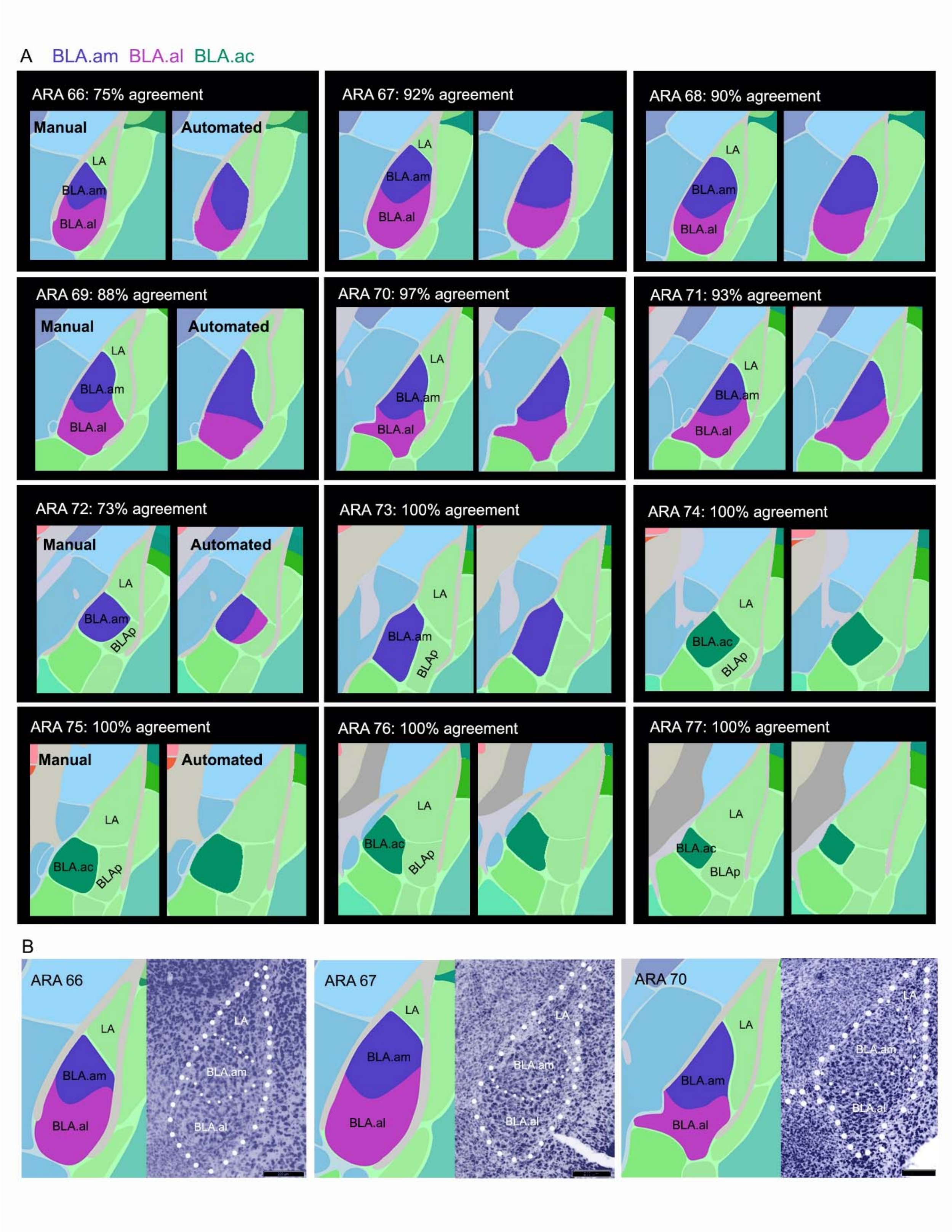

**Supplementary Figure 3**

**A.** Output of manual versus automated boundary demarcation for BLA.am, BLA.al, and BLA.ac across ARA levels 66-77. Initially, borders were drawn manually guided by mapped labels, but also by Nissl stains for each case. The same data was used as a training set for a machine learning algorithm, which produced similar border delineations of BLAa domains. On average, across all sections there was a 92% agreement between the manually drawn versus the automatically detected boundaries. The lowest agreements were at ARA 66 and 72 with 75% and 73% agreements, respectively. These percentages were reasonable for both levels given that at more rostral levels of BLAa like ARA 66, segregation of labels was far less clear, hence the exclusion of ARA levels 63-65 despite the presence of BLAa at those levels. ARA 72 is the first section where the BLA.al disappears so it is not surprising for the automated method to identify it as a transitioning section in which both BLA.am and some BLA.al are present. **B.** Importantly, the boundary between the BLA.am and BLA.al was also apparent from Nissl stains. A representative Nissl case showing the BLA.am BLA.al distinction is presented here with delineated boundaries for ARA levels 66-70.

**Supplementary Figure 4**

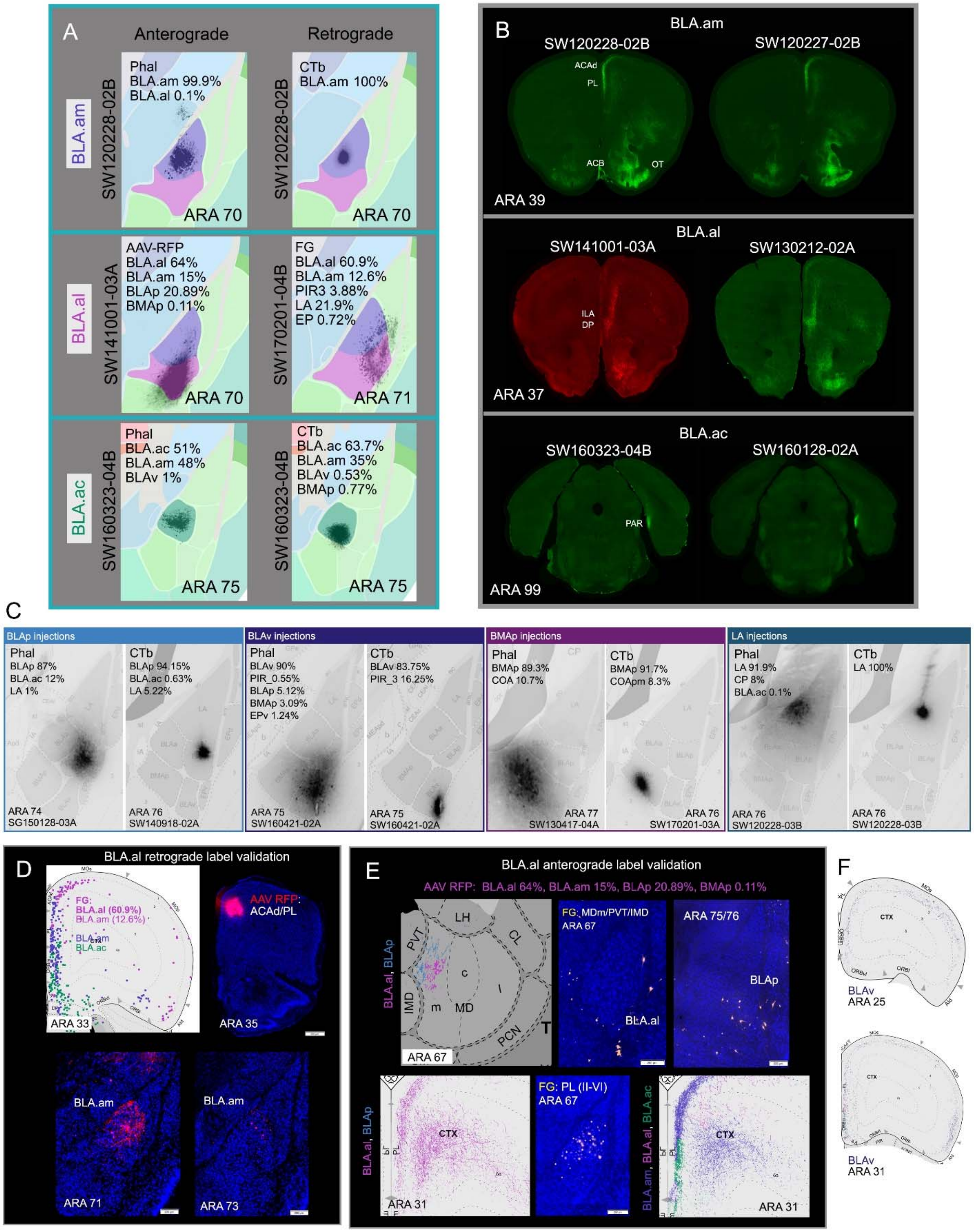

**Supplementary Figure 4**

**A.** Representative images of tracer injections in BLA.am, BLA.al, and BLA.ac. Percentages represent amount of tracer calculated in each ROI (see Supplementary methods). Although a single image is presented, percentages were calculated based on all sections that contained the tracer injection. **B.** Anterograde labeling of main injection cases and validation cases for BLA.am, BLA.al, and BLA.ac showing consistent patterns of labeling. **C.** Representative images and percentages of anterograde and retrograde tracer injections in BLAp, BLAv, BMAp, and LA. **D.** Examples of label cross validation using the BLA.al as an example. A FG injection in mostly the BLA.al that encroached into the BLA.am shows back-labeled neurons in ACAd (pink cells on retrograde visual) suggesting an ACAd→BLA.al connection. An anterograde injection in ACAd shows label only in BLA.am, suggesting the back-label in ACAd from the BLA.al FG injection was most likely from leakage into BLA.am (purple cells) (ACAd→BLA.am). *Abbreviations: ACAd: anterior cingulate cortex, dorsal part.* **E.** Two additional examples illustrating the importance of cross validation of connections. Top panels: an AAV RFP injection in mostly BLA.al that encroached into BLAp shows labeled fibers in MDm (pink fibers on anterograde visual) indicating a BLA.al→MDm connection. A FG injection in MDm back-labels only BLAp projection neurons suggesting the FG label from the BLA.al injection originates from BLAp (blue fibers) and not BLA.al (BLAp→MDm). Bottom panels: the same AAV-RFP BLA.al injection also leaked into BLA.am and labeled different layers of PL (pink fibers in anterograde visual to the left). A FG injection in PL back-labels mostly BLA.am neurons suggesting the labeled PL fibers from the BLA.al injection was from leakage into BLA.am. Note that the anterograde visual on the left is from grouping of BLA.al and BLAp and the one to the right is from grouping the BLA.al with BLA.am and BLA.ac. In the latter, the PL and its layers get assigned to the BLA.am injection site rather than to BLA.al due to the stronger BLA.am→PL connections at ARA level 31. *Abbreviations: MDm: mediodorsal thalamic nucleus, medial part; PL: prelimbic cortical area.* **F.** Anterograde visuals showing BLAv→MOs(I) projections at ARA 25 and 31. *Abbreviations: MOs: secondary motor cortex.*

#### Supplementary Figure 5

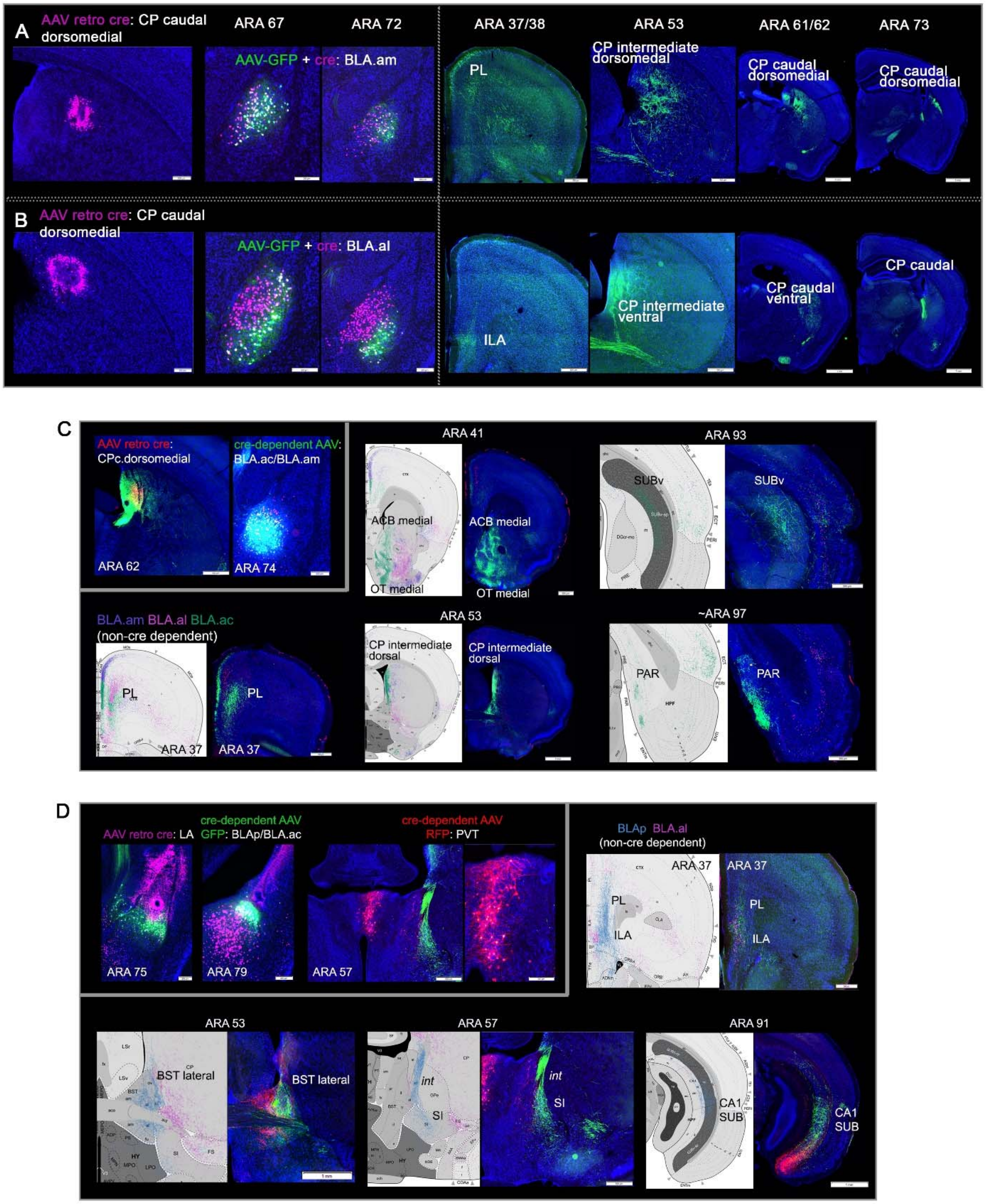

**Supplementary Figure 5**

**A.** Cre-dependent anterograde AAV validation of BLA.am projection neurons. Left panels: AAV retro Cre was injected into dorsomedial part of CP caudal to retrogradely deliver Cre to BLA.am neurons (specifically to CP projecting BLA.am neurons. A Cre-dependent anterograde AAV tracer was injected into BLA.am. Note pink (retro Cre-AAV) and green (AAV-GFP) co-labeled cells in BLA.am. Right panels: anterograde fiber labeling in PL(II/VI), CP intermediate, and CP caudal look identical to the non Cre-dependent BLA.am anterograde case. *Abbreviations: PL: prelimbic cortical area; CP: caudoputamen.* **B.** Left panels: AAV retro Cre was injected into dorsomedial part of CP caudal to retrogradely deliver Cre to BLA.am neurons, which also labeled some BLA.al projection neurons. A Cre-dependent anterograde AAV tracer was injected into BLA.al. Note pink (retro Cre-AAV) and green (AAV-GFP) co-labeled cells in BLA.al. Right panels: anterograde fiber labeling in ILA, CP intermediate, and CP caudal are identical to the non-Cre-dependent BLA.al anterograde case, showing that BLA.al neurons project to ILA, to mostly CP ventral, although some projections to dorsal regions are also evident, but unlike the BLA.am case, are not as dense to CP caudal. *Abbreviations: ILA: infralimbic cortical area; CP: caudoputamen.* **C.** Inset: AAV retro Cre was injected into dorsomedial part of CP caudal to retrogradely deliver Cre to BLA.am, which also labeled BLA.ac projection neurons. A Cre-dependent anterograde AAV tracer was injected primarily into the BLA.ac. Note red (retro Cre-AAV) and green (AAV-GFP) co-labeled cells in BLA.ac. Anterograde visuals are presented for the non-Cre dependent tracing to demonstrate similarity in labels in the ventral parts of the PL (ARA 37), medial ACB (ARA 41), medial OT (ARA 41), CP intermediate (ARA 53), SUBv (ARA 93), and PAR (ARA 97). *Abbreviations: PL: prelimbic cortical area; ACB: nucleus accumbens; OT: olfactory tubercle; CP: caudoputamen; SUBv: subiculum, ventral part; PAR: parasubiculum.* **D.** Inset: AAV retro Cre was injected into LA, which labeled projection neurons in BLAp and some in BLA.ac. A Cre-dependent anterograde AAV GFP tracer was injected primarily into BLAp and a Cre-dependent anterograde AAV-RFP into the PVT. Note pink (retro Cre-AAV) and green (AAV-GFP) co-labeled cells in BLAp and pink (retro-Cre-AAV) and red (AAV-RFP) in the PVT. PVT is shows to underscore the distinct topographic projections from BLAp projection neurons. Anterograde visuals are presented for the non-Cre dependent tracing to demonstrate consistency of labels in deep layers of PL and ILA (ARA 37), BST lateral (ARA 53), SI (ARA 57), and CA1 and SUBv (ARA 91). *Abbreviations: PL: prelimbic cortical area; ILA: infralimbic cortical area; BST: bed nucleus of stria terminalis; SI: substantia innominata; SUBv: subiculum, ventral part.*

#### Supplementary Figure 6

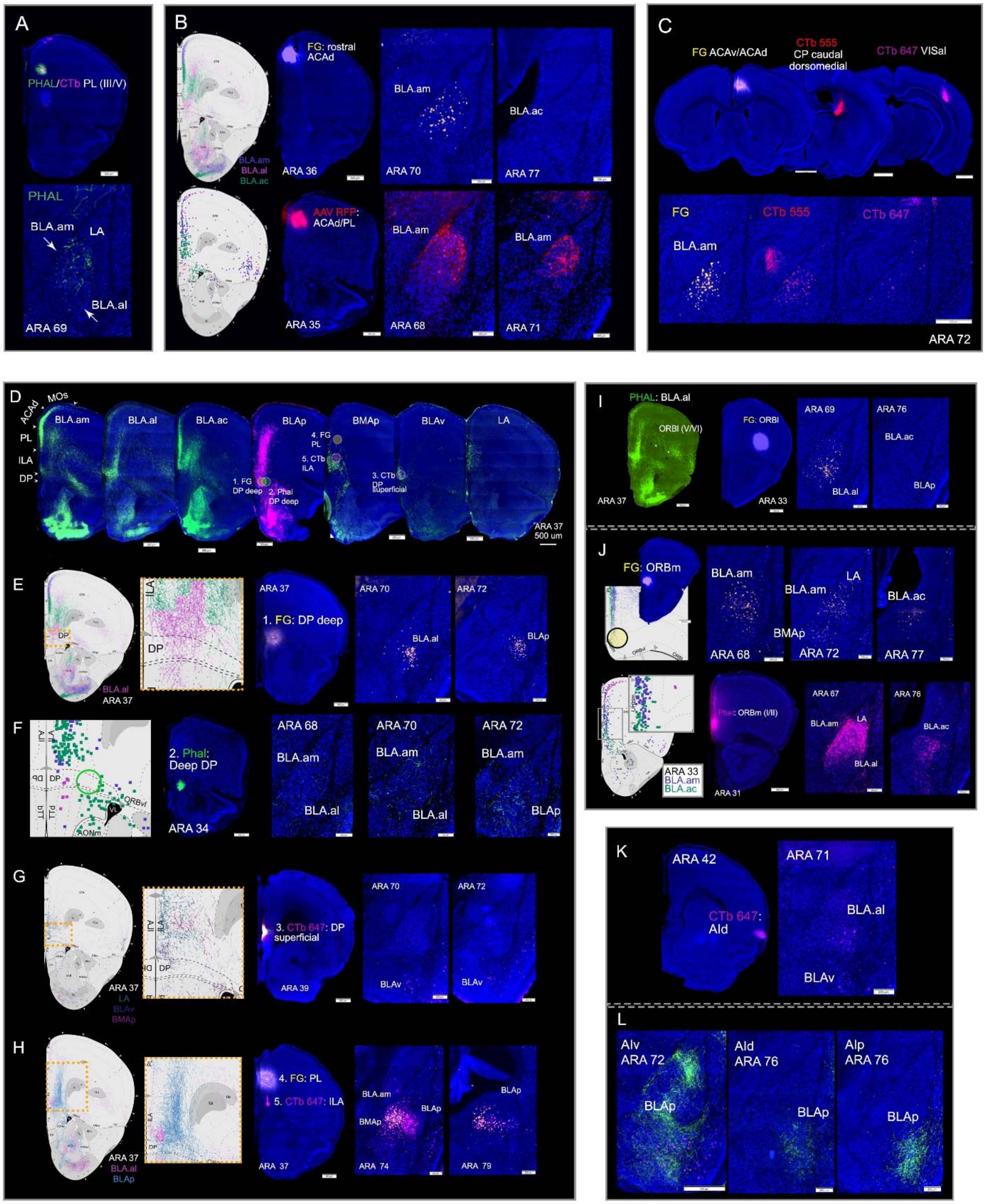

**Supplementary Figure 6**

**A.** Phal/CTb injected in deeper layers of PL (prelimbic cortical area) showing sparse label in BLA.am, confirming the sparser connections of PL layer III with BLA.am. **B.** Top panels: anterograde visual to the left shows strong projections from BLA.am neurons to rostral ACAd (anterior cingulate cortex, dorsal part). A FG injection in rostral ACAd labels projection cells in BLA.am only confirming the BLA.am→rostral ACAd connection. Bottom panels: retrograde visual to the left shows back-labeled neurons in rostral ACAd from a BLA.am injection. An AAV RFP injection in rostral ACAd confirms this rostral ACAd→BLA.am connection. **C.** Three retrograde tracers injected in ACA (FG), caudal CP dorsomedial (CTb 555), and in deep layers of VISal (CTb 647) back-label BLA.am projection neurons at ARA 72 validating BLA.am projection to visual processing areas (BLA.am→ACA/CP/VISal). *Abbreviations: ACAd: anterior cingulate cortex, dorsal part; ACAv: anterior cingulate cortex, ventral part: CP: caudoputamen; VISal: visual cortical area anterolateral part.* **D.** Projections to medial prefrontal cortical areas (MPF) from injections in BLA.am, BLA.al, BLA.ac, BLAp, BMAp, BLAv, and LA. Numbers denote injections made in different MPF regions in E-H to validate these connections. **E.** Anterograde visuals show BLA.al neuron projections to deep layers of DP (dorsal peduncular area). FG injection 1 in deep DP back-labels projections neurons in BLA.al and BLAp validating BLA.al/BLAp→DP deep layer connections. **F.** In general, projections from DP back to BLA are weak as shown by the retrograde visual to the left. Injection 2, a Phal injection in deep layers of DP, whose location is denoted by the ellipse on the visual, confirms these weak DP deep→BLA connections. **G.** Anterograde visuals to the left show BLAv projections to superficial layers of DP. Inset is a magnification of boxed region on the visual. Injection 3, a CTb injection in superficial layers of DP back-labels BLAv projection neurons validating the connection (BLAv→DP superficial). **H.** Anterograde visual showing BLAp projections to deep layers of PL and ILA. Inset is magnification of boxed region on the visual. Injections 4 and 5 are FG and CTb injections in PL (all layers) and ILA (deep layers). FG back-labels projection cells in BLAp and BLA.am confirming those connections (BLA.am/BLAp→PL), while CTb labels projection neurons mostly in BLAp and some in BMAp, validating BLAp/BMAp→PL (deeper layer) connections. *Abbreviations: PL: prelimbic cortical area; ILA: infralimbic cortical area.* **I.** Left panel shows BLA.al projections to ORBl (orbital area, lateral part), a connection validated by a FG injection in ORBl that back-labels BLA.al projection neurons (BLA.al→ORBl). **J.** Top panels: anterograde visual shows BLA.am and BLA.ac projections to ORBm (orbital area, medial part). A FG injection in ORB, denoted by the ellipse on the visual, back-labels projection neurons in BLA.am, BLA.ac, and also BMAp validating BLA.am/BLA.ac/BMAp→ORBm connections. Bottom panels: retrograde visual shows ORBm labeled neurons from injections in BLA.am and BLA.ac. Inset is magnification of boxed region on the visual. A Phal injection in the ORBm labels the BLA.am, BLA.ac, and LA validating BLA.am/BLA.ac/LA→ORBm projections. **K.** Validation of BLA.al/BLAv→AId connection with CTb injection in AId (agranular insular cortical area, dorsal part). **L.** AI projections to BLAp. Phal injections in AIv, AId, and AIp show relatively weak projections to BLAp. *Abbreviations: AId: agranular insular cortical area, dorsal part: AIv: agranular insular cortical area, ventral part; AIp: agranular insular cortical area, posterior part.*

#### Supplementary Figure 7

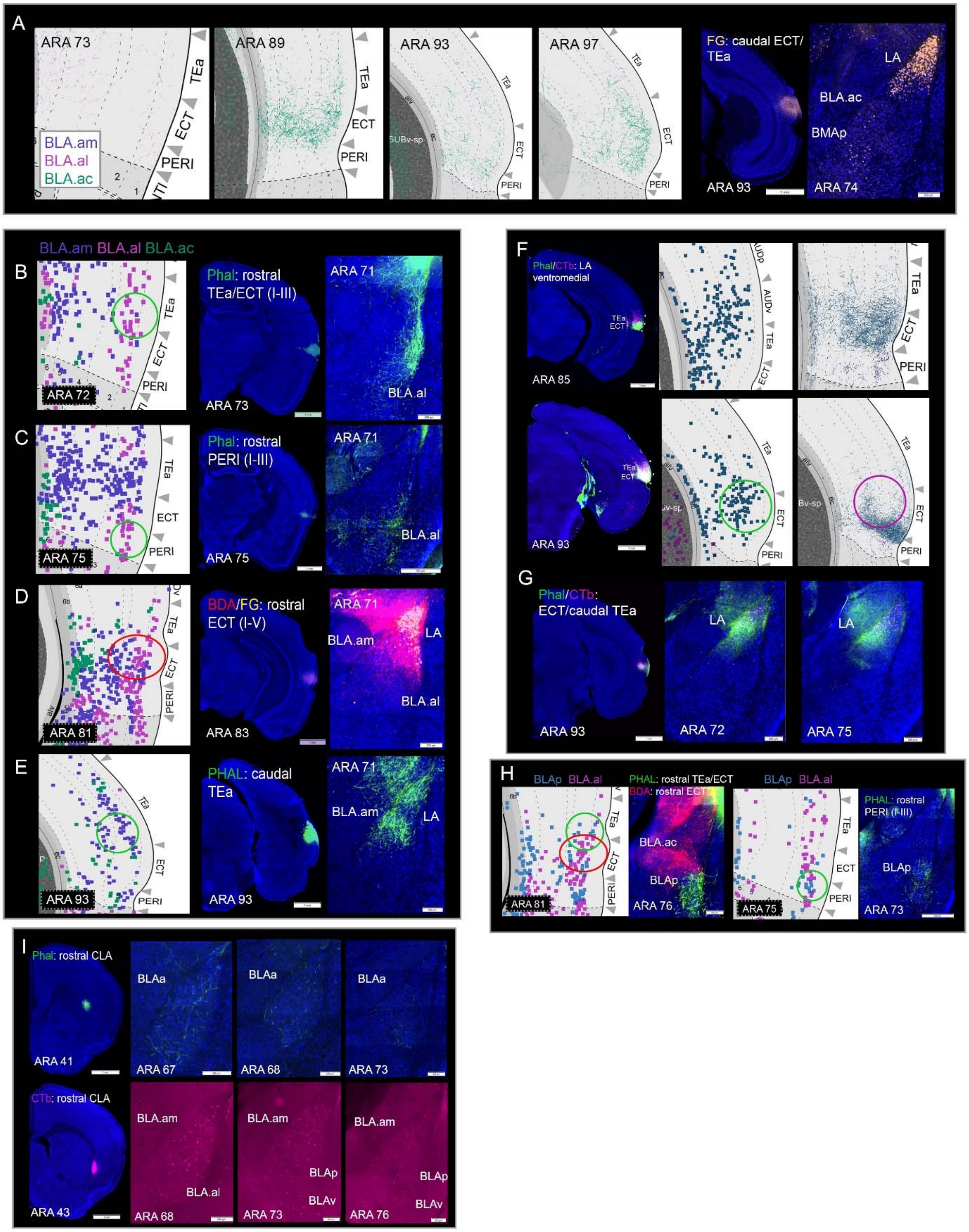

**Supplementary Figure 7**

**A.** Anterograde visuals showing weak projections from BLA.am, BLA.al, and BLA.ac to TEa, ECT, and PERI. Some BLA.ac projections are evident in caudal ECT. A FG injection in caudal ECT validates this weak BLA.ac→ECT caudal projection, but also shows weak projections from BMAp to ECT caudal (BMAp→ECT caudal), and shows that strongest projections to ECT caudal are from LA projection neurons. *Abbreviations: TEa: temporal association area; ECT: ectorhinal cortical area; PERI: perirhinal cortical area.* **B-E.** Topographic projections from perirhinal areas to BLAa. Left: retrograde visuals show rostral TEa/ECT (B), rostral PERI (C), and rostral ECT (D) labeled cells following retrograde tracer injections in BLA.al (layers II-III), but also in BLA.am (layers III-VI). E. shows caudal TEa cells labeled following retrograde injection in BLA.am. Ellipses denote location of anterograde Phal (green) or AAV-RFP (red) injections to validate rostral TEa/ECT/PERI→BLA.al (II/III) and caudal TEa→BLA.am connections to the right. D also shows strong reciprocal connections between rostral ECT and LA, while E demonstrates projections from caudal TEa→LA. **F.** Top panels: raw data and visuals showing LA projection neurons heavily innervating ECT and TEa. Panels also show ECT and TEa cells projecting strongly to LA at ARA levels 85 and 93. Strong Phal and CTb labeling is present in the TEa and ECT following a coinjection into LA (ventromedial part). Ellipses denote locations of anterograde Phal (green) and CTb (pink) injections to validate the LA→TEa/ECT and TEa/ECT→LA connections. **H.** Retrograde visuals show labeled cells in rostral regions of TEa, ECT, and PERI following a retrograde tracer injection into BLAp. Ellipses denote locations of anterograde tracer injections Phal (green) and BDA (red) to validate rostral TEa/ECT/PERI→BLAp projections. Note fiber labels in BLAp following anterograde tracer injections **I.** Top panels: Phal anterograde tracer in rostral claustrum (CLA) shows weak CLA→BLAa connections. Bottom panels: CTb retrograde tracer in rostral CLA shows weak BLAa→CLA connections.

### Supplementary Figure 8

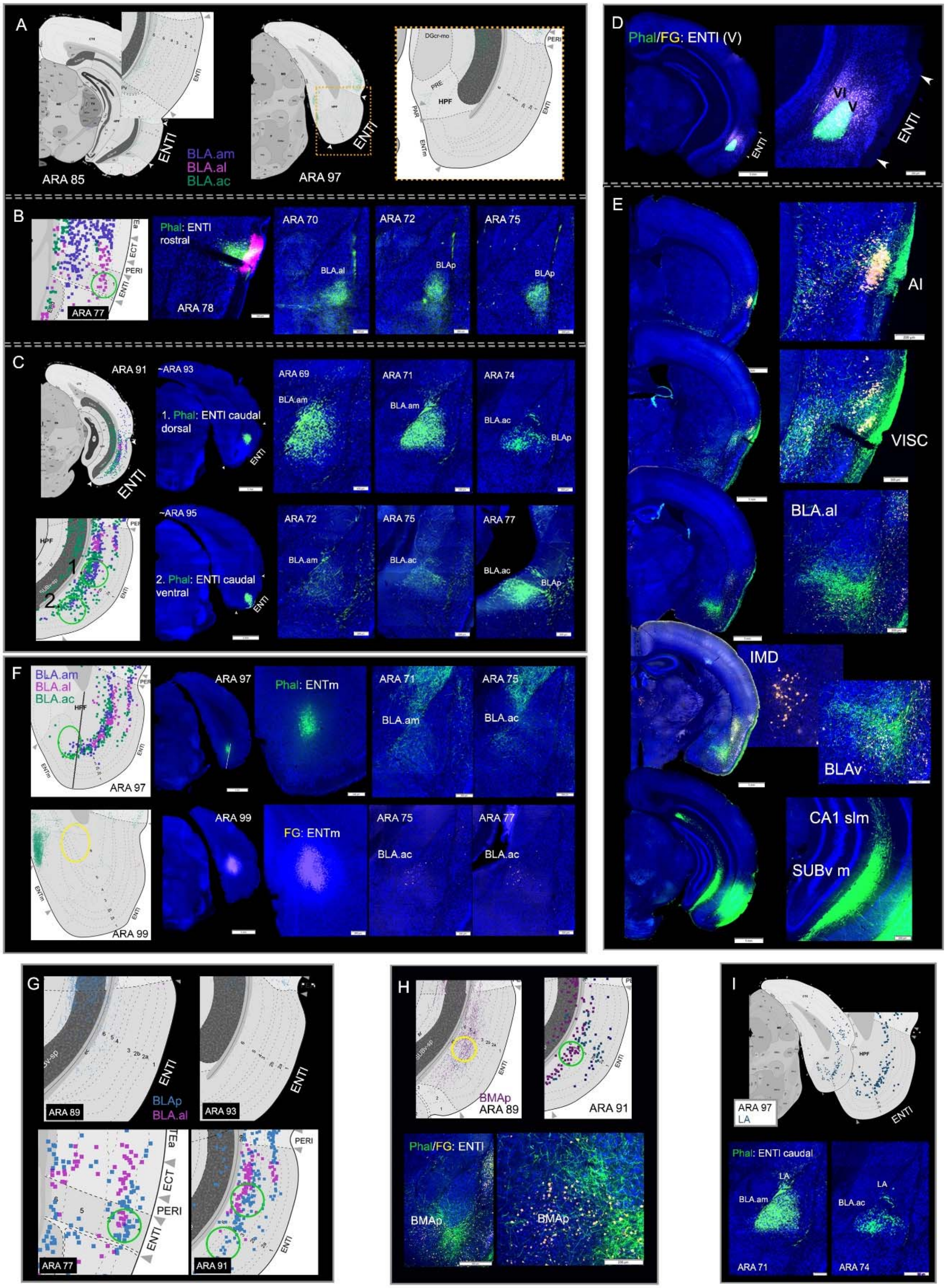


Add ARA 85 retrograde visual for BLAv
to show ENTl(V)-->BLAv

**Supplementary Figure 8**

**A.** Anterograde visuals from anterograde tracer injections placed in BLA.am, BLA.al, and BLA.ac showing weak projections from BLAa to ENTl (entorhinal cortical area, lateral part) at ARA levels 85 and 97. Inset is magnification of boxed region on the visual. **B.** ENTl shows topographic projections to different BLAa domains. Retrograde visual to the left shows back-labeled rostral ENTl projection neurons from a BLA.al retrograde injection. Ellipse denotes location of Phal (green) anterograde tracer injection that validates the rostral ENTl→BLA.al connection, and also shows rostral ENTl→BLAp projection. **C.** Left panels show retrograde visual with labeled caudal dorsal and ventral ENTl cells from retrograde injections in BLA.am and BLA.ac. Ellipses denote Phal injection locations in the dorsal (injection 1) and ventral (injection 2) caudal ENTl. Phal injections validate caudal dorsal ENTl→BLA.am/BLA.ac and caudal ventral ENTl→BLA.ac connections. In injection 1, note the sparser label in BLA.ac compared to BLA.am. In injection 2, note the strong label in BLA.ac. **D.** An anterograde tracer in the BLAv showed selective input to ENTl(V), a region that also receives input from AI (agranular insular cortical area) (Figure 6 U-V). The connectivity of ENTl(V) was examined with a Phal and FG injection. **E.** First, note FG labels in AI and BLAv projection neurons confirming the BLAv/AI→ENTl(V) connection. In addition, FG injection shows that this region receives input from VISC and IMD, two region also involved in gustatory/visceral processing. The Phal injection shows the output of ENTl(V) to AI, VISC, BLA.al, and SUBv. *Abbreviations: AI: agranular insular cortical area: VISC: visceral cortical area; IMD: intermediodorsal thalamic nucleus; SUBv: subiculum, ventral part.* **F.** Top panels show weak projections from ENTl neurons to BLA.am and BLA.ac. Bottom panels show weak, but observable projections from BLA.ac to ENTm (BLA.ac→ENTm). **G.** Top panels: weak or absent projections from BLAp neurons to ENTl. Bottom panels: retrograde visuals with back-labeled rostral (ARA 77) and caudal dorsal and ventral ENTl (ARA 91) neurons from a retrograde injection in BLAp. Ellipses denote Phal (green) injection locations that correspond to injections in B-C and validate rostral ENTl/caudal dorsal ENTl/caudal ventral ENTl→BLAp connections. **H.** Top panels: anterograde and retrograde visuals showing BMAp→ENTl(IV-VI) and ENTl(IV-VI)→BMAp connections. Ellipses denote locations of FG and Phal injections that validate those connections as seen in bottom panels. Note Phal labeled fibers and FG labeled cells in BMAp following injections. **I.** Caudal ENTl→LA connections shown by retrograde visual and validated via a Phal injection.

#### Supplementary Figure 9

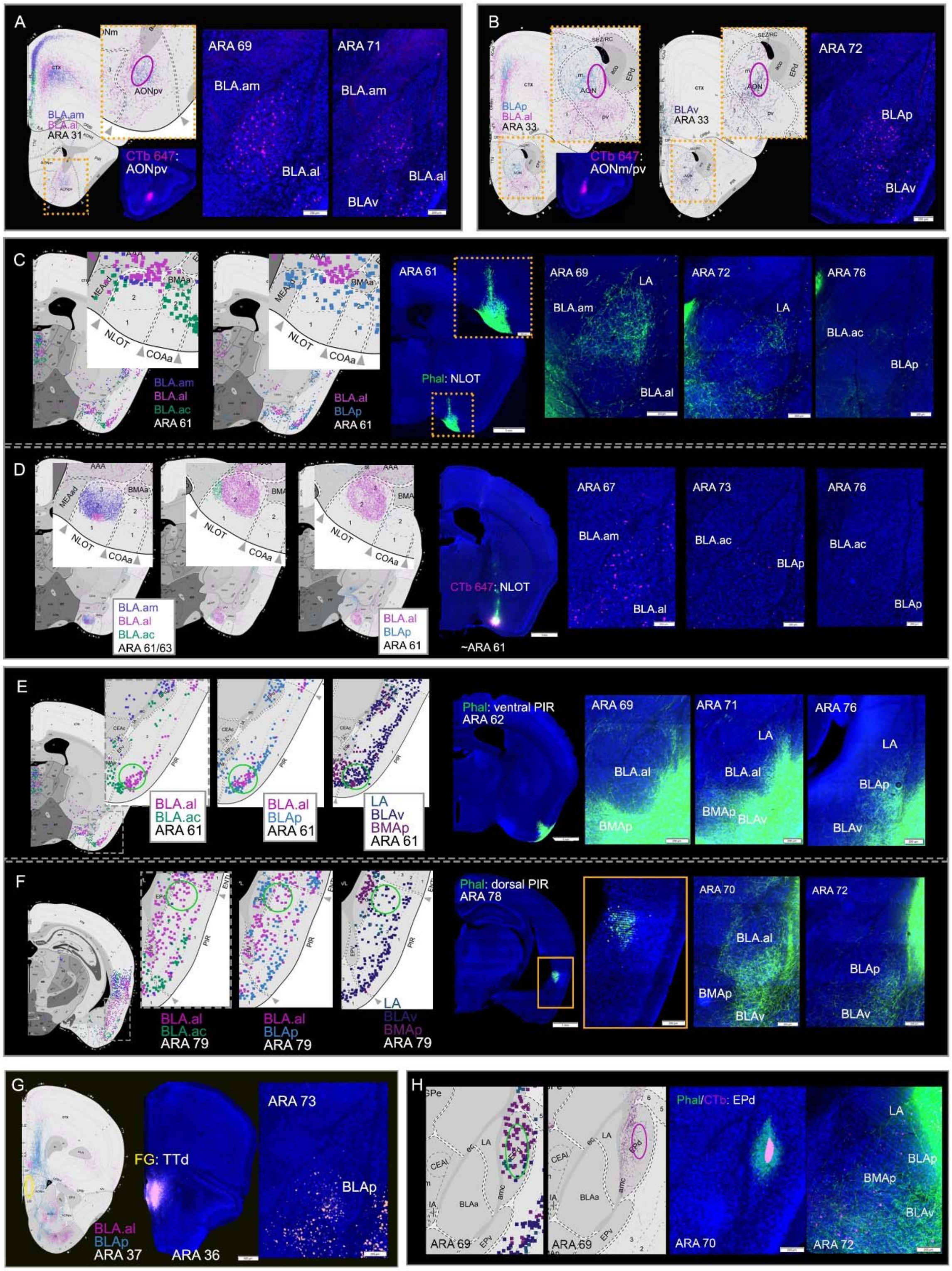

**Supplementary Figure 9**

**A.** Anterograde visuals show BLA.am and BLA.al neuron projections to AONpv (anterior olfactory nucleus, posteroventral part). Inset is magnification of boxed region on visual. Ellipse denotes location of CTb 647 injection that validates BLA.am/BLA.al→AONpv connection, but also back-labels BLAv projection neurons, showing the BLAv→AONpv connection. **B.** Anterograde visuals to the left show BLAp and BLAv projection neurons targeting the AONm (anterior olfactory nucleus, medial part). Insets are magnifications of boxed regions on visuals. A retrograde CTb injection in AONm validates the BLAp/BLAv→AONm connection. **C.** Retrograde **v**isuals to the left show labeled NLOT(I/II) (nucleus of lateral olfactory tract) neurons following retrograde tracer injections in BLA.am, BLA.al, and BLAp. A Phal injection in NLOT labels fibers in BLA.am, BLA.al, and LA validating the NLOT→BLA.am/BLA.al/LA connections, but not the NLOT→BLAp connection. Note the absence of labeled fibers in BLAp. Labeled NLOT projection cells may have been due to slight (~5%) leakage into the LA. Inset is magnification of boxed injection site region. **D.** Anterograde visuals show projection neurons in BLA.am and BLA.al targeting the NLOT. Note the absence of BLAp fibers in NLOT in BLA.al versus BLAp comparison. To the right, a CTb retrograde injection in NLOT validates the BLA.am/BLA.al→NLOT connections. Note absence of labeled cells in BLAp and BLA.ac. **E.** Retrograde visuals show labeled ventral PIR (piriform cortical area) cells following retrograde tracer injections in BLA.al, BLAp, BLAv, BMAp, and LA. Ellipses denote location of Phal injection to validate those connections. A Phal injection in ventral PIR confirms the ventral PIR→BLA.al/BLAp/BLAv/BMAp/LA connections. **F.** Left Retrograde visuals show labeled dorsal PIR neurons following retrograde tracer injections in BLA.al and BLAv. Ellipses denote location of Phal injection to validate those connections. A Phal injection in dorsal PIR labels fibers primarily in BLA.al, BLAv, but also in BLAp confirming the BLA.al/BLAv/BLAp→dorsal PIR projections. Inset shows magnified injection site location. **G.** Left: anterograde visual showing BLAp neuron projections to TTd/TTv (TTd: taenia tecta, dorsal part; TTv: taenia tecta, ventral part). Ellipse denotes location of FG injection that validates the BLAp→TTd connection. **H.** Left panels are retrograde and anterograde visuals showing EPd→BMAp and BMAp→EPd connections. *Abbreviations: EPd: dorsal endopiriform nucleus.* Ellipses denote Phal and CTb injection locations to validate those connections. The Phal EPd injection shows labeled fibers in the BMAp, but also in BLAp and BLAv (EPd→BMAp/BLAp/BLAv), while CTb labels cells only in BMAp (BMAp→EPd).

#### Supplementary Figure 10

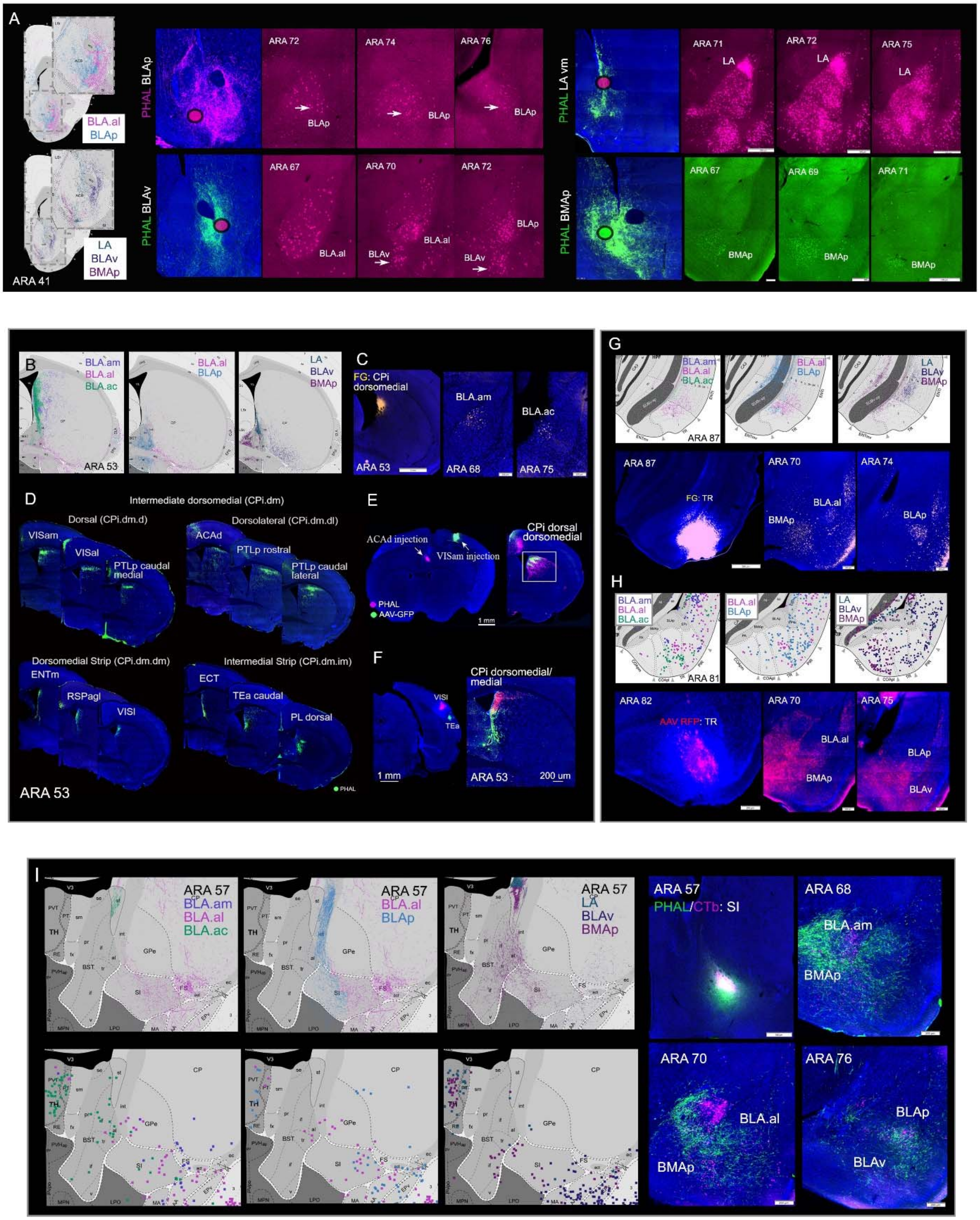

**Supplementary Figure 10**

**A.** Left panels: anterograde visuals showing BLAp and BMAp projections to medial ACB (nucleus accumbens), BLAv to ACB core, and LA to intermediate ACB. Raw images of projections from anterograde BLA injections to each of these ACB regions are shown to the left. Ellipses denote location of retrograde tracer injections to validate each BLA→ACB projection. A CTb 647 (pink) injection in medial ACB back-labels BLAp projection neurons (BLAp→medial ACB), a CTb 647 (pink) injection in ACB core labels BLAv neurons (BLAv→ACB core), a CTb 647 (pink) injection in medial ACB back-labels projection neurons in LA (LA→ACB medial), and a CTb 488 (green) injection in medial ACB back-labels BMAp neurons (BMAp→ACB medial). Note that for each case, retrograde tracers were placed strategically to overlap with anterograde fibers arising from BLAp, BLAv, LA, and BMAp injections. **B.** Anterograde visuals showing BLA.am and BLA.ac projections to dorsomedial part of CP intermediate (CPi) and BLA.al and BLAv projections to ventral CPi. **C.** A FG retrograde injection in dorsomedial CPi back-labels projection neurons in BLA.am and BLA.ac validating those connections (BLA.am/BLA.ac→CPi dorsomedial). **D.** Adapted from Hintiryan et al. (2016), which shows cortical projections to distinct domains in CPi. For example, projections from BLA.am to CPi overlap with fibers from the VISam, VISal, PTLp, and ACAd in dorsomedial regions, while projections from BLA.ac overlap with fibers from ENTm, RSPagl, and VISl in the CPi dorsal and dorsomedial domains. LA fibers overlap with projections from the ECT, TEa caudal, and PL in CPi medial domains. *Abbreviations: VISam: visual cortical area, anteromedial part; VISal: visual cortical area, anterolateral part; PTLp: posterior temporal association area; ACAd: anterior cingulate cortex, dorsal part; ENTm: entorhinal cortical area, medial part; RSPagl: retrosplenial cortical area, agranular part; VISl: visual cortical area, lateral part; ECT: ectorhinal cortical area; TEa: temporal association area; PL: prelimbic cortical area.* **E.** Shows fibers from ACAd and VISam terminating in CPi dorsal and dorsomedial regions, where BLA.am fibers also terminate. **F.** Fibers from VISl and TEa terminate in dorsomedial and medial parts of CPi, precisely where BLA.ac and LA fibers terminate. **G.** Top panels are anterograde visuals showing BLA.al, BLAp, and BMAp projections to TR (piriform-amygdalar transition area). Bottom panels show a TR FG retrograde injection that validates the BLA.al/BLAp/BMAp→TR connections. **G.** Top panels are retrograde visuals showing TR cells back-labeled from retrograde injections in BLA.al, BLAp, BLAv, and BMAp. Bottom panels show AAV RFP anterograde injection that validates TR→BLA.al/BLAp/BLAv/BMAp connections. **I.** Top left panels are anterograde visuals showing projection neurons in BLA.al, BLAp and BMAp targeting SI (substantia innominata). An SI CTb injection validates the BLA.al/BLAp/BMAp→SI connections. Note CTb back-labeled cells in BLA.al, BLAp, and BMAp. Bottom left panels are retrograde visuals showing SI labeled cells from retrograde injections made in BLA.al, BLAv, BMAp, validated with an SI Phal injection that labels the BLA.am, BLAv, BMAp, but also the BLA.am (BLA.am/BLA.al/BLAv/BMAp→SI).

#### Supplementary Figure 11

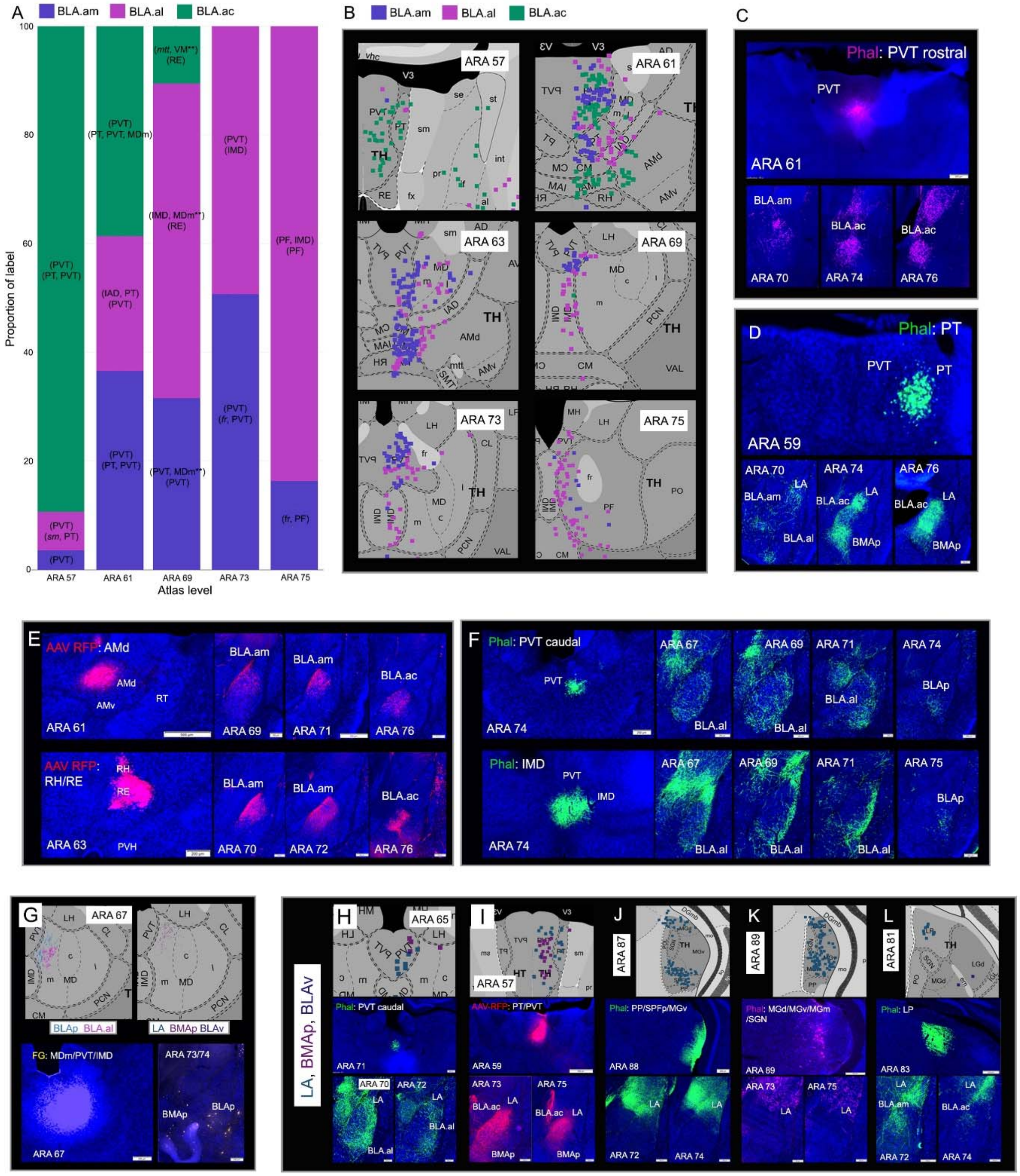

**Supplementary Figure 11**

**A.** Bar chart showing the proportion of labeled neurons in thalamic nuclei at ARA levels 57-75 resulting from single retrograde injections in BLA.am, BLA.al, and BLA.ac (n=1 each). In two separate parentheses, the ROIs for the grids with strongest label from each injection are also included for each ARA section. Note that a grid can include more than one ROI [e.g., (PT, PVT)] and in such cases, the ROIs are presented in order such that those that occupy most of the grid are presented first. In particular, this bar graph shows the topographic projections from projection neurons in more rostral versus caudal thalamic regions. Note the larger proportion of thalamic PVT labeling from BLA.ac injection at rostral levels (ARA 57 and 61) and the larger proportion of thalamic PVT label from BLA.am and BLA.al injections in more caudal levels (ARA 73). As such, this bar graph clearly illustrates the rostral and caudal PVT divide. The bar chart shows strongest thalamic input to BLA.am is from caudal PVT, strongest input to BLA.al comes from caudal PVT, PF, and IMD, and strongest input to BLA.ac from PT and rostral PVT. ** denotes connections that were not validated with anterograde tracing. *Abbreviations: PVT: paraventricular nucleus of the thalamus; PT: parataenial nucleus; MDm: mediodorsal nucleus of thalamus, medial part; IAD: interanterodorsal nucleus of the thalamus; mtt: mammilothalamic tract; RE: reunions nucleus; IMD: intermediodorsal nucleus of thalamus; fr: fasciculus retroflexus; PF: parafascicular nucleus.* **B.** Retrograde visuals from ARA 57-75 showing back-labeled thalamic nuclei from injections in BLA.am, BLA.al, and BLA.ac. **C.** Visuals in B clearly show PVT rostral (ARA 57, 61) and PT neurons target mostly BLA.ac, which are validated with Phal injections in these regions in C (PVT rostral) and **D** (PT). The latter also shows relatively strong PT projections to LA and BMAp (PT→BMAp/LA). **E.** Anterograde tracer injections in AMd and RH/RE validate the AMd→BLA.am/BLA.ac and RH/RE→BLA.am/BLA.ac connections. *Abbreviations: AMd: anteromedial nucleus of the thalamus, dorsal part; RT: reticular nucleus of the thalamus; RH: rhomboid nucleus; RE: reunions nucleus.* **F.** Anterograde tracer injections in PVT caudal and IMD validate the PVT caudal/IMD→BLA.al connections. PVT caudal also shows projections to BLAp (PVT caudal→BLAp). **G.** Top panels: anterograde visuals showing BLAp and BMAp neuron projections to MDm (mediodorsal thalamic nucleus, medial part). Bottom panels: FG injection in MDm validates the MDm→BLAp/BMAp connections. **H.** Top panel: retrograde visual showing PVT caudal cells labeled following a retrograde tracer injection in LA. Bottom panels: Phal tracer in PVT caudal validates the PVTcaudal→LA connection and also PVT caudal→BLA.am/BLA.al proejctions. **I.** Top panel: retrograde visual showing rostral PVT and PT cells labeled following retrograde tracer injections in LA and BMAp. Bottom panels: anterograde AAV-RFP injection validates PVT rostral/PT→LA/BMAp projections, but also the PVT rostral/PT→BLA.ac projection. **J.** Top panel: retrograde visual showing MGd and SPFp labeled cells following retrograde tracer injection in the LA. Bottom panels: Phal injection in PP/SPFp/MGv validates the PP/SPFp/MGv→LA connection. **K.** Top panel: retrograde visual showing MGd/MGv labeled cells following retrograde tracer injection in LA. Bottom panels: Phal injection in MGd/MGv validates MGd/MGv→LA projections. **L.** Top panel: retrograde visual showing PT labeled cells following retrograde tracer injection in the LP. Bottom panels: Phal injection in LP validates the LP→LA connection, but also shows LP→BLA.am connection. *Abbreviations: PP: peripeduncular nucleus; SPFp: subfascicular nucleus, parvicellular part; MGd: medial geniculate complex, dorsal part; MGv: medial geniculate complex, ventral part; MGm: medial geniculate complex, medial part; LP: lateral posterior thalamic nucleus.*

**Supplementary Figure 12**

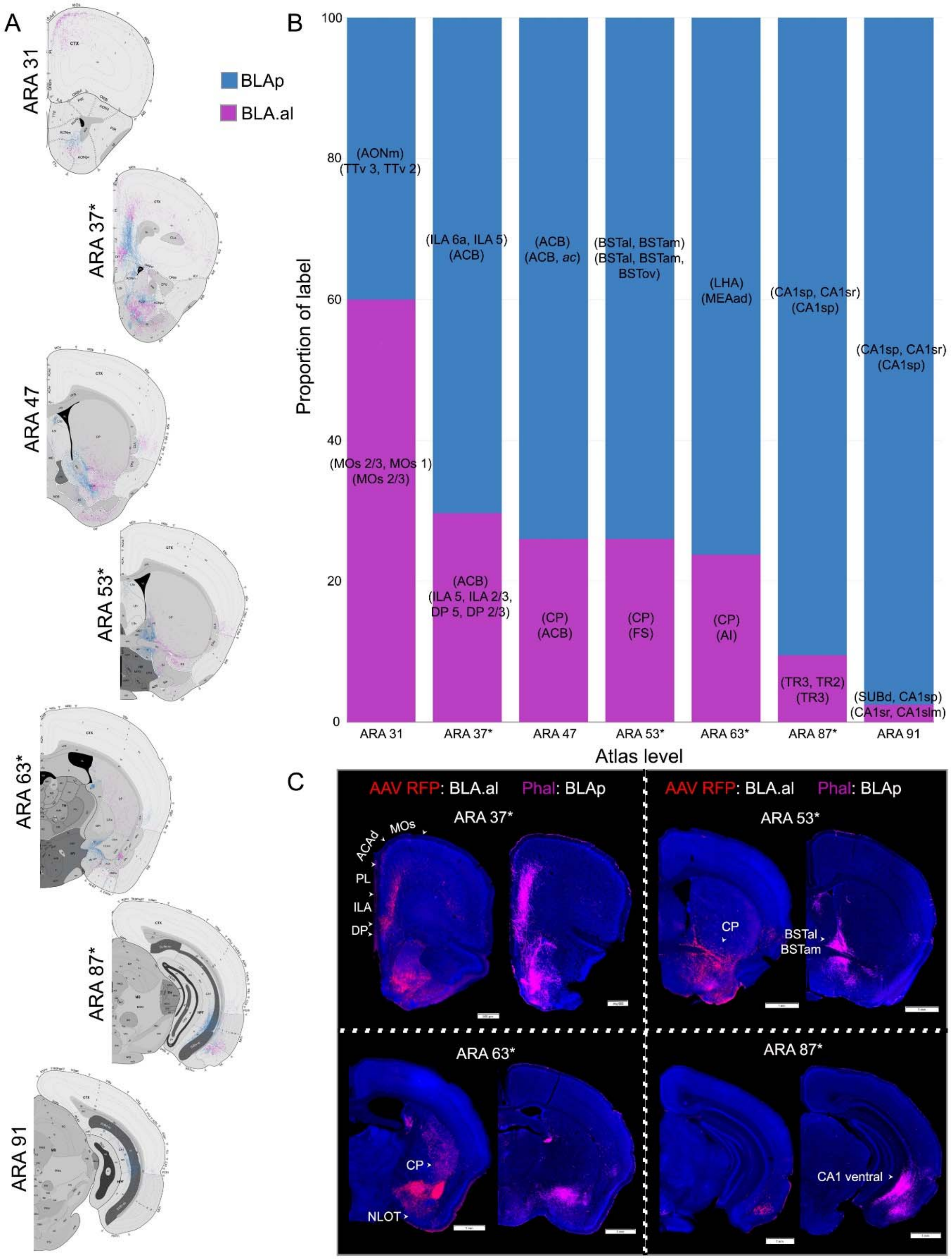

**Supplementary Figure 12**

**A.** Anterograde visual from ARA 31-91 showing the discrepant labeling from BLA.al versus BLAp projection neurons. **B.** Bar chart showing the proportion of anterograde labeling across ARA levels 31-91 resulting from single injections made in BLA.al and BLAp (n=1 each). In two separate parentheses, the ROIs for the grids with strongest label from each injection are also included for each ARA section. Note that a grid can include more than one ROI for example, (BSTal, BSTam) and in such cases, the ROIs are presented in order such that those that occupy most of the grid are presented first. In particular, this bar graph shows the different targets of BLA.al and BLAp neurons. Note for example, at level 53, the primary target of BLAp projection cells are the BST, while primary targets for BLA.al neurons include the CP and FS. At level 87, a primary target for BLAp neurons includes the CA1, while for BLA.al the main target at that level is the TR. Raw data from atlas levels with an * are shown in C. *Abbreviations: BSTam: bed nucleus of stria terminalis, anteromedial part; BSTal: bed nucleus of stria terminalis, anterolateral part; CP: caudoputamen; FS: fundus of striatum; TR: piriform-amygdalar transition area.* **C.** Raw data from atlas levels with an * in B are shown (ARA 37, 53, 63, and 87). Note the difference in ILA and PL labeling at ARA 37. At ARA 53, BLAp neurons target BST, while those in BLA.al target CP, at ARA 63 BLA.al neurons target the CP caudal ventral while there is an absence of label in CP from the BLAp injection. Finally, at level 87 BLAp neurons target CA1, while there is a paucity of label from the BLA.al injection to hippocampal regions.

### Supplementary Figure 13

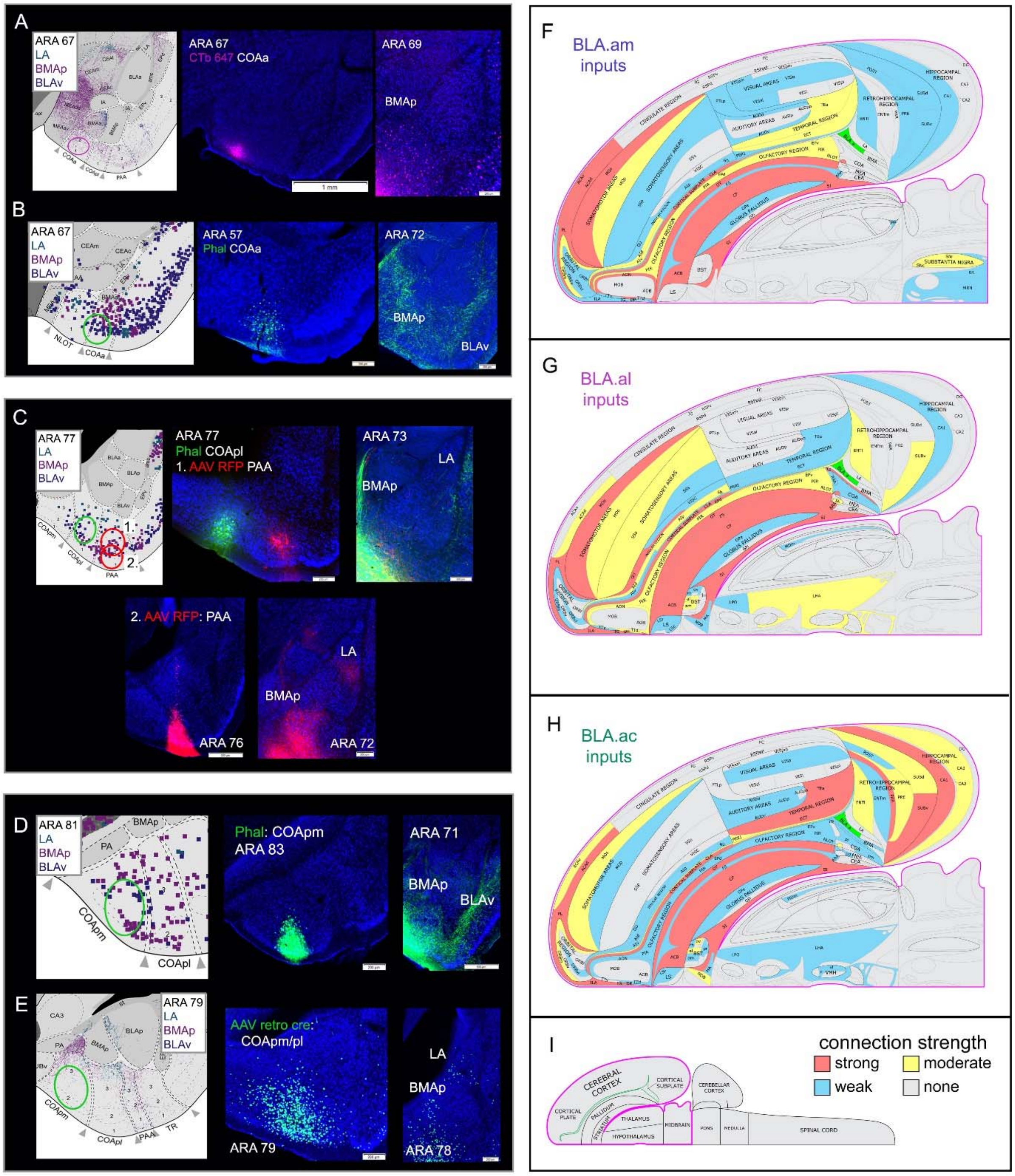

**Supplementary Figure 13**

**A.** Left: anterograde visual showing BMAp neuron projections to COAa (cortical amygdala area, anterior part), a connection validated with a CTb injection in COAa that back-labels BMAp neurons (BMAp→COAa). Ellipse on visual denotes location of COAa CTb injection site. **B.** Retrograde visual shows COAa labeled cells from retrograde tracer injections made in BMAp and BLAv. COAa→BMAp/BLAv connections are validated with a Phal injection in COAa, which labels fibers in BMAp and BLAv. Ellipse on visual denotes location of Phal injection site. **C.** Top panels: retrograde visual shows back-labeled COApl (cortical amygdala area, posterolateral part) and PAA (piriform-amygdala area) cells following retrograde tracer injections in BMAp and LA. These connections are validated with anterograde Phal (green ellipse) and AAV-RFP (1. red ellipse) injections in COApl and PAA, which label fibers in BMAp and LA, confirming COApl/PAA→BMAp/LA connections. Bottom panels: A second AAV-RFP injection in PAA (2. red ellipse) more clearly labels BMAp and LA as further validation of those connections. **D.** Retrograde visual showing COApm (cortical amygdala area, posteromedial part) labeled cells following retrograde tracer injections in BMAp and BLAv. A Phal injection (green ellipse) in COApm labels BMAp and BLAv validating those connections. **E.** Anterograde visual shows BMAp projections to COApm. An AAV retro Cre injection (green ellipse) in COApm back-labels projection cells in BMAp validating BMAp→COApm connection. **F**-**I**. The inputs of BLA.am (**F**), BLA.al (**G**), and BLA.ac (**H**) domains are represented at the macroscale (gray matter region resolution) on a partial flatmap representation of the mouse brain. The strength values of detected connections were binned into tertiles, and these are represented qualitatively as strong (red), moderate (yellow), and weak (blue). For list of abbreviations, see Table 1. **I.** Longitudinal half of the entire CNS flatmap showing orientation and major brain divisions (e.g., cerebral cortex, pallidum, thalamus). The key to the color codes for connection strength is also shown.
