## Supplemental methods for "Connectivity characterization of the mouse basolateral amygdalar complex"

To trace the connectivity of projection defined neurons, a Cre-dependent anterograde tracing strategy was employed, which involved the delivery of Cre to projection neurons of interest via an injection of AAVretro-EF1a-Cre into their target regions. Next, AAV1.CAG.FLEX.eGFP.WPRE.bGH and AAV1.CAG.FLEX.TdTomato.WPRE.bGH (Addgene; plasmid ID #s 51502, 51503) were delivered to the different Cre-expressing BLA neuronal populations. Finally, to retrieve the morphological information from different amygdalar projection neurons, Gdel-RV-4tdTomato and Gdel-RV-4eGFP injections were made into two different downstream targets of the neurons. Cloning of pRVΔG-4tdTomato [3] (Addgene #52500) and pRVΔG-4GFP [4] (Addgene #52487) has been described. Production of B19G-enveloped rabies virus was done as described previously [4-6] but using helper plasmids pCAG-B19N (Addgene #59924), pCAG-B19P (Addgene #59925), pCAG-B19G (Addgene #59921), pCAG-B19L (Addgene #59922), and pCAG-T7pol (Addgene #59926) for the rescue step[6]. The final titers were 2.14e10 infectious units/ml for RVΔG-4tdTomato(B19G) and 1.51e11 infectious units/ml for RVΔG-4GFP(B19G), as determined by infection of HEK 293T cells as described previously [4].

Tissue processing and imaging in 2D: Either one (Phal, CTb, FG, AAV retro Cre, Gdel-RV) or three (Cre- and non-Cre-dependent AAVs) weeks was allowed for tracer transport after which animals were perfused and their brains were extracted. For all but the morphology studies involving Gdel-RV injections, a 2D tissue processing workflow was followed. After an overdose injection of sodium pentobarbital, each animal was transcardially perfused with approximately 50 ml of 0.9% NaCl followed by 50 ml of 4% paraformaldehyde solution (PFA; pH 9.5). The brains were post-fixed in 4% PFA for 24-48 hours at 4°C after which they were embedded in 3% Type I-B agarose (Sigma-Aldrich) prior to sectioning. Four series of coronal sections were sliced at 50-μm thickness with a Compresstome (VF-700, Precisionary Instruments, Greenville, NC) and prepared for immunofluorescence staining.

One series of sections was immunostained for the antigen of interest (Phal or AAVretro-EF1a-Cre) using the free-floating method. Briefly, sections were transferred to a blocking solution containing normal donkey serum (Vector Laboratories) and Triton X-100 (VWR) for 1 hour. Following three 5-minute rinses, sections were incubated in a KPBS solution comprised of donkey serum, Triton, and the appropriate antibody [1:1000 rabbit anti-Phal antibody (Vector Laboratories, #AS-2300)] or 1:4000 mouse anti-Cre recombinase antibody (EMD Millipore, #MAB3120) for 48-72 hours at 4°C. Sections were rinsed 3 times in KPBS and then soaked for 3 hours in the secondary antibody solution, which contained donkey serum, Triton, and a 1:500 concentration of anti-rabbit IgG conjugated with Alexa Fluor® 488 or 647 (Invitrogen, 488: #A-21206; 647: #A-31573) for Phal staining. For Cre recombinase staining, the secondary solution contained donkey serum, Triton, and a 1:500 concentration of anti-mouse IgG conjugated with Alexa Fluor® 488 or 647 (Life Technology, 488: #A-21202; 647: #A-31571). Following 3 KBS rinses, the sections were counterstained with a fluorescent Nissl stain, NeuroTrace® 435/455 (NT; 1:500; Invitrogen, #N21479). The sections were then mounted and coverslipped using 65% glycerol.

Sections were scanned at 10x magnification as high-resolution virtual slide image (VSI) files using an Olympus VS120 high-throughput microscope using identical exposure parameters across all cases.

**Post image acquisition data processing**

Sections from each case were assigned and registered to a standard set of 32 corresponding Allen Reference Atlas levels ranging from 25-103. Variations in tissue cut angle can lead to challenges in atlas level assignments for individual sections. For example, for sections that contains both labeling and an injection site, the fittest corresponding atlas level for the location of tracer labels may differ from the level that best fits the injection sites. As such, these sections were registered twice; once to the atlas level most ideal for the labeling and a second time to the level most fit for the injection sites. In addition, atlas level assignments were made to optimize accurate annotation. Consequently, attention was particularly focused on regions that contained labeling. For example, sometimes due to cut angle discrepancies, the dorsal and ventral parts of a tissue section would ideally get matched and registered to different ARA levels, which technical limitations preclude. For such cases, the atlas levels most ideal to the location of the labels were selected. As such, the annotation would accurately reflect the anatomic location of the labeled cells or fibers.

Threshold parameters were individually adjusted for each case and tracer. Conspicuous artifacts in the threshold output were corrected in Photoshop. For a small proportion of cases, accurate alignment of sections to atlas template was precluded due to limitations of our registration or due to the cut angle of the section. Consequently, label would locate outside of correct ROI. For these cases, position of the label was adjusted to the correct ROI location. Further, oversaturation at the injection site prevents the detection of labeled fibers and boutons at the injection sites (BLAa) and their surrounding areas (LA, CEA). Therefore, with this injection strategy, it is difficult to assess intra-amygdalar connections especially within the BLA complex and the CEA. Due to proximity to the injections sites, these areas appear as dense label in the threshold output subsequently affecting annotation and analysis. To obviate this, these areas of potentially false dense label at or surrounding the injection sites (specifically the BLA complex and CEA) were filtered out at the thresholding stage for all BLAa cases.

| Injection | Structures omitted from analysis |
| --- | --- |

| | |
|---|---|
| BLA.am | LA, BLAa, BLAp, CEA |
| BLA.al | LA, BLAa, BLAp, CEA |
| BLA.ac | LA, BLAa, BLAp, CEA |
| BLAp | LA, BLAa, BLAp, CEA |
| BMAp | BMAp, BLAv, LA |
| BLAv | BMAp, BLAv, LA |
| LA | BMAp, BLAv, LA |

**Data analysis and visualization**

Injection site analysis: Some injection sites were rescanned under lower exposure parameters to acquire a more accurate view of the size and location of the injections. Sections containing the injections for each case were re-registered to ARA levels that best corresponded to the target BLA complex structures. For injections that spanned more than a single section, each section was registered and analyzed. An injection site annotation algorithm (detailed below) was run on the sections to identify injection location. The annotation was run atop a custom ARA atlas that contained the manually delineated boundaries of the BLAa domains.

Injection site annotation: Correct annotation of the spatial location of injection sites is critical in interpreting neuroanatomical data. In both anterograde and retrograde tracing experiments, the injection sites are typically surrounded by irregularly shaped, very high intensity background pixels. These high intensity background regions yield little useful connectivity information and tend to skew the overall connectivity quantification results as well as interfering with injection site annotation. To quantify the injection sites robustly and consistently, we employ a combination of multi-scale wavelet decomposition, non-linear adaptive intensity adjustment and maximally stable external region (MSER) detection.

Wavelet decomposition encodes both frequency and spatial information of the input data by successive application of high and low pass filters. Given a 2D image $f(x, y)$, its discrete wavelet decomposition of level $L$ is as following:

$$f(x,y) = \sum_i \sum_j C_{L,ij}^L \cdot \Phi_{L,ij}(x,y) + \sum_l \sum_i \sum_j C_{l,ij}^H \cdot \Psi_{l,ij}^H(x,y) + \sum_l \sum_i \sum_j C_{l,ij}^V \cdot \Psi_{l,ij}^V(x,y) + \sum_l \sum_i \sum_j C_{l,ij}^D \cdot \Psi_{l,ij}^D(x,y)$$

Where $C_l^L$ is the level $l$ coefficients for the low pass band, and $C_l^H, C_l^V, C_l^D$ are the level $l$ coefficients for horizontal, vertical and diagonal detail bands ($\Phi_l$ and $\Psi_l$ are the scaling and wavelet functions). For a brain section image $f_{inj}$ containing an injection site, we decompose the image into 5 levels and remove the details from levels 1, 2 and 3. At levels 4 and 5, diagonal detail coefficients with large magnitudes are amplified, while horizontal and vertical coefficients with small magnitudes are dampened. Level 5 low pass band coefficients are also dampened and further smoothened with a gaussian kernel. The reverse wavelet transformation of the modified coefficients $\widetilde{C_{L,ij}^{L}}, \widetilde{C_{l,ij}^{H}}, \widetilde{C_{l,ij}^{V}}, \widetilde{C_{l,ij}^{D}}$ yields a reconstruction of the image $\widetilde{f_{inj}}$ with much of the background intensity in the injection site removed:

$$\widetilde{f_{inj}} = \sum_i \sum_j \widetilde{C_{L,ij}^{L}} \cdot \Phi_{L,ij}(x,y) + \sum_l \sum_i \sum_j \widetilde{C_{l,ij}^{H}} \cdot \Psi_{l,ij}^{H}(x,y) + \sum_l \sum_i \sum_j \widetilde{C_{l,ij}^{V}} \cdot \Psi_{l,ij}^{V}(x,y) + \sum_l \sum_i \sum_j \widetilde{C_{l,ij}^{D}} \cdot \Psi_{l,ij}^{D}(x,y)$$

The contrast of the reconstructed image $\widetilde{f_{inj}}$ is enhanced using the local adaptive mapping described in [7]. For a normalized image $f(x,y) \in [0,1]$, an initial intensity mapping $T$ is defined as

$$T(f(x,y),p) = sin^2\left(\frac{\pi}{2} f(x,y)^p\right), \qquad p > 0$$

Using the first order Taylor expansion approximation of $\sin(u)$, the mapping is rewritten as

$$T(f(x,y),c) = \frac{\pi^2}{4} f(x,y)^c, \quad c = 2p$$

The mapping argument $c$ is defined as $c = c_1 \cdot \frac{f_g(x,y)+\epsilon}{(1-f_g(x,y))+\epsilon} + c_2$, where $f_g(x,y) = f(x,y) * g(x,y)$ denotes the convolution between $f(x,y)$ and a gaussian kernel $g(x,y)$, while $c_1, c_2$ are user specified values. Locally adaptive contrast enhancement is achieved as following:

$$E(x,y) = \frac{f(x,y)}{f_g(x,y)} T(f(x,y),c) + \frac{f(x,y)}{f_g(x,y)} \frac{\partial T(f(x,y),c)}{\partial c} \cdot (f(x,y) - f_g(x,y))$$

The contrast enhancement further suppresses background pixels at injection site. Finally, the injection site is extracted as a blob by MSER from $E(\widetilde{f_{inj}}(x,y))$, the result of applying wavelet filtering and adaptive contrast enhancement to the injection site image.

<u>BLAa boundary demarcation:</u> Despite sharing many common input and output pathways, the medial, lateral, and caudal parts of BLAa each has discerningly connections with different regions of the brain. These division specific connections were used to compute the boundaries between BLAa divisions.

We first examined a collection of coronal sections containing BLAa to identify a subset of sections $S$, where division specific connections can be observed. Each section $s \in S$ was then associated with a division label $y_s \in Y = \{medial, lateral, caudal\}$, and registered to the Allen Reference Atlas (ARA). Due to the large z dimension sampling gap of 50 µm between adjacent ARA levels, the BLAa division boundary is computed individually for each ARA level. For any given level $l$, let $S^l$ be the subset of sections in $S$ that best matches with ARA at level $l$. Tracer signal foreground pixels were segmented for each section in $S^l$ to obtain a set of coordinates $D_s^l = \{(x_{s11,}x_{s12}), (x_{s21,}x_{s22}), \ldots\ x_{sn1,}x_{sn2}\}$. All $(x_{si1,}x_{si2})$ pairs inherit the section division label $y_s$. We used $D^l = \bigcup_{i=1}^{card(S^l)} D_s^l$ and its corresponding division labels $\boldsymbol{y^l}$ to train an ensemble of RBF kernel support vector machines (SVM). The ensemble then determines BLAa subdivisions by assigning a division label to all spatial locations within BLAa.

A SVM classifier solves the following optimization problem:

$$\min_{w,b,\zeta} \frac{1}{2} w^T w + C \sum_{i=1}^{n} \zeta_i$$

$$s.t. \quad y_i(w^T \varphi(\boldsymbol{x}_i) + b) \geq 1 - \zeta_i$$

$$\zeta_i \geq 0$$

The classifier has two parameters: the soft margin parameter $C$ and the kernel (radial basis function $\varphi$) parameter $\gamma$. A total of 64 classifiers were trained with $C$ and $\gamma$ take values on a grid of powers of 2. For a classifier $SVM_i$, a model accuracy $a_i$ is computed as the average accuracy from 10 3-fold stratified cross-validation training and testing. Given any point $\boldsymbol{x} = (x_1, x_2)$ within BLAa at level $l$, denote the division prediction by $SVM_i$ as $y_i(\boldsymbol{x})$. The BLAa division of $\boldsymbol{x}$ at level $l$ is determined by a weighted averaging ensemble:

$$E^l(\boldsymbol{x}) = \underset{y \in Y}{\text{argmax}} \sum_i a_i \cdot I(y_i(\boldsymbol{x}) = y)$$

Dense classification across BLAa with $E^l$ produces the optimal BLAa divisions based on division specific anatomical pathways.

<u>Community detection</u>*:* We performed community detection (modularity maximization) on both grid and ROI annotated data. As a first step, the overlap annotation per each group was aggregated into a single matrix. We set a minimum threshold value of 0.0045 for labeled pixels (anterograde) and 8 for labeled cells (retrograde), and removed entries that did not meet the threshold. These minimum values were set to exclude potential false positives, but also to exclude extremely light connections that would be difficult to explain (e.g., 2 labeled cells or a few labeled fibers in an ROI). Once the aggregated matrix was constructed, we further normalized so that the total labelling across each injection site (typically close in the first place) was adjusted to equal with the injection site featuring maximum total labeling. On this normalized matrix we applied the Louvain [8] algorithm at a single scale (gamma 1.0). As the result of this greedy algorithm is non-deterministic, we performed 100 separate executions, and subsequently calculated a consensus community structure [9] to characterize the 100 executions as a single result.

Connectivity matrices, community color coding and data visualization: For matrix visualization, we applied the grid_communities algorithm (bctpy; https://github.com/aestrivex/bctpy, https://sites.google.com/site/bctnet/) to modularize ROIs based on community assignment, as well as prioritize connections along the diagonal. Such a visualization provides a high level overview of the connectivity (Figure 3H-I; Figure 4F-G).

Community coloring provides more detail (i.e. drill down from the matrix overview) by visually encoding segmentation by community, and ultimately, injection site (Figure 1A, anterograde and retrograde visuals). To carry out this process, we developed software to programmatically march through each segmented pixel per each image. Using either the grid location or ROI name, the algorithm looked up the corresponding community assigned during the consensus community step. Using a table containing a color assigned (by the authors) to each injection site, the algorithm retrieved the injection site associated with the community, and colored the pixel with the corresponding injection site color value. A subsequent step took advantage of the fact that each pixel was assigned to only a single community by aggregating all colorized images corresponding to a given atlas level into a single representative image.

Proportional stacked bar charts (with and without focus): It is challenging to represent grid-based overlap connectivity in matrix form. A naive visualization labels row and columns by grid location, which is hardly informative. One solution is to use color coding to identify community labels [1]. But comprehending that visualization requires comparisons with at least one other image to qualify the anatomical region that the labeling occurs in. Such a matrix also does not identify where the labelling occurs along the rostral to caudal axis, for example.

To overcome these limitations, we visualized the connectivity data using a proportional, stacked bar chart approach. The chart categorizes data by atlas level. The community coloring is retained, but region labels identify the two ROIs that the labelling occurs most within. A focused version of this visualization isolates the analysis to a specific set of ROIs. Another view splits the ROIs into quadrants (and other combinations of medial-lateral, dorsal-ventral) to resolve termination at higher resolution.

**3D workflow for assessment of neuronal morphology**

To assess whether projection neurons located in the 3 domains of the BLAa were morphologically distinct, representative neurons in each of the domains were labeled via a G-deleted rabies injection in the caudal dorsomedial caudoputamen (for BLA.am), caudal ventral caudoputamen (for BLA.al), and in the medial accumbens (for BLA.ac) (Figure 1D). One week was allowed for tracer transport following injections, after which the animals were perfused. A 3D tissue processing workflow was followed for implementation of the SHIELD clearing protocol [10, 11].

3D tissue processing: Mice were transcardially perfused with ice-cold saline and SHIELD perfusion solution. The brains were extracted and incubated in the SHIELD perfusion solution at 4°C for 48 hours. The SHIELD perfusion solution was replaced with the SHIELD OFF solution and tissues were incubated at 4°C for 24 hours. The SHIELD OFF solution was replaced with the SHIELD ON solution and the tissues were incubated at 37°C for 24 hours. The whole brain was cut into 250- (for BLA.am and BLA.al) or 400 (for BLA.ac) µm sections and were cleared in the SDS buffer at 37°C for 72 hours. The sections were then washed three times with KPBS and incubated in KPBS at 4°C for 24 hours.

3D imaging protocol: Sections were mounted and coverslipped onto 25x75x1mm glass slides with an index matching solution 100% (EasyIndex, LifeCanvas Technologies, #EI-Z1001). Sections were imaged with a high speed spinning disk confocal microscope (Andor Dragonfly 202 Imaging System, Andor an Oxford Instruments Company, CR-DFLY-202-2540). 10x magnification (NA 0.40, Olympus, UPLXAPO10X) was used to acquire an overview after which 30x magnification (NA 1.05, Olympus, UPLSAPO30xSIR) was used to image through the BLA ipsilateral to the injection site at 1 µm z steps. The BLA contralateral to the injection was also imaged for the BLA.ac case. Sections were imaged with four excitation wavelengths (nm): 405 (blue Nissl background), 488 (green for rabies), 561 (red for rabies) with respective emission detection wavelengths of 450, 525, and 600.

Statistical analysis: To obtain an overall view of the dendritic morphology of BLA projection neurons located in the BLA.am, BLA.al, and BLA.ac, we applied the classic [14] and modified Sholl [15] analysis using Fiji ImageJ and L-Measure, respectively. The resulting classic Sholl scatter plot shows dissimilarity between groups, especially within 100-200 nm radius from the cell body (Supplementary Figure 1I). A three-dimensional Sholl-like analysis showed that in BLA.ac neurons the distribution of dendritic surface area was more peaked towards the median of the relative path distance from the soma as opposed to the more uniformly distributed dendritic surface of BLA.am and BLA.al neurons (Figure 8A). A visual comparison shows that the BLA.ac neurons are qualitatively more distinct from those in the BLA.am and BLA.al.

Quantitative morphological parameters depicting the somas and dendrites were obtained from Aivia and statistical analyses were performed using the R computing environment (RStudio Version 1.1.463). Standard measurements representing cell body morphology included volume, sphericity, spherical diameter, X, Y, Z and Euclidean skewness, cell body height, width, depth, and surface to volume ratio. Measurements of dendritic morphology included number of primary dendrites, branches, bifurcations, terminal tips, nodes, fractal dimension, tortuosity, local bifurcation angle, remote bifurcation angle, partition asymmetry, tips to dendrite ratio, branch generation, and Rall's ratio.

Overall, the results suggest that neurons located in each of the BLAa domains are morphologically distinct. Using all of the measured morphological parameters principal component analysis (PCA) was run to reduce the dimensionality and create a 3D scatterplot [16, 17]. The PCA shows the segregation of BLAa domain specific neurons based on the measured features (Figure 8B).

Two-sided pairwise Wilcoxon rank sum tests were performed, and the parameters that survived FDR correction for multiple testing with a q-value <0.05 are reported. The significant group differences are presented with whisker plots (Figure 8F), and the degree of their significance is visualized in a matrix plot (Figure 8E). Although significant differences in several morphological features were detected across all pairwise comparisons, generally, neurons in the BLA.am and BLA.al significantly differed from those in the BLA.ac on a greater number of morphological features. This is also evidenced by the dendrogram on top of the matrix that shows the hierarchical clustering based on group similarity (Figure 8E).

Specifically, BLA.am neurons differed from BLA.al neurons in somatic features like cell body height (W=59.5, p=0.04, [0.22, 4.18]) and in dendritic morphology like fractal dimension (W=69, p=0.001, [0.006, 0.017]) and tortuosity (W=67, p=0.0023, [0.02, 0.07]).

Neurons in the BLA.am and BLA.al significantly differed from those in the BLA.ac on cell body features like cell width (BLA.am vs BLA.ac, W=43, p=0.038, [1.76, 8.36]; BLA.al vs. BLA.ac, W=46. p=0.043. [0.44, 5.94]), skewness along the X dimension (BLA.am vs BLA.ac, W=44, p=0.012, [6.5, 32.0]; BLA.al vs. BLA.ac, W=49, p=0.012, [6.5,33.5]), and sphericity (BLA.am vs BLA.ac, W=47, p=0.004, [0.01, 0.036]; BLA.al vs. BLA.ac, W=50, p=0.0072, [0.012, 0.038). They also differed in a number of dendritic features like the number of bifurcations (BLA.am vs. BLA.ac, W=6.5, p=0.041, [-12.0 -0.99]; BLA.al vs. BLA.ac, W=4.5, p=0.027, [-10.0, -1.99]), number of branch generations (BLA.am vs. BLA.ac, W=2, p=0.004, [-0.95, -0.19]; BLA.al vs BLA.ac, W=1, p=0.004, [-0.1, -0.35]), and tips to dendrite ratio (BLA.am vs. BLA.ac, W=4, p=0.018, [-1.86, -0.4]; BLA.al vs BLA.ac, W=2.5, p=0.014, [-1.8, -0.47]).

BLA.am neurons are differed from those in the BLA.ac on additional features including cell height (W=46, p=0.016, [1.98, 5.72]), fractal dimension (W=48, p=0.001, [0.009, 0.02]), tortuosity (W=48, p=0.002, [0.02, 0.08]), soma volume (W=45, p=0.014, [1136.7, 3795.5]), surface to volume ratio (W=1, p=0.004, [-0.06, -0.001]), and spherical diameter (W=45, p=0.014, [1.37, 4.1]).

Domain-specific BLAa neurons did not significantly differ from one another in the number of primary dendrites, number of nodes, number of branches, terminal tips, local bifurcation angle, remote bifurcation angle, partition asymmetry, Rall's ratio, depth, Y, Z, or Euclidean skewness.

Finally, differences between neurons in the contralateral and ipsilateral BLA.ac were not detected for any morphological feature.

In a final analysis, we characterized our data using an alternative tool that uses topological data analysis that retains potentially more morphological information. Specifically, we used the persistence-based neuronal feature vectorization framework [18] to summarize pairwise distances between neurons [19]. Our experiments used code provided by the authors online, where we first computed persistence diagrams using NeuronTools (https://github.com/Nevermore520/NeuronTools) and then computed inter-neuron distances using the Wasserstein metric (https://bitbucket.org/grey_narn/geom_matching/src). The results were visualized in matrix plot created using R (Figure 8G-H). Similar to the PCA analysis, the data showed that groups of projections neurons in BLA.am, BLA.al, and BLA.ac all differed from one another, but the largest differences were between BLA.am/BLA.al neurons compared to those in BLA.ac (Figure 8H).

**Slice recordings**

Three weeks following the viral injections of AAV-hSyn-ChR2-YFP (UPenn Vector Core) and red retrograde microbeads (Lumafluor Inc.), animals were decapitated following isoflurane anesthesia and the brain was rapidly removed and immersed in an ice-cold dissection buffer (composition: 60 mM NaCl, 3 mM KCl, 1.25 mM NaH2PO4, 25 mM NaHCO3, 115 mM sucrose, 10 mM glucose, 7 mM MgCl2, 0.5 mM CaCl2; saturated with 95% O2 and 5% CO2; pH = 7.4). Coronal slices at 350 μm thickness were sectioned by a vibrating microtome (Leica VT1000s), and recovered for 30 min in a submersion chamber filled with warmed (35°C) ACSF (composition:119 mM NaCl, 26.2 mM NaHCO3, 11 mM glucose, 2.5 mM KCl, 2 mM CaCl2, 2 mM MgCl2, and 1.2 NaH2PO4, 2 mM Sodium Pyruvate, 0.5 mM VC). BLA neurons (labeled with fluorescent microbeads) surrounded by EYFP+ fibers were visualized under a fluorescence microscope (Olympus BX51 WI). Patch pipettes (~4–5 MΩ resistance) filled with a cesium-based internal solution (composition: 125 mM cesium gluconate, 5 mM TEA-Cl, 2 mM NaCl, 2 mM CsCl, 10 mM HEPES, 10 mM EGTA, 4 mM ATP, 0.3 mM GTP, and 10 mM phosphocreatine; pH = 7.25; 290 mOsm) were used for whole-cell recordings. Signals were recorded with an Axopatch 700B amplifier (Molecular Devices) under voltage clamp mode at a holding voltage of –70 mV for excitatory currents, filtered at 2 kHz and sampled at 10 kHz. Tetrodotoxin (TTX, 1 μM) and 4-aminopyridine (4-AP, 1 mM) were added to the external solution for recording monosynaptic responses to blue light stimulation (5 ms pulse, 3 mW power, 10–30 trials).

### References


1. Hintiryan, H.*, et al.* The mouse cortico-striatal projectome. *Nature neuroscience* **19**, 1100-1114 (2016).
2. Zingg, B.*, et al.* Neural networks of the mouse neocortex. *Cell* **156**, 1096-1111 (2014).
3. Chatterjee, S.*, et al.* Nontoxic, double-deletion-mutant rabies viral vectors for retrograde targeting of projection neurons. *Nature neuroscience* **21**, 638 (2018).
4. Wickersham, I.R., Sullivan, H.A. & Seung, H.S. Production of glycoprotein-deleted rabies viruses for monosynaptic tracing and high-level gene expression in neurons. *Nature protocols* **5**, 595-606 (2010).
5. Wickersham, I.R. & Sullivan, H.A. Rabies viral vectors for monosynaptic tracing and targeted transgene expression in neurons. *Cold Spring Harb Protoc* **2015**, 375-385 (2015).
6. Chatterjee, S.*, et al.* Nontoxic, double-deletion-mutant rabies viral vectors for retrograde targeting of projection neurons. *Nat Neurosci* **21**, 638-646 (2018).
7. Zhou, Z., Sang, N. & Hu, X. A parallel nonlinear adaptive enhancement algorithm for low-or high-intensity color images. *EURASIP Journal on Advances in Signal Processing* **2014**, 70 (2014).
8. Blondel, V.D., Guillaume, J.L., Lambiotte, R. & Lefebvre, E. Fast unfolding of communities in large networks. *Journal of Statistical Mechanics: Theory and Experiment* **2008**, P10008 (2008).
9. Lancichinetti, A. & Fortunato, S. Consensus clustering in complex networks. *Scientific reports* **2**, 336 (2012).
10. Kim, S.Y.*, et al.* Stochastic electrotransport selectively enhances the transport of highly electromobile molecules. *Proceedings of the National Academy of Sciences of the United States of America* **112**, E6274-6283 (2015).

11. Park, Y.G., *et al.* Protection of tissue physicochemical properties using polyfunctional crosslinkers. *Nature biotechnology* (2018).
12. Cabeen, R., Laidlaw, D. & Toga, A. Quantitative imaging toolkit: Software for interactive 3D visualization, data exploration, and computational analysis of neuroimaging datasets. in *Proceedings of the International Society for Magnetic Resonance in Medicine (ISMRM)* (2018).
13. Cleveland, W.S. & Devlin, S.J. Locally weighted regression: an approach to regression analysis by local fitting. *Journal of the American statistical association* **83**, 596-610 (1988).
14. Kroner, A., *et al.* TNF and increased intracellular iron alter macrophage polarization to a detrimental M1 phenotype in the injured spinal cord. *Neuron* **83**, 1098-1116 (2014).
15. Scorcioni, R., Polavaram, S. & Ascoli, G.A. L-Measure: a web-accessible tool for the analysis, comparison and search of digital reconstructions of neuronal morphologies. *Nature protocols* **3**, 866 (2008).
16. Tsiola, A., Hamzei-Sichani, F., Peterlin, Z. & Yuste, R. Quantitative morphologic classification of layer 5 neurons from mouse primary visual cortex. *Journal of Comparative Neurology* **461**, 415-428 (2003).
17. Vasques, X., Vanel, L., Villette, G. & Cif, L. Morphological neuron classification using machine learning. *Frontiers in neuroanatomy* **10**, 102 (2016).
18. Li, Y., Wang, D., Ascoli, G.A., Mitra, P. & Wang, Y. Metrics for comparing neuronal tree shapes based on persistent homology. *PloS one* **12**, e0182184 (2017).
19. Kerber, M., Morozov, D. & Nigmetov, A. Geometry helps to compare persistence diagrams. *Journal of Experimental Algorithmics (JEA)* **22**, 1.4 (2017).
